# Motor learning leverages coordinated low-frequency cortico-basal ganglia activity to optimize motor preparation in humans with Parkinson’s Disease

**DOI:** 10.1101/2024.09.13.612783

**Authors:** Kara N. Presbrey, Thomas A. Wozny, Kenneth H. Louie, Simon Little, Philip A. Starr, Reza Abbasi-Asl, Doris D. Wang

**Author notes:** **Corresponding author:** Kara Presbrey. Co-Senior Authors (Reza Abbasi-Asl, Doris D. Wang).

## Abstract

Learning dexterous motor sequences is crucial to autonomy and quality of life but can be altered in Parkinson’s Disease (PD). Learning involves optimizing pre-movement planning (preplanning) of multiple sequence elements to reduce computational overhead during active movement. However, it is unclear which brain regions mediate preplanning or how this process evolves with learning. Recording cortico-basal ganglia field potentials during a multi-day typing task in four individuals with PD, we found evidence for network-wide multi-element preplanning that improved with learning, facilitated by functional connectivity. In both cortex and basal ganglia, pre-movement gamma (γ, 30–250 Hz) activity, historically linked to population spiking, distinguished between future action sequences and became increasingly predictive with learning. For motor cortex γ, this increase was tied to learning-related cross-frequency coupling led by cortically-driven network delta (δ, 0.5–4 Hz) synchrony. More generally, coordinated network δ supported a complex pattern of learning-driven cross-frequency couplings within and between cortex and basal ganglia, including striatal lead of cortical beta (β, 12–30 Hz) activity, reflecting the specialized roles of these brain regions in motor preparation. In contrast, impaired learning was characterized by practice-driven decreases in γ’s predictive value, limited cross-frequency coupling and absent network δ synchrony, with network dynamics possibly altered by pathologically high inter-basal ganglia δ synchrony. These results suggest that cortically-led δ phase coordination optimized cortico-basal ganglia multi-element preplanning through enhanced recruitment of higher-frequency neural activity. Neurostimulation that enhances cortico-basal ganglia δ synchrony may thus hold potential for improving skilled fine motor control in PD.

## INTRODUCTION

Fine motor control is a fundamental aspect of human motor function. Skilled hand movements often require learning a sequence of finger movements, and proficiency is vital to maintaining autonomy. In Parkinson’s Disease (PD), progressive decline in fine motor sequence learning and control, not solely attributable to hallmark motor symptoms, detrimentally impacts quality of life, with needs unmet by conventional deep brain stimulation (DBS) and dopamine replacement therapy^1–16^. This decline may be linked to dysfunction in motor preparation, so closed-loop DBS targeting pathological variations in preparatory neural activity could remediate symptoms^17–20^. However, the learning-dependent neural dynamics of fine motor sequence initiation are poorly understood.

Before the onset of rapid fine motor sequences like typing, humans can plan multiple sequence elements (multi-element preplanning), and learning involves the optimization of this process^19–23^. For sequences composed of at least one differing element within the first few elements, this predicts sequence-specific pre-movement neural activity that is optimized with learning. Indeed, neurophysiology studies in rodents and nonhuman primates suggest that multi-element preplanning is facilitated by the sequence-specific serial activation of neurons in motor cortical and basal ganglia (BG) ensembles, with motor improvement partly driven by increased consistency of ensemble spiking patterns^24–32^. However, in humans, it is unknown which brain regions have sequence-specific pre-movement neural dynamics or how sequence-specific activity changes with learning.

The neural processes that promote consistent ensemble firing patterns with learning also remain unclear, but recent investigation highlights the potential role of pre-movement oscillatory network dynamics. Human studies suggest that network-wide beta (β) desynchronization enables an increase in motor cortical excitability—reflected by a shift to the excitatory phase of motor cortical delta (δ)—which facilitates the activation of motor cortical ensembles to initiate movement^24,29,33–51^. Work in animal models suggests that learning-driven corticostriatal δ synchrony enhances δ-ensemble spike coupling in striatum and motor cortex, resulting in the consistent ensemble firing patterns associated with motor improvement^24,29,52^. Learning-driven changes in β’s influence of cortical excitability could also support motor improvement. However, these proposed network interactions have not been tested with cortico-basal ganglia electrophysiology and directed connectivity analysis in humans or animal models.

We postulated that motor cortex and basal ganglia regions all support multi-element preplanning in PD, which network activity optimizes with successful learning. To test this, we evaluated the learning-dependent preparatory motor control network dynamics in four individuals with PD. We recorded cortico-basal ganglia field potentials while subjects performed a multi-day, multi-sequence typing task (**Figure 1A**). We hypothesized that β→δ→spike interactions influence motor cortex regardless of learning stage but that, with practice, coordinated cortico-basal ganglia δ activity increases the consistency of sequence-specific cortical and basal ganglia spiking patterns through δ→spike coupling. Field potential gamma (γ) activity correlates with neural population firing^53^. Thus, γ activity could reflect temporal patterns and variability in ensemble activity. This anticipates specific β→δ→γ interactions, as well as sequence-specific motor cortex and basal ganglia γ activity that is increasingly predictive of future action sequences with learning (**Figure 1B**). We tested these predictions using single-trial classification of neural activity and directed connectivity analysis.

**Figure 1.**
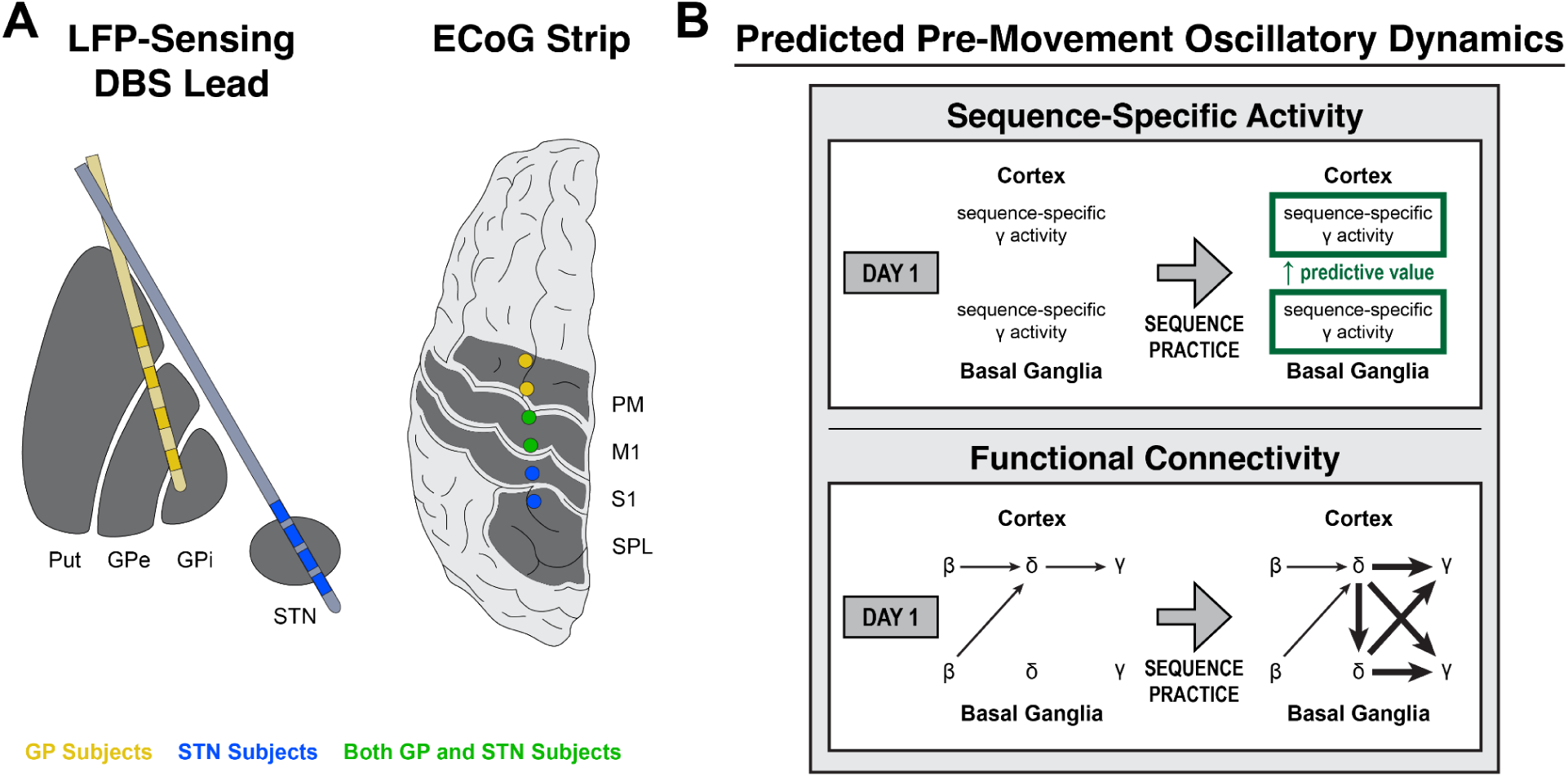
Predicted learning-related cortico-basal ganglia activity during motor sequence initiation. (**A**) Illustration of lead targeting for subject groups (LFP, local field potential; DBS, deep brain stimulation; ECoG, electrocorticography; Put, putamen; GPe, globus pallidus externus; GPi, globus pallidus internus; STN, subthalamic nucleus; PM, premotor cortex; M1, primary motor cortex; S1, primary somatosensory cortex; SPL, superior parietal lobule). (**B**) Diagram of predicted learning-related changes in sequence-specific activity and functional connectivity prior to the onset of motor sequences over multiple days of practice. (Top) For rapid, sequential finger movements, learning involves the optimization of preplanning for multiple sequence elements (multi-element preplanning), potentially implemented by increased reliability of sequence-specific ensemble firing patterns in cortex and basal ganglia. As γ activity correlates with population spiking, this could be reflected by sequence-specific γ activity that becomes increasingly predictive of future action sequences with practice. (Bottom) Oscillatory network dynamics are thought to drive general motor initiation and may display learning-dependent changes that lead to the increased reliability of ensemble activity patterns associated with motor improvement. One possibility is that network β desynchronization enables increased motor cortical excitability, reflected as a shift to the excitatory phase of cortical δ. In turn, excitability facilitates activation of motor cortical ensembles to produce movement, reflected by increasing γ amplitude. With motor learning, increased cortico-basal ganglia δ synchrony facilitates enhanced ensemble recruitment in cortex and basal ganglia, reflected by δ-γ coupling. We thus predicted the presence of network β→cortical δ→cortical γ interactions on all days and that, with practice, increased cortico-basal ganglia δ synchrony would accompany δ-γ coupling in cortex and basal ganglia.

## METHODS

### Study criteria

Four individuals enrolled in parent clinical trials (NCT03582891 and NCT04675398) for adaptive deep brain stimulation (DBS) for Parkinson’s Disease (PD) participated in this study (**Figure 2**, **Table 1**). Subjects had sufficiently severe movement disorder symptoms, inadequately treated by oral medication, and requested surgical intervention. No subjects exhibited significant untreated depression, significant cognitive impairment, previous cranial surgery, drug or alcohol abuse, or evidence of a psychogenic movement disorder. For an exhaustive list of overarching clinical trial inclusion and exclusion criteria, see NCT03582891 and NCT04675398. Additional prescreening was performed for the typing task. Inclusion criterion: enthusiastic desire to participate in the task. Exclusion criteria: hand or wrist pain when typing, dyslexia, uncorrected visual impairment, sleep apnea, travel to other time zones in the past three months. Subjects were also instructed not to consume nicotine or alcohol for the duration of the experiment. Subjects gave informed consent, and the University of California San Francisco Institutional Review Board pre-approved experimental design.

**Figure 2.**
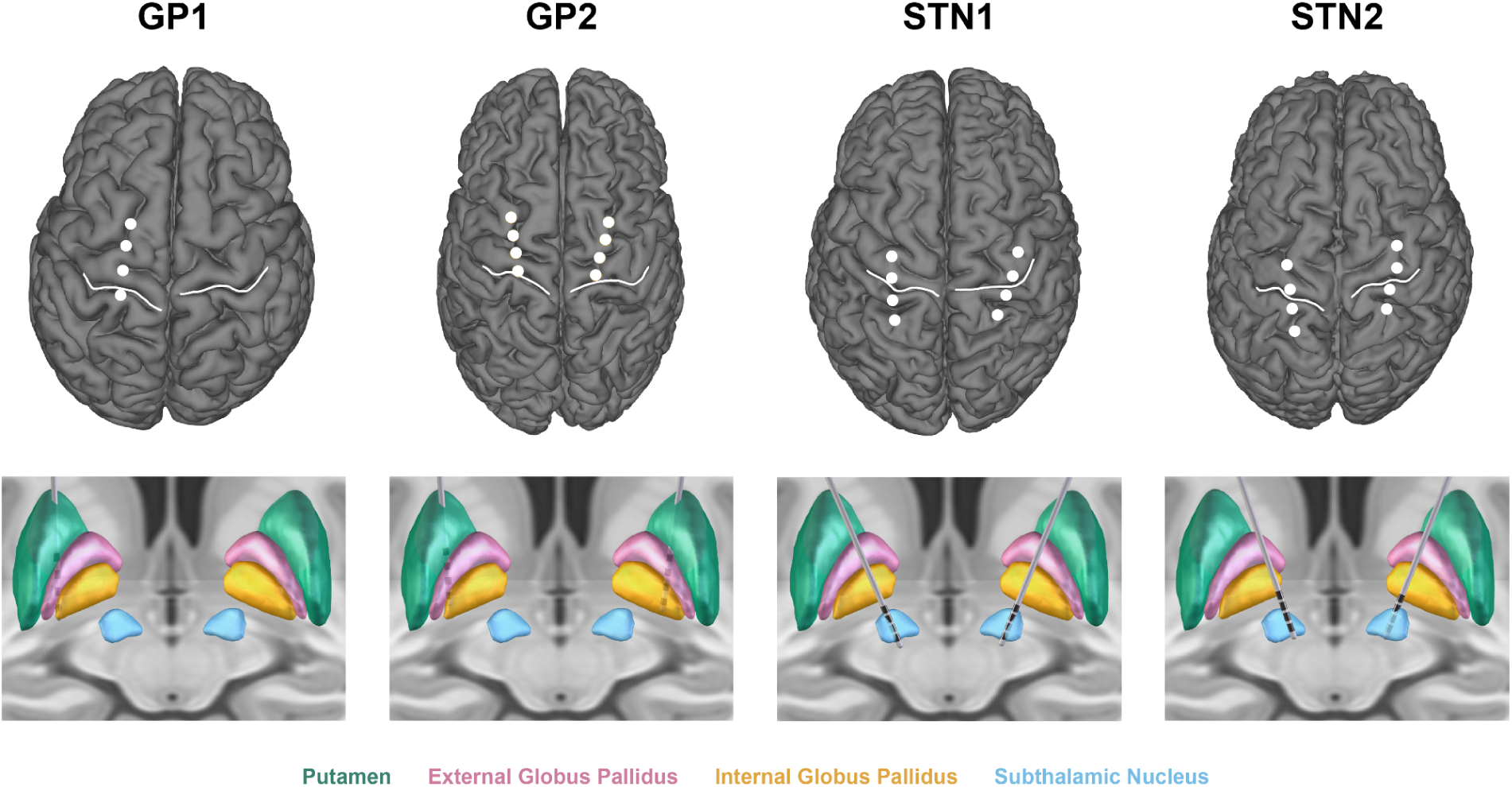
Cortical and subcortical lead reconstructions. (Top) Sensorimotor quadripolar electrocorticography strips, central sulcus (white), and (Bottom) quadripolar deep brain stimulation leads localized within the basal ganglia.

**Table 1.** Subject demographic and clinical information. MoCA, Montreal Cognitive Assessment; UPDRS, Unified Parkinson’s Disease Rating Scale. *Given the small sample size, age and sex have been omitted to retain participant privacy*.

|  | Dominant Hand | Pre-Op MoCA | Pre-Op UPDRS ON Med | Pre-Op UPDRS OFF Med | % Change ON Med | Primary Symptoms |
| --- | --- | --- | --- | --- | --- | --- |
| GP1 | R | 28 | 9 | 18 | -50% | Hand tremor, bradykinesia, gait |
| GP2 | R | 27 | 17 | 31 | -45% | Hand tremor, bradykinesia, gait |
| STN1 | R | 26 | 5 | 32 | -84% | Hand tremor, bradykinesia |
| STN2 | R | 28 | 10 | 31 | -68% | Hand tremor, bradykinesia |

### Task design

On each day of a multi-day explicit motor learning experiment, subjects practiced typing two 5-element sequences in interleaved blocks using their dominant (right) hand while neural activity was recorded from the contralateral hemisphere (**Figure 3A**). The task design within each day was a variation of the common discrete sequence production task^54^. At the start of each session, subjects memorized that day’s sequences during an initial Verification Period. In this Verification Period, they were briefly shown one sequence to memorize before repeatedly typing it from memory until achieving three consecutive fully correct repetitions. This was repeated with the second sequence. Subjects were then instructed to, in the subsequent training blocks, react as quickly as possible and type as quickly and accurately as possible. In each training block, they practiced only one sequence. They typed one sequence repetition from memory in response to each cue. Each practice block started with its own Verification Period for the sequence for that block. The sequence was not shown again in that block. Green go cues appeared after an exponentially jittered delay from the 5th keypress of the previous trial (range: 0.85s–3.75s, μ = 1.75, p = 0.4 for Lilliefors test for h0 = exponential). A 10-second break followed each block. Subjects’ hands and the keypad were completely visually occluded, and the sequences were never displayed during typing. At the end of the task each day, subjects were assessed on the upper limb component of the Movement Disorder Society Unified Parkinson’s Disease Rating Scale (MDS-UPDRS). The day before the experiment, subjects were familiarized with the task and keypad with a practice run-through (Familiarization).

**Figure 3.**
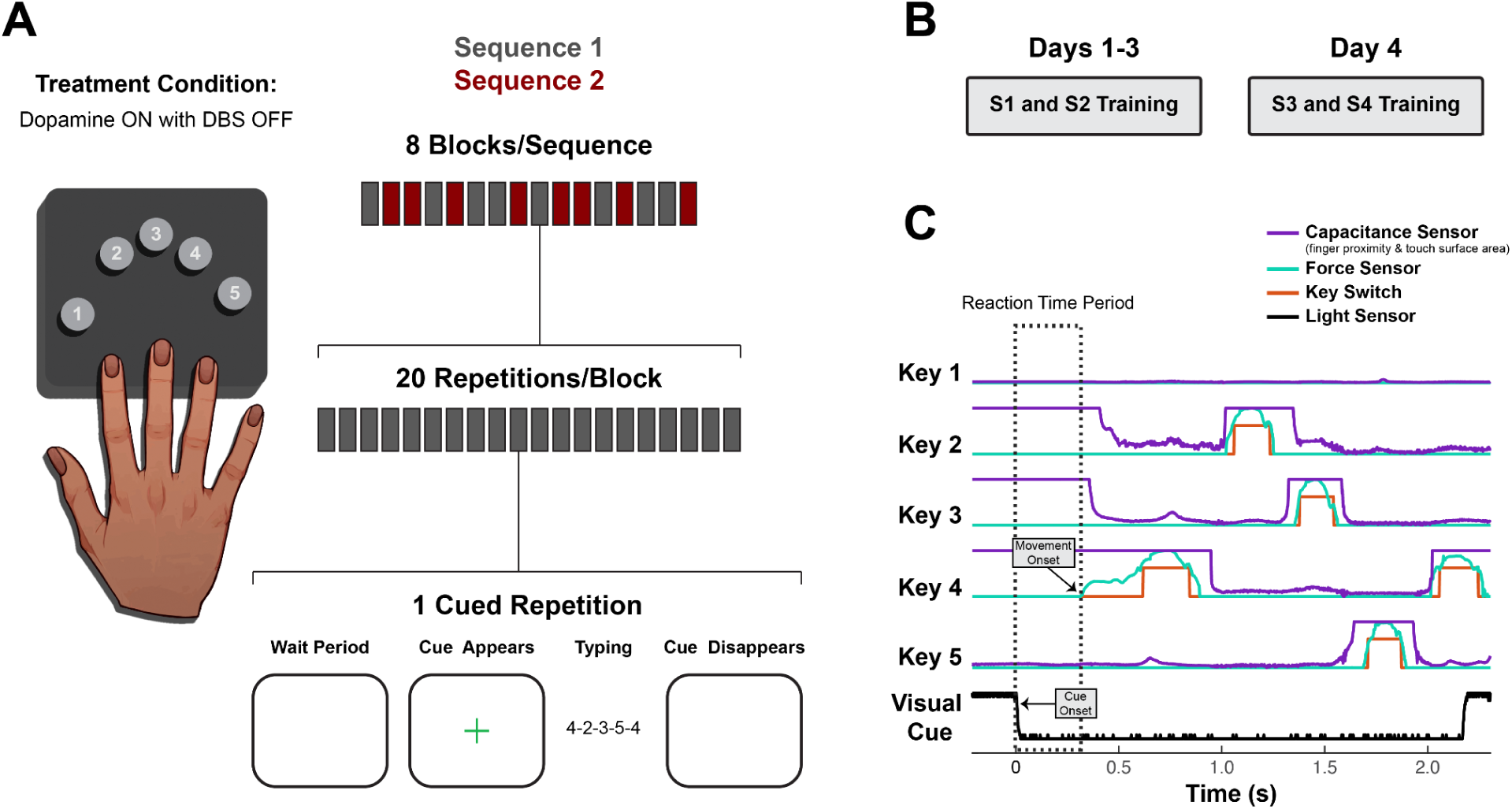
Experimental design and behavioral data collection. (**A**) On each day, subjects practiced typing two sequences. Interleaved practice blocks each contained 20 repetitions of visually cued sequence production for a single sequence. Subjects performed the task while on dopamine medication, and no DBS was delivered during the task or between days. (**B**) Days 1–3 employed novel Sequences 1 and 2 (S1 and S2), and Day 4 employed novel Sequences 3 and 4 (S3 and S4). (**C**) The reaction time period (dashed box) used for neural analysis is demonstrated in an example trial showing raw data from a custom behavioral setup used to capture finger movement (using capacitive proximity/touch sensors, force-sensitive resistors and mechanical key switches) and the visual cue (using a photodiode placed on the task computer screen). Capacitance sensors were calibrated to detect proximity changes of fingers hovering 0 to 2 cm above the keys (capacitance variation around low values). They could also detect changes in surface area of finger contact with the key associated with changes in force subthreshold for the force sensors (capacitance variation between low/mid-range and ceiling values). Thus, capacitance sensor readings were used for motor onset detection, except when motor onset began with a finger already in full contact with the key, in which case force sensor readings were used (as in the first keypress of this example trial).

They received two novel sequences on Day 1 and again on Day 4 (**Figure 3B**, **Table 2**). No sequences contained repeated adjacent elements, rising or falling triplets, or the thumb. All sequences paired for comparison within and between days started with the same first and last elements.

**Table 2.** Sequences. No sequences contained repeated adjacent elements, rising or falling triplets, or the thumb. All sequences paired for comparison within and across days started with the same first and last elements. Block order of S1 and S2 was switched on Day 2.

|  | Familiarization | Day 1 | Day 2 | Day 3 | Day 4 |
| --- | --- | --- | --- | --- | --- |
| **GP1** | 2-4-5-3-2<br>2-3-5-4-2 | 4-2-3-5-4<br>4-5-3-2-4 | 4-5-3-2-4<br>4-2-3-5-4 | 4-2-3-5-4<br>4-5-3-2-4 | 4-2-5-3-4<br>4-3-5-2-4 |
| **GP2** | 2-5-3-4-2<br>2-4-3-5-2 | 3-4-2-5-3<br>3-5-2-4-3 | 3-5-2-4-3<br>3-4-2-5-3 | 3-4-2-5-3<br>3-5-2-4-3 | 3-2-4-5-3<br>3-5-4-2-3 |
| **STN1** | 2-4-3-5-2<br>2-5-3-4-2 | 3-4-2-5-3<br>3-5-2-4-3 | 3-5-2-4-3<br>3-4-2-5-3 | 3-4-2-5-3<br>3-5-2-4-3 | 3-2-4-5-3<br>3-5-4-2-3 |
| **STN2** | 2-4-5-3-2<br>2-3-5-4-2 | 4-2-3-5-4<br>4-5-3-2-4 | 4-5-3-2-4<br>4-2-3-5-4 | 4-2-3-5-4<br>4-5-3-2-4 | 4-2-5-3-4<br>4-3-5-2-4 |

All subjects performed the experiment within one month after DBS surgery, before turning on DBS. No DBS was delivered during or between experimental sessions. To limit the effect of medication-related motor fluctuations, all experimental sessions were conducted at a consistent time across days within each subject’s medication ON period.

### Data collection

In each brain hemisphere, a four-contact DBS lead spanned basal ganglia (BG) nuclei, and a four-contact electrocorticography (ECoG) paddle spanned sensorimotor cortex (**Figure 2**). Bipolar recording of subcortical local field potentials (LFPs) and sensorimotor electrocorticography (ECoG) signal granted coverage of the following approximate regions in the left (contralateral) brain hemisphere. Subjects GP1 and GP2: globus pallidus (GP), putamen (Put), M1 or primary sensorimotor cortex (M1/S1), premotor cortex. Subjects STN1 and STN2: ventral subthalamic nucleus (vSTN), dorsal subthalamic nucleus (dSTN), M1, parietal cortex (spanning S1 and superior parietal lobule).

Leads from each brain hemisphere (Medtronic 3387 for globus pallidus, 3389 for subthalamic nucleus and 0913025 for cortex) were connected to a bidirectional neural interface in the ipsilateral chest (Medtronic Summit RC+S B35300R). LFP and ECoG signals were recorded at 500 Hz throughout the task. Channels were referenced to the metal casing of the implanted pulse generator. On-device hardware low and high pass filtered the data at 450 Hz and 0.85 Hz, amplified it, then performed another low pass filter at 1700 Hz. Task events and keystroke data were captured in 4-kHz sweeps using a portable custom-made device run by a Teensy 4.1 microcontroller (**Figure 3C**), which acted as the master clock and motherboard for a custom keypad, visual stimulus detector and electrical impulse detector. To ensure accurate detection of finger movement onset/offset, even when finger position started above but not touching the key, a combination of custom capacitive proximity sensors (carved from copper sheet metal, 3DDeluxe), force-sensitive resistors (FSRs, Alpha MF01A-N-221-A04) and linear mechanical key switches (CHERRY MX1A-LxxA/B) were used for each digit. An FSR was fixed atop each custom keycap (3DDeluxe).

A small resin disk with a centered bulge less than a millimeter tall was fixed atop each FSR. This ensured even that off-center finger contact with the key face would result in force distribution to the FSR’s center active zone sufficient to drive detectable FSR activity. The resin disks also insulated the FSRs from the proximity sensors, which were cut from copper sheet metal and fixed atop each resin disk. To maximize proximity sensor read rate, each proximity sensor was sampled by its own Teensy 3.2 microcontroller, which each transmitted readings to the Teensy 4.1. Proximity sensors were covered with insulating tape and calibrated to detect proximity changes of fingers hovering up to ∼2 cm above the keys. The capacitance sensors also detected changes in surface area of finger contact with the key. This enabled detection of changes in finger contact slight enough that the associated change in force was subthreshold for the force sensors. A photodiode (Everlight Electronics Co Ltd, PD333-3C/H0/L2) fixed to the task computer screen captured the timing of visual stimuli and progression of experimental epochs. For neural-behavioral data stream alignment, a unique temporal pattern of fifteen single DBS pulses was delivered at the start and end of each experimental session and detected along the metal casing of the pulse generator by an external electrical signal detector (MikroElectronika EEG Click MIKROE-3359). All sensors were calibrated and checked for electrical interference and cross-talk at the start of each experimental session.

### Behavioral analysis

To eliminate outlier trials, we excluded incorrect trials and any trials with a reaction time (RT, cue onset to movement onset) or trial duration (movement onset to offset) exceeding three standard deviations of the block average for correct trials.

To evaluate overall learning, we computed a block performance index for each subject. Each block performance index is the sum of block average error rate (1 – block accuracy), reaction time (cue presentation to movement initiation) and trial duration (movement onset to offset). For each subject, the block average trial durations and reaction times were each first min-max scaled to [0, 1], using data from all days to derive the minimum and maximum values. Lower performance index values indicate better performance.

### Neural analysis

All significance testing for neural analysis utilized permutation testing that simulates error within the null distribution, and secondary tests were performed only to assess the direction of primary detected effects. Multiple comparison correction was therefore not performed.

### Trial selection

In addition to the behavioral cutoffs applied for behavioral analysis, the following trial exclusion criteria and trial subsampling methods were performed for neural analysis. Trials with less than 25 ms between final/first movements associated with adjacent trials were excluded. Subsequently, subsampling was performed within a given day to match trial counts between sequences within each group of four blocks to avoid a possible imbalance over time, e.g., 75% of remaining Sequence 1 (S1) trials coming from the first half of the session and 75% of remaining Sequence 2 (S2) trials coming from the second half of the session. Within each group of four blocks, sequence subsampling followed epoch-specific selection rules. Only trials with RT ≥ 100 ms were considered. Trials from the higher count sequence were subsampled to match trials from the lower count sequence based on RT durations. Finally, random subsampling matched trial counts across days within each subject for each epoch type.

### Signal preprocessing

Neural signal preprocessing used the following pipeline. Data from each channel was linearly detrended, demeaned and high-pass filtered at 0.25 Hz using a two-pass FIR filter. Electrical noise was excluded in the frequency domain. Two-pass Kaiser FIR filters with normalized transition widths of ≤ 0.1 were used for all subsequent bandpass filtering.

Data intended for single-trial classification, amplitude analysis and undirected phase analysis was filtered with the following passbands: δ (0.5–4 Hz), theta (4–8 Hz), alpha (8–12 Hz) and β (12–30 Hz for amplitude analysis and cross-frequency coupling, 12–20 Hz and 20–30 Hz for single trial classification). For γ, filters were logarithmically spaced from 30 to 250 Hz. High γ (70–250 Hz) center frequencies were used for all analyses involving γ, while slow (30–50 Hz) and mid (50–70 Hz) γ center frequencies were used only for single-trial classification. These filters were all non-overlapping in the frequency domain to reduce collinearity between adjacent frequency bands when performing single-trial classification. The Hilbert transform computed the analytic signal. For undirected cross-frequency coupling (CFC) analysis using pairwise phase consistency (PPC), the resulting amplitude envelope of each β and γ center frequency was filtered with the same bandpass filter previously used to extract δ, followed by a second application of the Hilbert transform.

For analysis of directed phase coherence (including directed CFC) using phase slope index (PSI), δ was instead extracted using linearly spaced passbands (0.5, 3.25; 0.75, 3.5; 1, 3.75; 1.25, 4). For directed δ-β and δ-γ coupling analysis, these linearly spaced δ passbands were applied to the β amplitude envelope and to the amplitude envelope of each center frequency of high γ (70–250 Hz), followed by a second application of the hilbert transform.

For artifact screening, filter-Hilbert was used to estimate 70–250 Hz broadband amplitude, which was then *z*-scored over the entire session. Any trial in which *z* ever surpassed 8 standard deviations was omitted from neural analysis.

### Single-trial classification

If two sequences contain at least one differing element within the first few elements, then multi-element preplanning necessitates some sequence-specific neural activity. To test for neural activity related to multi-element preplanning, we thus tested for sequence-specific preparatory neural dynamics using single-trial classification of pre-movement neural activity. To evaluate learning-driven optimization of multi-element preplanning, we then evaluated change in sequence-specific predictive value of neural activity with practice by testing change in model performance across days.

A different classifier was trained on data from each recording channel on each day, and mean decoding accuracy was used to estimate the discriminability of sequence-specific neural activity. S1- or S2-labeled trial data was extracted from the RT period in the *t_n_* ms immediately prior to sequential movement onset, where *t_n_* was the average RT on Day 3 for Subject *n*. Data then underwent feature selection and logistic classification with *L_1_* regularization. For each subject, trials per sequence were balanced across classes, channels and days.

Time-frequency regions with maximal differences between sequences were selected as features. To assess, e.g., differences in narrowband amplitude dynamics in PM for S1 vs. S2 on Day 3 for GP1, we calculated the two-sided *t-*statistic for amplitude at each time-frequency point. For each of 10,000 permutations, trial labels were shuffled, and the *t*-statistic was recalculated. Thus, each time-frequency point had an associated null distribution of 10,000 *t*-statistic values. Time-frequency points at which the test value fell below the 80^th^ percentile compared to its respective null distribution were masked. In each of the remaining islands of features for each center frequency, the time point with the highest percentile score relative to its null was selected as a feature to use in the model. For all resulting features, corresponding amplitude values were taken from S1 and S2 trial data. For phase data, the same process was implemented, save for two differences. Phase opposition sum^55^ was used instead of the *t*-statistic, and since phase is a circular process, each selected phase value was converted into two features: sin(phase) and cos(phase). Each amplitude and phase feature was median-centered and scaled according to its interquartile range to have unit variance.

Hyperparameter optimization, model training and model testing were performed with nested cross validation. The inverse *L_1_* regularization constant (*λ^-1^*) was optimized per classifier in 10-fold, 10-repeat stratified cross validation performed on a stratified 90% subset of the data. The following *λ^-1^* values were tested: 5E-2, 1E-1, 5E-1, 1, 5, 1E1, 5E1, 1E2, 5E2, 1E3, 5E3, 1E4, 5E4, 1E5, 5E5. Greater shrinkage produced performance at or below chance level. The selected value for *λ^-1^* was then used for final model training and testing on the full dataset with 10-fold, 100-repeat stratified cross validation.

Permutation testing evaluated for significant sequence-specific neural activity in each brain region and how the predictive value of neural activity changed with practice. Right-sided permutation testing assessed significance of model mean decoding accuracy relative to chance. The outer 10-fold, 100-repeat stratified cross validation was repeated 1,000 times with permuted trial labels, and the resulting 1,000 null values were compared to test sample mean decoding accuracy. Two-sided permutation testing assessed the change in mean decoding accuracy across days for a given channel. Mean decoding accuracies for each of the 100 repeats per day were permuted across days, and the between-day difference in overall mean decoding accuracy was recalculated for each permutation as the null value.

### Feature importance testing

To evaluate how learning-driven changes in sequence-specific neural activity might be reflected in the spectral characteristics of field potential recordings, the absolute importance of various signal properties to the performance of trained models was tested and compared across days. We permuted, in the test set, the trial labels for all phase *or* amplitude features associated with a given canonical frequency band, as the majority of across-frequency or across-phase/amplitude feature correlations were not high (ρ < 0.5). For each fold in each repeat of the 10-fold 100-repeat outer cross-validation used for prior model training and testing, the test data trial labels for the respective trained model were permuted once for a given feature group, and the resulting change in test accuracy from test performance was computed. Change in decoding accuracy was then averaged across all 10 folds in each of the 100 repeats. Feature groups with negligible negative accuracy decreases were set to zero in data plots for visual clarity. No groups showed a negative accuracy decrease greater than 1% in any model.

We then repeated group permutation testing, except with all phase and amplitude features for δ through β grouped together and likewise for low γ through high γ. Change in decoding accuracy was averaged across folds per repeat before between-day permutation testing.

### Coherence analysis

Functional connectivity was evaluated over the course of practice to assess which network interactions may support learning-driven optimization of multi-element preplanning. Single-trial plots indicated that δ phase aligned to cue in various regions, so data was aligned to cue and evaluated in a window length of the mean RT of Day 3 per subject. Nonparametric cluster-based permutation across time, with a cluster size correction, was used for all phase analyses. To simplify data visualization, significant across-day effects associated with low and insignificant levels of within-day local or interregional coherence were not typically depicted with shaded time regions in the figures, but the *p*-values are still reported in the data tables.

For all phase analyses, we first computed each metric within sequence before averaging the resulting time series across sequences prior to statistical testing. This was intended to address two main issues. First, we expected possible sequence-specificity in spatiotemporal patterns of neural activity that could be reflected in mesoscale spectral activity—an idea for which both single-trial classification and single-trial δ phase plots then provided confirmatory evidence. This implies that different sequences could be associated with different characteristic γ amplitude envelope morphologies, which may display different phase-specific coupling patterns with δ. Second, in cases for which two sequences were not performance-matched on a given day (e.g., Sequences 3 and 4), one may observe differences in activity between sequences due to performance level (rather than learning stage). In either case, a reasonable approach would be to respect the sequence-specific relationships, so we first computed metrics within each sequence. However, we also expected the general oscillatory network dynamics associated with learning to be the same regardless of sequence, so we then averaged the resulting metric time series across sequences before statistical testing.

Inter-trial δ phase locking value (PLV) assessed cue-aligned consistency of local δ phase^56^. For a given recording channel, the resulting time series (one for each sequence) were smoothed with a 150 ms-long Gaussian window and averaged across sequences. To test for significant PLV on a given day, we randomly sampled phase data from the duration of the session for each permutation. PLV was computed for each sequence null group using the appropriate number of trials for each sequence. The magnitudes of the resulting two null PLV time series were then smoothed and averaged across trial groups to attain a single null time series for that permutation. This was repeated for each permutation. To test for a significant difference in PLV time series between days, we calculated the test time series by subtracting the PLV time series from one day from that of the other day. For each permutation, trials were shuffled across days but within sequence.

Inter-trial δ pairwise phase consistency assessed cue-aligned interregional δ phase coherence^57^. For each channel pair, the resulting time series (one for each sequence) were smoothed with a 150 ms-long Gaussian window and averaged across sequences. For baseline PPC testing, methods were identical to those used in baseline PLV testing, except the null was constructed by sampling channel data as pairs, i.e., the baseline distribution corresponded to an actual estimate of baseline session-wide coherence for that channel pair, not to the level of coherence that would be expected if the two channels were coupled only randomly. For between-day PPC permutation testing, test time series were calculated by subtracting the PPC time series between days, and for each permutation, phase data for both channels in a given channel pair were shuffled together across days but within sequence.

To assess the direction of pairwise δ phase relationships observed with PPC, phase slope index between two channels was computed per time point for each sequence^58^. PSI values were not smoothed before being averaged across sequences. To assess whether significant δ phase lead/lag occurred with respect to chance, i.e., neither channel led the other, rather than with respect to session baseline, data for each permutation was randomly sampled from the session duration separately for each channel in the pair, for each sequence.

Pairwise phase consistency and phase slope index were also used to estimate undirected and directed coherence, respectively, for cross-frequency couplings. δ phase was paired with the δ phase of the β or γ amplitude envelope. Computations analogous to those used for δ synchrony analysis were performed, except for two modifications. For δ-high γ coupling, PPC or PSI was calculated separately for each narrowband within 70–250 Hz for each sequence. The result was averaged across γ center frequencies before averaging across sequences. Second, for baseline permutation testing, the phase data was held constant while the amplitude data was sampled from the session. Shuffling amplitude while holding phase constant was intended to test for significant coherence *given* a specific phase distribution.

### Use of artificial intelligence tools

DALL-E generated the first draft of the hand in **Figure 3A**. After initial drafting of the manuscript, ChatGPT and Gemini were used to improve text concision.

## RESULTS

### Behavioral stratification based on sequence learning

Subjects were behaviorally stratified for sequence learning based on changes in a composite measure of block performance—a block performance index—for which lower value corresponds to better performance (**Figure 4A**). Day 1 to Day 3 comparisons of pooled block performance indices for each of S1 and S2 evaluated within-sequence practice-driven performance changes. Significant decrease in performance index suggests sequence learning, but improvement on S1 and S2 could also have been driven by more general task learning, e.g., optimization of task-related cognitive processes and motor familiarization with the experimental apparatus^19,59–64^. Even so, superior performance of S1 and S2 on Day 3 compared to that upon subsequent presentation of novel sequences on Day 4 would suggest some sequence learning had in fact occurred for S1 and S2. Thus, subjects were labeled improvers (ID ending in 1) only if their performance both improved from Day 1 to Day 3 and worsened when presented with novel sequences on Day 4. A Day 1 to Day 4 comparison to assess general task learning is confounded by behavioral interference between the familiar sequences and those presented on Day 4, so we did not attempt to behaviorally stratify based on task learning.

**Figure 4.**
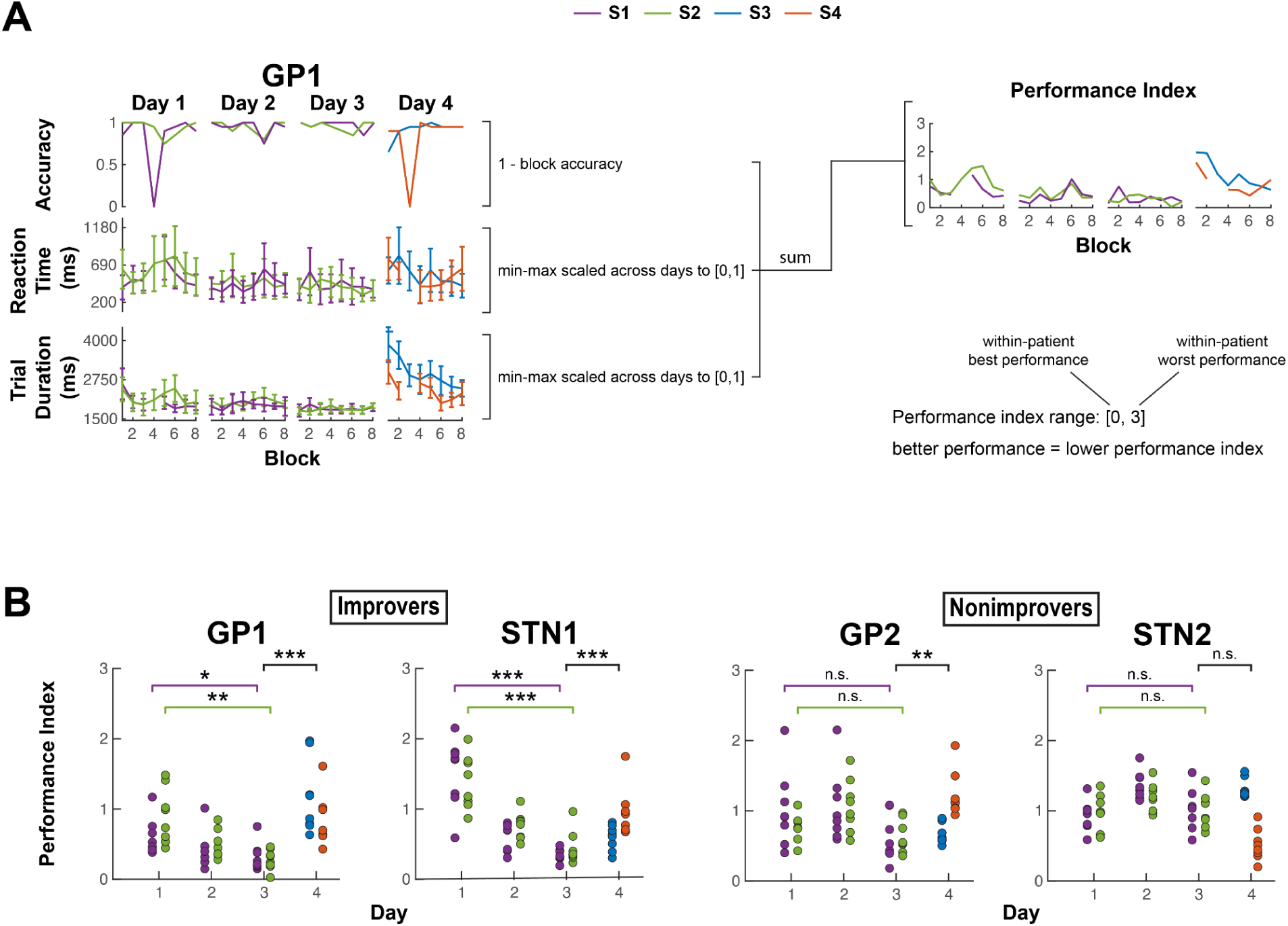
Behaviorally distinguishing improvers and nonimprovers. **(A)** Example calculation of block performance index (PI) from block average data. Each block performance index is the sum of block average error rate [1 – accuracy], reaction time [cue onset to movement onset] and trial duration [movement onset to offset]. For each subject, the block average trial durations and reaction times were each first min-max scaled to [0, 1], using data from all days to derive the minimum and maximum values. Error bars indicate ± *s*. **(B)** Block performance index for each subject. Comparison across Days 1 and 3 for each of S1 and S2 assessed within-sequence practice-driven performance changes. To help evaluate whether performance changes from Day 1 to Day 3 were at least in part related to sequence learning and not solely attributable to changing familiarity with the task and keypad, performance was also compared between pooled Day 3 sequences and pooled Day 4 sequences. Subjects were labeled improvers (ID ending in 1) only if their performance both improved from Day 1 to Day 3 and worsened when presented with novel sequences on Day 4 (For all comparisons: *α =* 0.05, two-sided, two-sample *t*-test with unequal variance. For within-sequence comparisons and pooled sequence comparisons, *n* = 8 and 16 sequence blocks per group, respectively, except for GP1 Day 1 S1 and GP1 Day 4 for which *n* = 7 and 15 due to exclusion of 0% accuracy blocks, as composite performance would be poorly defined). \**p* < 0.05, \*\**p* < 0.01, \*\*\**p* < 0.001.

GP1 and STN1 showed indications of sequence learning, whereas GP2 and STN2 did not (**Figure 4B, Supplementary Figures 1–2**). In GP1 (S1: *p* = 0.032; S2: *p* = 0.002) and STN1 (S1: *p* < 0.001; S2: *p* < 0.001), performance index improved for both sequences from Day 1 to Day 3, then worsened when practicing novel sequences on Day 4 (GP1: *p* < 0.001; STN1: *p* < 0.001). Neither GP2 nor STN2 showed significant improvements in Sequence 1 (S1) and Sequence 2 (S2) performance index (*p* > 0.05 for all), and only GP2’s performance index significantly differed between Days 3 and 4 (GP2: *p* = 0.005; STN2: *p* > 0.05). These results suggest behavioral stratification as follows: improvers (GP1 and STN1) and nonimprovers (GP2 and STN2).

Sleep durations and end-of-session upper limb scores on the Movement Disorders Society Unified Parkinson’s Disease Rating Scale (MDS-UPDRS) were similar between groups^65^ (**Table 3**). This suggests relative differences in performance improvement may not have been due to large differences in sleep or motor symptom presentation in the typing arm.

**Table 3.**
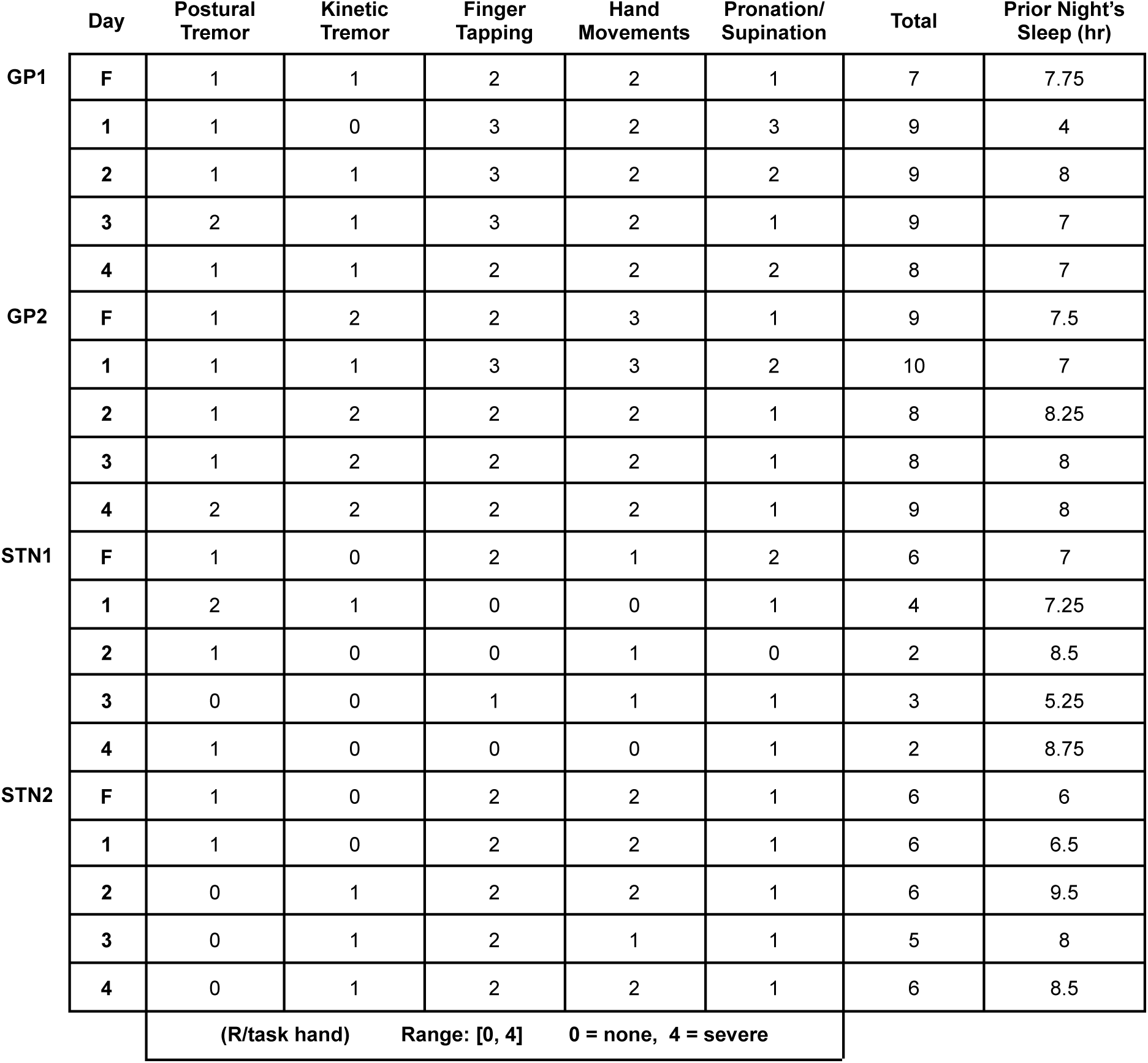
Supplementary data collection. Scores for the upper limb component of the UPDRS performed immediately after the typing task each day and the prior night’s sleep duration, collected with a sleep journal. F, Familiarization.

### Pre-movement cortical and basal ganglia γ activity is sequence-specific and increasingly predictive of sequence content with performance improvement

We next assessed which brain regions might participate in multi-element preplanning and how this changes with learning. We used single-trial classification of pre-movement neural activity to predict the identity of the upcoming sequence. Model performance thus quantified sequence-specific predictive value of neural activity. For each experiment day and each recording channel per subject, we performed feature selection on neural data preceding sequential movement onset and trained a model to predict the identity of the sequence that the subject was about to type (**Figure 5A–B**, **Supplementary Figure 3**). To isolate practice-driven changes in sequence-specific neural activity from changes that occur as a byproduct of the changing behavior, we only directly compared neural activity between sequences for which overall behavioral performance was similar. In most subjects, performance levels significantly differed between Sequences 3 and 4 (Days 1–3: *p* > 0.05 in all subjects; Day 4: *p* > 0.05 in GP1, *p* = 0.002 in GP2, *p* = 0.031 in STN1, *p* < 0.001 in STN2) (**Supplementary Figure 4**), so Day 4 data was excluded from single-trial classification analysis. To reduce the influence of neural activity related to the first sequence element, we designated the same digit as the first element in all sequences for each subject. See *Methods: Neural analysis* for additional measures taken to reduce the influence of confounds.

**Figure 5.**
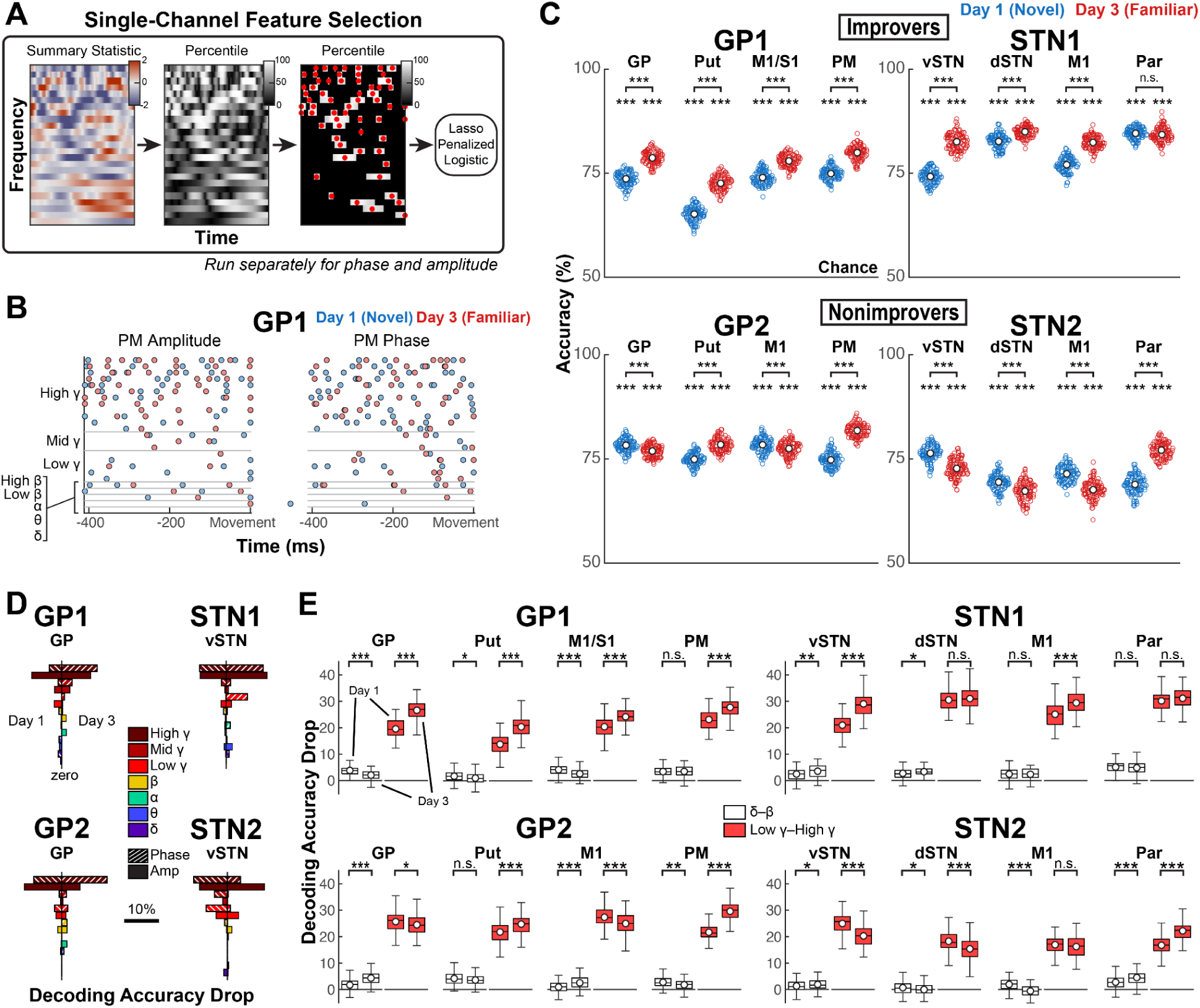
Pre-movement sequence-specific γ activity, present in all brain regions, demonstrates practice-driven increases and decreases in discriminability in improvers and nonimprovers, respectively. (**A**) Visualization of feature selection pipeline. Features were selected separately for each channel on each day in each subject. (Left) For selection of amplitude features, the S1 vs. S2 two-sided *t*-statistic was computed at each time-frequency point. (Middle) The *t*-statistic at each time-frequency point was recomputed for 10,000 permutations of trial labels to determine the percentile ranking of the test value at each time-frequency point relative to its null distribution. (Right) Time-frequency points falling below their respective 80th-percentile cutoffs were masked, and in each of the remaining time-frequency regions, the time point achieving the highest percentile was selected as an amplitude feature. This process was repeated for phase data, using phase opposition sum as the summary statistic. Each resulting phase feature was split into two features that corresponded to the cartesian phase coordinates. Classification utilized 10-fold 100-repeat lasso-penalized logistic classification. (**B**) Example selected features for Days 1 and 3 in GP1’s PM. (**C**) Mean decoding accuracy per model after feature selection (Comparison to chance: *α =* 0.05, one-sided, permutation testing with 1,000 resamples. Comparison across days: *α =* 0.05, two-sided, permutation test with 10,000 resamples. Empty circle reflects mean decoding accuracy across folds for one repeat; white circle reflects mean decoding accuracy across repeats. (**D**) Absolute decoding accuracy decreases for features grouped by canonical frequency band and signal property (amplitude or phase)—a subset of representative plots. (**E**) Absolute decoding accuracy decreases for features grouped by frequency into two groups: δ through β (0.5–30 Hz, phase and amplitude) and low γ through high γ (30–250 Hz, phase and amplitude) (Comparison across days: *α =* 0.05, two-sided, permutation testing with 10,000 resamples.). White circle reflects mean; black horizontal line reflects median. Box edges correspond to 25th and 75th percentiles. Whiskers span the entire data range excluding outliers. Outliers were computed as 1.5·*IQR* away from the upper or lower quartile and are not shown. For all analysis in this figure, *n* sequence trials per day = 238 for GP1, 206 for GP2, 158 for STN1, 182 for STN2. \**p* < 0.05, \*\**p* < 0.01, \*\*\**p* < 0.001.

We compared each model’s performance to chance and tested within-channel change in decoding accuracy across Days 1 and 3, with the caveat that Day 1 to Day 3 changes may reflect effects of both sequence practice and task exposure. Notably, sequence-specific activity was detected throughout the recorded network in all subjects (*p* < 0.001 for all models) (**Figure 5C**). Practice drove nearly network-wide increases of this activity’s predictive value in improvers (*p* < 0.001 for all except Par; STN1 Par: *p* > 0.05) but decreases in GP (*p* < 0.001), STN (*p* < 0.001 for vSTN and dSTN) and M1 (*p* < 0.001 for GP2 and STN2) in nonimprovers (significant increase in nonimprovers’ other channels with *p* < 0.001).

To assess which electrophysiological signal properties granted sequence-specific predictive value, we performed feature analysis. Minimal correlations (ρ < 0.5) between features grouped by phase, amplitude and canonical frequency band allowed grouped feature permutation testing (**Supplementary Figure 5**). Low γ (30–50 Hz), mid γ (50–70 Hz) and high γ (70–250 Hz) were collectively the most important, though the relative importance of γ phase and amplitude varied (**Figure 5D**, **Supplementary Figure 6**). Thus, for formal statistical analysis, we divided features into only two groups: 1) δ through β and 2) low γ through high γ and found that γ activity was the primary driver of practice-driven changes in model performance (**Figure 5E**, **Supplementary Figure 7**). In most brain regions, the across-day change in accuracy drop linked to γ features was large and in a direction that paralleled practice-driven change in model performance (GP1: *p* < 0.001 in all brain regions; GP2: *p* = 0.025 in GP, *p* < 0.001 otherwise; STN1: *p* < 0.001 in vSTN and M1, *p* > 0.05 in dSTN and Par; STN2: *p* > 0.05 in M1, *p* < 0.001 otherwise). In contrast, effects for δ through β features were small and did not consistently track model performance across days (GP1: *p* < 0.001 in GP and M1/S1, *p* = 0.013 in Put, *p* > 0.05 in PM; GP2: *p* < 0.001 in GP and M1, *p* > 0.05 in Put, *p* = 0.001 in PM; STN1: *p* = 0.004 in vSTN, *p* = 0.020 in dSTN, *p* > 0.05 in M1 and Par; STN2: *p* = 0.049995 in vSTN, *p* = 0.024 in dSTN, *p* < 0.001 in M1 and Par). These findings suggest network-wide participation in multi-element preplanning through sequence-specific population activity that was optimized with learning and reflected in γ activity.

### Improvement is associated with cortically-led δ phase synchrony in response to cue

We next evaluated how oscillatory network dynamics may have facilitated the observed learning-related changes in sequence-specific activity. Assuming that the overall architecture of functional connectivity is sequence-general even for sequence-specific spatiotemporal patterns of neural activity, we calculated functional connectivity for each sequence and averaged the result across sequences per day before statistical testing. This minimized confounds due to performance differences between learning stage-matched sequences. Thus, we could analyze all experimental sessions, and across-day comparisons of functional connectivity paralleled behavioral stratification. Sequence practice-related effects were those that occurred in overlapping time regions between *both* Days 1 and 3 and Days 3 and 4; though we did not behaviorally stratify task learning, we did examine task exposure-related neural activity by comparing Days 1 and 4.

The supposition that δ phase facilitates recruitment of sequence-specific ensemble activity implies δ phase-spike coding, which ultimately predicts that δ phase consistently aligns to motor events. Cortical δ phase locks to movement-related visual cues in healthy subjects, so we computed δ phase locking value (PLV) to cue in our subjects after confirming all recording channels had δ amplitude sufficient for phase estimation^36,39,56,66^ (**Figure 6A**, **Supplementary Figures 8–9**). In GP subjects, PLV was significant throughout motor cortex on all days, increasing with task exposure in M1 (or M1/S1). However, only GP1 had significant PLV in basal ganglia on all days and increasing PLV in GP and PM with task exposure. In STN subjects, M1 PLV was also significant on all days. In other channels (not surgically targeted in GP subjects), PLV effects differed between STN1 and STN2. In STN1, PLV increased with task exposure in vSTN and Par and with both sequence practice and task exposure in dSTN—effects absent in STN2, for whom PLV was mostly insignificant or diminished with task exposure. These results suggest that cue-aligned δ phase in motor cortex, striatum and, increasingly, dSTN was important to sequence learning and reveal an enhancement of motor network-wide δ phase alignment to cue with task exposure that occurred only in improvers.

**Figure 6.**
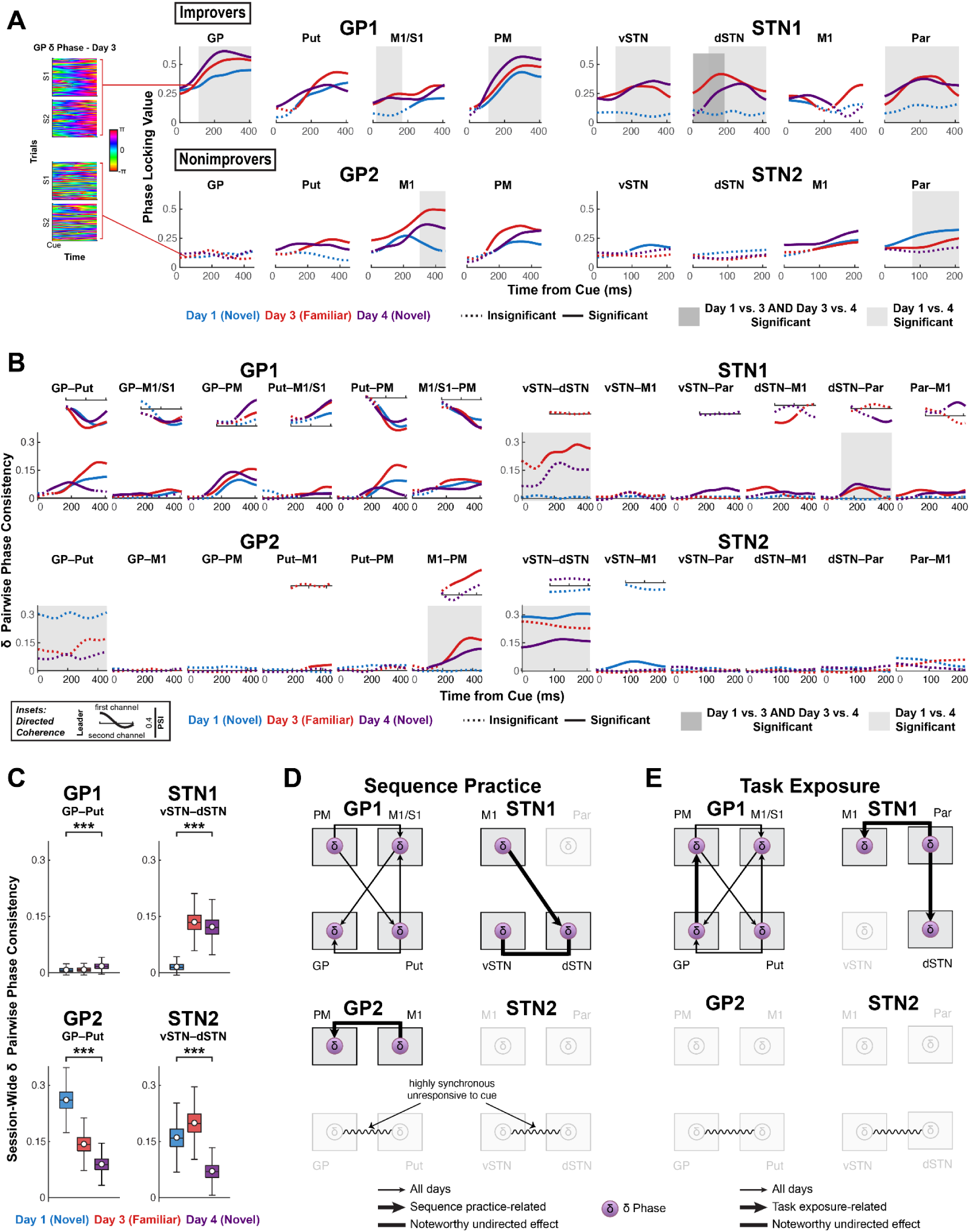
Improvement is associated with cortically-led network δ phase synchrony, to which sequence learning and task exposure add distinct effects, while lack of improvement is associated with highly synchronous BG δ. (**A**) (Left) All Day 3 single-trial δ phase time series after cue onset in the pallidum of GP1 and GP2. (Right) Phase locking value (PLV) after cue onset on Days 1, 3 and 4. Solid line indicates PLV significantly higher than chance (*α =* 0.05, one-sided, cluster-based permutation test with 10,000 resamples. See **Table 4** for *p*-values.). Shaded box indicates significant difference in PLV between days (α = 0.05, two-sided, cluster-based permutation with 10,000 resamples. See **Table 5** for *p*-values.). (**B**) Effects in δ synchrony. (Large plots) δ pairwise phase consistency (PPC, undirected measure) time series for all channel pairs. Solid line indicates PPC significantly higher than session-wide baseline, i.e., *h_0_* = coherence aligned cue is the same as general coherence levels not aligned to cue (rather than *h_0_* = no coherence) (α = 0.05, one-sided, cluster-based permutation with 10,000 resamples. See **Table 6** for *p*-values.). Shaded box indicates significant difference in PPC between days (α = 0.05, two-sided, cluster-based permutation with 10,000 resamples. See **Table 7** for *p*-values.). (Insets) Phase slope index (PSI, directed measure) for significant PPC time series. Solid line indicates significant PSI (*h_0_*= no channel leads, α = 0.05, two-sided, cluster-based permutation with 10,000 resamples. See **Table 8** for *p*-values.). (**C**) Session-wide baseline basal ganglia δ pairwise phase consistency averaged across time. Calculated by taking the null distribution of time series resampled from each session in (**B**) and averaging each null PPC sample across time. Change in session baseline across Days 1 and 4 was tested (*α =* 0.05, two-sided, permutation testing with 10,000 resamples. See **Table 9**: *p*-values.). White circle reflects mean; black horizontal line reflects median. Box edges correspond to 25th and 75th percentiles. Whiskers span entire data range excluding outliers. Outliers were computed as 1.5·*IQR* away from the upper or lower quartile and are not shown. \*\*\**p* < 0.001. (**D**) Network diagrams illustrating sequence practice-related δ coherence effects. (**E**) Network diagrams illustrating task exposure-related δ coherence effects.

**Table 4.** Baseline testing of local cue-related δ phase locking value: *p*-values. Blank cells indicate no time regions passed initial thresholding. Multiple values in a single cell correspond to multiple time regions that passed initial thresholding. Earlier time regions are listed first.

|  |  | *p*-value |  |  |  |
| --- | --- | --- | --- | --- | --- |
|  | Region | *n* | Day 1 | Day 3 | Day 4 |
| **GP1** | GP | 238 | **0.000100** | **0.000100** | **0.000100** |
|  | Put | 238 | **0.002300** | **0.002500** | **0.000100** |
|  | M1/S1 | 238 | **0.015898** | **0.000100** | **0.000100** |
|  | PM | 238 | **0.000500** | **0.000500** | **0.000900** |
| **GP2** | GP | 202 |  | 0.155084 |  |
|  | Put | 202 |  | **0.009099** | **0.000100** |
|  | M1 | 202 | **0.001400** | **0.000100** | **0.000300** |
|  | PM | 202 | **0.003100** | **0.001700** | **0.004000** |
| **STN1** | vSTN | 192 |  | **0.000100** | **0.000100** |
|  | dSTN | 192 |  | **0.000100** | **0.001200** |
|  | M1 | 192 | **0.045195**<br>**0.130987** | **0.042196**<br>**0.036096** | **0.008199** |
|  | Par | 192 | 0.120588<br>0.158384 | **0.000100** | **0.000100** |
| **STN2** | vSTN | 182 | **0.034497** |  | 0.098290 |
|  | dSTN | 182 |  |  |  |
|  | M1 | 182 | **0.024198** | **0.021398** | **0.000100** |
|  | Par | 182 | **0.000100** | **0.000100** | 0.106189<br>0.078192 |

**Table 5.** Across-day testing of local cue-related δ phase locking value: *p*-values. Blank cells indicate no time regions passed initial thresholding. Multiple values in a single cell correspond to multiple time regions that passed initial thresholding. Earlier time regions are listed first.

|  |  | *p*-value |  |  |  |
| --- | --- | --- | --- | --- | --- |
|  | Region | *n* | Day 1 vs. 3 | Day 3 vs. 4 | Day 1 vs. 4 |
| **GP1** | GP | 238 | **0.035096** |  | **0.002700** |
|  | Put | 238 | 0.195580<br>0.075092 | 0.082892 | 0.094291 |
|  | M1/S1 | 238 | **0.0150985**<br>0.195780 |  | **0.041896** |
|  | PM | 238 |  |  | **0.002400** |
| **GP2** | GP | 202 |  |  |  |
|  | Put | 202 | **0.043596** |  | 0.170983<br>0.170983 |
|  | M1 | 202 | **0.020798** | 0.074493 | **0.044496** |
|  | PM | 202 | **0.022298** | 0.206479 |  |
| **STN1** | vSTN | 192 | **0.000100** |  | 0.189781<br>**0.001500** |
|  | dSTN | 192 | **0.000100** | **0.022098** | **0.001200** |
|  | M1 | 192 | 0.108189 | **0.032097** |  |
|  | Par | 192 | **0.000600** |  | **0.000100** |
| **STN2** | vSTN | 182 |  |  |  |
|  | dSTN | 182 |  |  |  |
|  | M1 | 182 |  |  |  |
|  | Par | 182 |  |  | **0.017498** |

**Table 6.** Baseline testing of δ PPC: *p*-values. Blank cells indicate no time regions passed initial thresholding. Multiple values in a single cell correspond to multiple time regions that passed initial thresholding. Earlier time regions are listed first.

|  |  | *p*-value |  |  |  |
| --- | --- | --- | --- | --- | --- |
|  | Region | *n* | Day 1 | Day 3 | Day 4 |
| **GP1** | GP-Put | 238 | **0.001400** | **0.001300** | **0.003200** |
|  | GP-M1/S1 | 238 | **0.016798** | **0.001800** | **0.007499**<br>0.124588 |
|  | GP-PM | 238 | **0.004500** | **0.000400** | **0.000400** |
|  | Put-M1/S1 | 238 | 0.094291<br>**0.042696** | **0.005799** | **0.012799** |
|  | Put-PM | 238 | **0.004100** | **0.004900** | **0.002000** |
|  | M1/S1-PM | 238 | **0.002900** | **0.004100** | **0.000500** |
| **GP2** | GP-Put | 202 |  |  |  |
|  | GP-M1 | 202 |  |  |  |
|  | GP-PM | 202 |  |  |  |
|  | Put-M1 | 202 |  | **0.037796** |  |
|  | Put-PM | 202 | 0.164084<br>0.106589 |  | 0.060794 |
|  | M1-PM | 202 |  | **0.004500** | **0.000600** |
| **STN1** | vSTN-dSTN | 192 |  | 0.174083<br>**0.005999** | 0.126987 |
|  | vSTN-M1 | 192 |  | 0.076692 | 0.054195 |
|  | vSTN-Par | 192 |  |  | **0.001900** |
|  | dSTN-M1 | 192 | 0.180782 | **0.010099** | **0.002100** |
|  | dSTN-Par | 192 |  | **0.003400** | **0.000600** |
|  | Par-M1 | 192 |  | **0.000300** | **0.002400** |
| **STN2** | vSTN-dSTN | 182 | **0.000100** |  | **0.000100** |
|  | vSTN-M1 | 182 | **0.005699** |  |  |
|  | vSTN-Par | 182 | 0.130587 |  | 0.074693 |
|  | dSTN-M1 | 182 |  |  |  |
|  | dSTN-Par | 182 | 0.088791 |  | 0.111189<br>0.087591 |
|  | Par-M1 | 182 |  |  |  |

**Table 7.** Across-day testing of δ PPC: *p*-values. Blank cells indicate no time regions passed initial thresholding. Multiple values in a single cell correspond to multiple time regions that passed initial thresholding. Earlier time regions are listed first.

|  |  | *p*-value |  |  |  |
| --- | --- | --- | --- | --- | --- |
|  | Region | *n* | Day 1 vs. 3 | Day 3 vs. 4 | Day 1 vs. 4 |
| **GP1** | GP-Put | 476 | 0.121488 | **0.022398** | 0.089191 |
|  | GP-M1/S1 | 476 |  |  |  |
|  | GP-PM | 476 | 0.182882 |  | 0.073093 |
|  | Put-M1/S1 | 476 | 0.140886 |  | 0.075392 |
|  | Put-PM | 476 | 0.105789 | **0.028097** | 0.092891 |
|  | M1/S1-PM | 476 |  |  |  |
| **GP2** | GP-Put | 404 | **0.000100** | 0.118988 | **0.000100** |
|  | GP-M1 | 404 |  |  |  |
|  | GP-PM | 404 | **0.006199** |  | **0.028197**<br>**0.007699** |
|  | Put-M1 | 404 | **0.022098** | 0.242376 | 0.208179 |
|  | Put-PM | 404 |  |  | 0.067693 |
|  | M1-PM | 404 | **0.001700** |  | **0.000400** |
| **STN1** | vSTN-dSTN | 384 | **0.000100** | 0.053395<br>0.095890 | **0.000100** |
|  | vSTN-M1 | 384 |  | 0.161384 |  |
|  | vSTN-Par | 384 |  | **0.017998** | 0.174183 |
|  | dSTN-M1 | 384 | **0.020598** | 0.130387 |  |
|  | dSTN-Par | 384 | **0.005199** | 0.197480<br>0.138986 | **0.000600** |
|  | Par-M1 | 384 | 0.185581 |  |  |
| **STN2** | vSTN-dSTN | 364 |  | 0.086191 | **0.000100** |
|  | vSTN-M1 | 364 | **0.006799** | 0.115288 | 0.069793 |
|  | vSTN-Par | 364 |  |  |  |
|  | dSTN-M1 | 364 |  |  |  |
|  | dSTN-Par | 364 |  |  | 0.163184 |
|  | Par-M1 | 364 |  | 0.131287 |  |

**Table 8.** Baseline testing of δ PSI: *p*-values. Blank cells indicate no time regions passed initial thresholding. Multiple values in a single cell correspond to multiple time regions that passed initial thresholding. Earlier time regions are listed first. NA, not applicable.

|  |  | *p*-value |  |  |  |
| --- | --- | --- | --- | --- | --- |
|  | Region | *n* | Day 1 | Day 3 | Day 4 |
| **GP1** | GP-Put | 238 | 0.081592<br>**0.000300** | **0.000100** | **0.000300** |
|  | GP-M1/S1 | 238 | **0.003400** | **0.001600** | **0.000800** |
|  | GP-PM | 238 |  | **0.032897** | **0.005799** |
|  | Put-M1/S1 | 238 | **0.018798** | 0.088191<br>**0.001300** | **0.001600** |
|  | Put-PM | 238 | **0.001500** | **0.000800** | **0.000300** |
|  | M1/S1-PM | 238 | **0.000400** | **0.000600** | **0.000600** |
| **GP2** | GP-Put | 202 | NA | NA | NA |
|  | GP-M1 | 202 | NA | NA | NA |
|  | GP-PM | 202 | NA | NA | NA |
|  | Put-M1 | 202 | NA |  | NA |
|  | Put-PM | 202 | NA | NA | NA |
|  | M1-PM | 202 | NA | **0.000600** | 0.089691 |
| **STN1** | vSTN-dSTN | 192 | NA |  | NA |
|  | vSTN-M1 | 192 | NA | NA | NA |
|  | vSTN-Par | 192 | NA | NA |  |
|  | dSTN-M1 | 192 | NA | **0.004400** | 0.059394 |
|  | dSTN-Par | 192 | NA |  | **0.012499** |
|  | Par-M1 | 192 | NA |  | **0.013399** |
| **STN2** | vSTN-dSTN | 182 |  | NA |  |
|  | vSTN-M1 | 182 |  | NA | NA |
|  | vSTN-Par | 182 | NA | NA | NA |
|  | dSTN-M1 | 182 | NA | NA | NA |
|  | dSTN-Par | 182 | NA | NA | NA |
|  | Par-M1 | 182 | NA | NA | NA |

**Table 9.** Across-day testing of session-wide null PPC: *p*-values.

|  | *p*-value |
| --- | --- |
| GP1 | 0.000100 |
| GP2 | 0.000100 |
| STN1 | 0.000100 |
| STN2 | 0.000100 |

**Table 10.** Baseline testing of intraregional δ-γ ^δ^ PPC: sequence learning-related *p*-values. Blank cells indicate no time regions passed initial thresholding. Multiple values in a single cell correspond to multiple time regions that passed initial thresholding.

|  |  | *p*-value |  |  |  |
| --- | --- | --- | --- | --- | --- |
|  | Region | *n* | Day 1 | Day 3 | Day 4 |
| GP1 | GP | 238 |  |  |  |
|  | Put | 238 | 0.284870 |  |  |
|  | M1/S1 | 238 |  | **0.000400** | **0.046695** |
|  | PM | 238 |  | 0.097890 | 0.203880 |
| GP2 | GP | 202 | 0.19548<br>**0.034697** |  |  |
|  | Put | 202 |  |  | 0.198180 |
|  | M1 | 202 |  | 0.297770<br>**0.001100** | 0.058994 |
|  | PM | 202 |  | 0.130990 |  |
| STN1 | vSTN | 192 | 0.184380 |  |  |
|  | dSTN | 192 |  |  | 0.304670 |
|  | M1 | 192 |  | 0.070593 | **0.024998** |
|  | Par | 192 | **0.041596** | 0.166280 |  |
| STN2 | vSTN | 182 |  |  |  |
|  | dSTN | 182 | **0.000100** | 0.143390<br>**0.010499** |  |
|  | M1 | 182 | **0.006999** | **0.000100** | **0.000100** |
|  | Par | 182 |  | **0.040096** | **0.002900** |

**Table 11.**
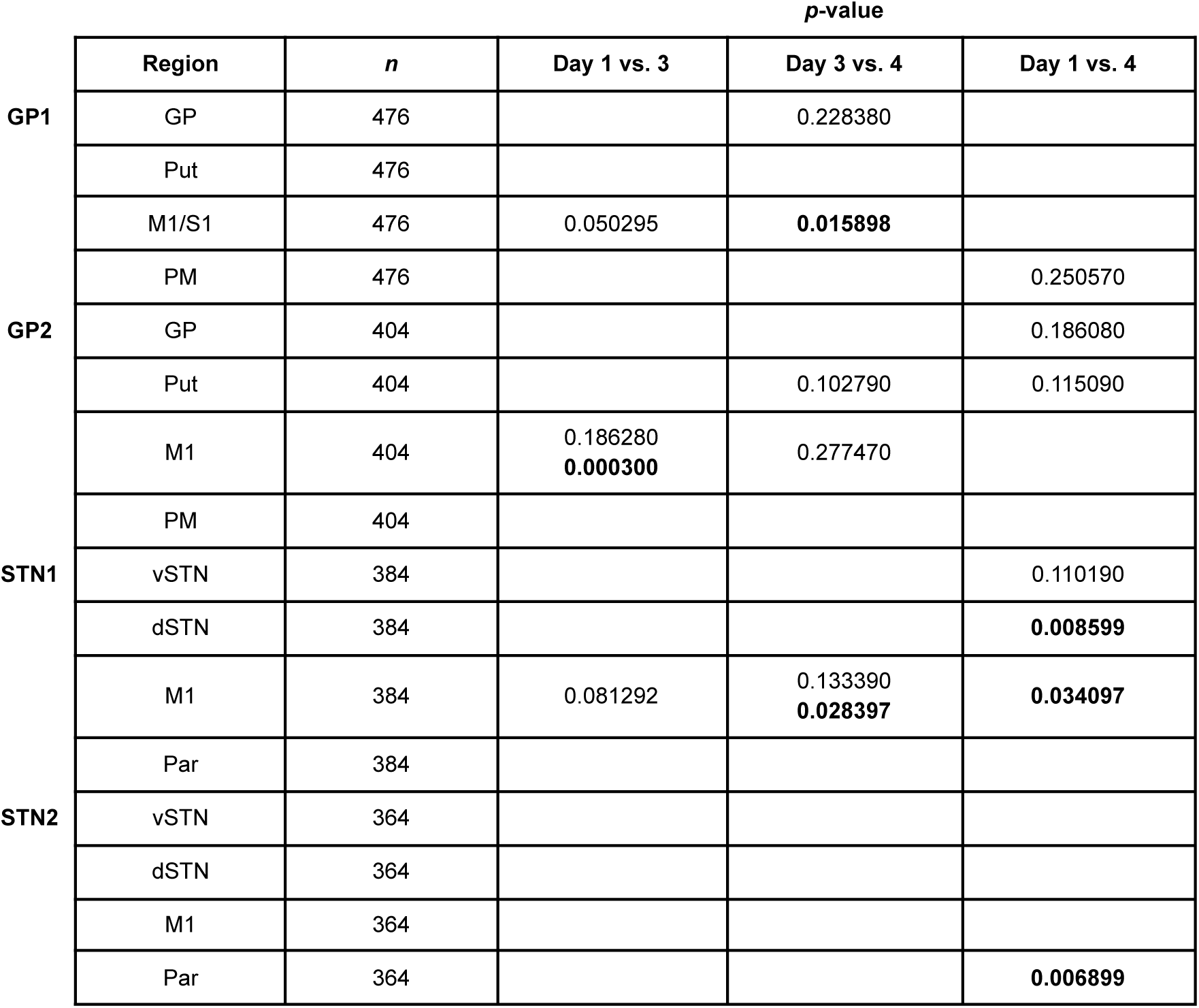
Across-day testing of intraregional δ-γ ^δ^ PPC: sequence learning-related *p*-values. Blank cells indicate no time regions passed initial thresholding. Multiple values in a single cell correspond to multiple time regions that passed initial thresholding.

|  |  | *p*-value |  |  |  |
| --- | --- | --- | --- | --- | --- |
|  | Region | *n* | Day 1 vs. 3 | Day 3 vs. 4 | Day 1 vs. 4 |
| **GP1** | GP | 476 |  | 0.228380 |  |
|  | Put | 476 |  |  |  |
|  | M1/S1 | 476 | 0.050295 | **0.015898** |  |
|  | PM | 476 |  |  | 0.250570 |
| **GP2** | GP | 404 |  |  | 0.186080 |
|  | Put | 404 |  | 0.102790 | 0.115090 |
|  | M1 | 404 | 0.186280<br>**0.000300** | 0.277470 |  |
|  | PM | 404 |  |  |  |
| **STN1** | vSTN | 384 |  |  | 0.110190 |
|  | dSTN | 384 |  |  | **0.008599** |
|  | M1 | 384 | 0.081292 | 0.133390<br>**0.028397** | **0.034097** |
|  | Par | 384 |  |  |  |
| **STN2** | vSTN | 364 |  |  |  |
|  | dSTN | 364 |  |  |  |
|  | M1 | 364 |  |  |  |
|  | Par | 364 |  |  | **0.006899** |

**Table 12.** Baseline testing of intraregional δ-γ ^δ^ PSI: sequence learning-related *p*-values. Blank cells indicate no time regions passed initial thresholding. Multiple values in a single cell correspond to multiple time regions that passed initial thresholding. Earlier time regions are listed first. NA, not applicable.

|  |  | *p*-value |  |  |  |
| --- | --- | --- | --- | --- | --- |
|  | Region | *n* | Day 1 | Day 3 | Day 4 |
| **GP1** | GP | 238 | NA | NA | NA |
|  | Put | 238 | NA | NA | NA |
|  | M1/S1 | 238 | NA | **0.001200** | **0.016998** |
|  | PM | 238 | NA | NA | NA |
| **GP2** | GP | 202 | **0.021698** | NA | NA |
|  | Put | 202 | NA | NA | NA |
|  | M1 | 202 | NA |  | NA |
|  | PM | 202 | NA | NA | NA |
| **STN1** | vSTN | 192 | NA | NA | NA |
|  | dSTN | 192 | NA | NA | NA |
|  | M1 | 192 | NA | NA | **0.031397** |
|  | Par | 192 | **0.008699**<br>**0.026997** | NA | NA |
| **STN2** | vSTN | 182 | NA | NA | NA |
|  | dSTN | 182 |  |  | NA |
|  | M1 | 182 |  | **0.000100** |  |
|  | Par | 182 | NA | **0.005599** | **0.017498** |

**Table 13.** Baseline testing of movement-related γ amplitude synchronization: *p*-values. Blank cells indicate no time regions passed initial thresholding. Multiple values in a single cell correspond to multiple time regions that passed initial thresholding. For each statistical test, 10,000 bootstrap samples of gamma amplitude were generated. The p-value was computed as percent of bootstrap sample means < 0.

| | | $p$-value | | | |
| --- | --- | --- | --- | --- | --- |
| | Region | $n$ | Day 1 | Day 3 | Day 4 |
| GP1 | GP | 238 | 0.884712 | 0.192081 | 0.956804 |
|  | Put | 238 | 0.558444 | 0.238276 | 0.788321 |
|  | M1/S1 | 238 | **0.000100** | **0.000100** | **0.000100** |
|  | PM | 238 | 0.120588 | **0.009799** | **0.000100** |
| GP2 | GP | 206 | 0.308669 | 0.762524 | 0.695830 |
|  | Put | 206 | 0.232177 | 0.936106 | **0.047995** |
|  | M1 | 206 | **0.000100** | **0.000900** | **0.000100** |
|  | PM | 206 | 0.522548 | 0.770323 | 0.067493 |
| STN1 | vSTN | 154 | 0.984202 | 0.202380 | 0.185681 |
|  | dSTN | 154 | 0.996300 | **0.021198** | 0.627437 |
|  | M1 | 154 | **0.006599** | **0.000100** | **0.001100** |
|  | Par | 154 | 0.853415 | **0.000100** | **0.001100** |
| STN2 | vSTN | 184 | 0.317568 | 0.297170 | 0.198980 |
|  | dSTN | 184 | 0.686131 | 0.794721 | 0.779622 |
|  | M1 | 184 | **0.000100** | **0.000100** | **0.000100** |
|  | Par | 184 | 0.967303 | 0.647535 | 0.988501 |

Such anatomically widespread δ phase alignment to cue in improvers suggests cross-area coordinated δ activity sufficient to facilitate coordinated recruitment of motor cortical and striatal ensembles with learning^29^. To assess this, we analyzed interregional δ phase coupling. Using pairwise phase consistency (PPC), we tested for an *increase*, relative to session-wide baseline, in undirected phase coherence aligned to cue and compared PPC between days^57^. When undirected coherence was significant, we compared the phase slope index (PSI; directed coherence) to chance (no channel leads)^58^. Two brain regions with a common input could show significant and stable undirected coherence even if a phase lead developed with learning. Undirected coherence exceeded baseline too infrequently to justify systematic between-day PSI testing, so we also noted as sequence practice- or task exposure-related any directed coherence that occurred only on specific days (Day 3 for sequence practice; Days 3 and 4 or Day 4 for task exposure).

Only improvers demonstrated cortically-led network δ synchrony (**Figure 6B**, **Figure 6D–E**). This synchrony increased above session-wide baseline in response to the cue and demonstrated sequence learning- and task exposure-related effects in improvers. In GP1, PM led M1/S1 and Put, and Put led GP, with small but significant M1/S1-striatal coherence associated with leads for M1/S1→GP and Put→M1/S1. GP led PM with task exposure. These effects were absent in GP2, for whom M1 instead led PM for familiar sequences. In STN1, M1 led dSTN for familiar sequences, and Par led dSTN and M1 with task exposure. Significant cue-related inter-STN PPC for familiar sequences indicates STN was recruited to network δ synchrony, partly by M1 and perhaps by an unrecorded region. In STN2, no directed phase leads occurred, and significant inter-STN PPC lacked the consistent local δ phase needed for event-locked coordinated phase coding (**Figure 6A**). Notably, in both nonimprovers, inter-BG δ synchrony started high and decreased with task exposure—an effect paralleled in session-wide inter-BG δ coherence, opposite that observed in improvers (**Figure 6C**). High inter-basal ganglia δ synchrony in nonimprovers was linked to a lack of cue-aligned local BG δ phase and an absence of cortico-basal ganglia δ synchrony.

### Sequence learning is associated with δ-γ coupling within and between cortex and basal ganglia

Having identified coordinated δ activity theoretically capable of supporting δ→spike coupling to facilitate the sequence-specific activity reflected in classification analysis, we next directly evaluated δ-γ coupling. We assessed both local and cross-area δ phase-high γ coupling. Using pairwise phase consistency and phase slope index, we calculated undirected and directed coherence between δ phase and the δ phase of the γ amplitude envelope (γ_h_^δ^). For interregional cross-frequency coupling (CFC), we report all results for which at least one subject demonstrates an effect of either sequence practice or task exposure.

Contrary to expectation, local δ→γ_h_^δ^ coupling was rare, suggesting that local δ→spike coupling may not have been a primary mechanism facilitating sequence-specific activity (**Supplementary Figure 10**). Practice-dependent γ_h_^δ^→δ coupling was more common, though also inconsistent across subjects. Nonetheless, M1 high γ amplitude still significantly increased in the RT period on all days in all subjects (**Supplementary Figure 11**), as expected for successful movement. This indicates that activity reflected in local δ→γ_h_^δ^ coupling was not a necessary step in general motor initiation for these subjects^44,67^.

Sequence learning was more associated with a range of interregional δ-γ_h_^δ^ effects (**Figure 7A–B**). These effects differed depending on the involved brain regions, which is consistent with the specialized roles of different regions in the cortico-basal ganglia network. In GP1 but not GP2, sequence practice was associated with premotor lead of M1/S1, as well as M1/S1 and PM lead of putamen. In STN1 but not STN2, sequence practice was associated with dSTN lead of M1 and vSTN, as well as M1 lead of Par. In STN2, Par instead led M1 on Days 1 and 3, but this effect diminished with task exposure. Otherwise, there were no task exposure-related effects in any subjects. Notably, δ led γ_h_^δ^ only for M1 γ_h_^δ^ with sequence learning in improvers, potentially reflecting underlying δ→spike coupling that led to the improvement-related increase in M1 γ’s predictive value (**Figure 5**).

**Figure 7.**
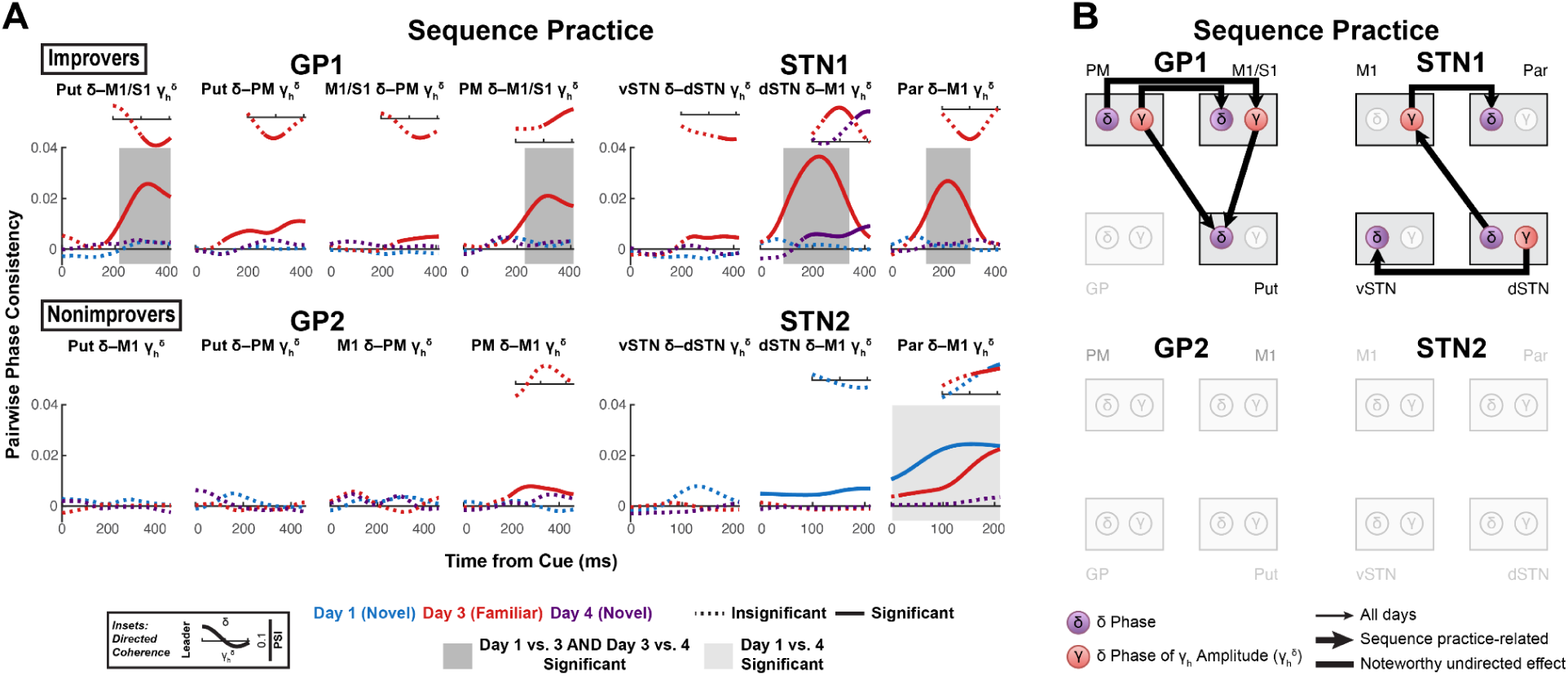
Sequence learning is associated with corticocortical, cortico-basal ganglia and inter-basal ganglia δ-γ couplings. (**A**) Sequence practice-related effects in interregional δ-γ ^δ^ coherence. (Large plots) Pairwise phase consistency (PPC, undirected measure) was calculated between δ phase and the δ phase of the high γ amplitude envelope. Solid line indicates significant PPC (*h_0_* = coherence is not higher than expected given the phase distribution, *α =* 0.05, one-sided, cluster-based permutation with 10,000 resamples. See **Table 14** for *p*-values.). Shaded box indicates significant difference in PPC between days (*α =* 0.05, two-sided, cluster-based permutation with 10,000 resamples. See **Table 15** for *p*-values.). (Insets) Phase slope index (PSI, directed measure) for significant PPC time series. Solid line indicates significant PSI (*h_0_* = no channel leads, *α =* 0.05, two-sided, cluster-based permutation with 10,000 resamples. See **Table 16** for *p*-values.). (**B**) Network diagrams illustrating sequence practice-related δ-γ ^δ^ effects.

**Table 14.** Baseline testing of interregional δ-γ ^δ^ PPC: sequence learning-related *p*-values. Blank cells indicate no time regions passed initial thresholding. Multiple values in a single cell correspond to multiple time regions that passed initial thresholding.

|  |  | *p*-value |  |  |  |
| --- | --- | --- | --- | --- | --- |
|  | Region | *n* | Day 1 | Day 3 | Day 4 |
| **GP1** | Put-M1/S1 | 238 |  | 0.194480<br>**0.002200** | 0.178580 |
|  | Put-PM | 238 |  | **0.000500** | 0.141290 |
|  | M1/S1-PM | 238 |  | **0.026197** | 0.305570 |
|  | PM-M1/S1 | 238 | 0.051495<br>0.309370 | **0.001500** | 0.086391 |
| **GP2** | Put-M1 | 202 |  |  |  |
|  | Put-PM | 202 | 0.137190 |  | 0.140690 |
|  | M1-PM | 202 | 0.312570 | 0.072593 | 0.199380<br>0.299170 |
|  | PM-M1 | 202 |  | **0.001500** | 0.103290 |
| **STN1** | vSTN-dSTN | 192 |  | **0.007399** | 0.110590 |
|  | dSTN-M1 | 192 | 0.333870 | **0.000100** | **0.002100** |
|  | Par-M1 | 192 | 0.151680 | **0.001700** |  |
| **STN2** | vSTN-dSTN | 182 |  |  |  |
|  | dSTN-M1 | 182 | **0.000100** |  |  |
|  | Par-M1 | 182 | **0.000100** | **0.002700** |  |

**Table 15.** Across-day testing of interregional δ-γ ^δ^ PPC: sequence learning-related *p*-values. Blank cells indicate no time regions passed initial thresholding. Multiple values in a single cell correspond to multiple time regions that passed initial thresholding.

|  |  | *p*-value |  |  |  |
| --- | --- | --- | --- | --- | --- |
|  | Region | *n* | Day 1 vs. 3 | Day 3 vs. 4 | Day 1 vs. 4 |
| **GP1** | Put-M1/S1 | 238 | **0.047695**<br>**0.002000** | 0.223680<br>**0.005400** | 0.192780 |
|  | Put-PM | 238 | 0.060894<br>0.051795 | 0.051095<br>0.152480 |  |
|  | M1/S1-PM | 238 | 0.050395 |  |  |
|  | PM-M1/S1 | 238 | **0.008599** | **0.007899** |  |
| **GP2** | Put-M1 | 202 |  |  |  |
|  | Put-PM | 202 | 0.267070 |  | 0.257170 |
|  | M1-PM | 202 |  |  |  |
|  | PM-M1 | 202 | 0.069393 |  |  |
| **STN1** | vSTN-dSTN | 192 | **0.003200** |  |  |
|  | dSTN-M1 | 192 | **0.001100** | **0.000800** | 0.076792<br>0.136990 |
|  | Par-M1 | 192 | **0.004600** | **0.011999** |  |
| **STN2** | vSTN-dSTN | 182 |  |  |  |
|  | dSTN-M1 | 182 |  |  |  |
|  | Par-M1 | 182 |  |  | **0.000100** |

**Table 16.** Baseline testing of interregional δ-γ ^δ^ PSI: sequence learning-related *p*-values. Blank cells indicate no time regions passed initial thresholding. Multiple values in a single cell correspond to multiple time regions that passed initial thresholding. Earlier time regions are listed first. NA, not applicable.

|  |  | *p*-value |  |  |  |
| --- | --- | --- | --- | --- | --- |
|  | Region | *n* | Day 1 | Day 3 | Day 4 |
| **GP1** | Put-M1/S1 | 238 | NA | **0.005399** | NA |
|  | Put-PM | 238 | NA | **0.025097** | NA |
|  | M1/S1-PM | 238 | NA | **0.031497** | NA |
|  | PM-M1/S1 | 238 | NA | **0.006699** | NA |
| **GP2** | Put-M1 | 202 | NA | NA | NA |
|  | Put-PM | 202 | NA | NA | NA |
|  | M1-PM | 202 | NA | NA | NA |
|  | PM-M1 | 202 | NA |  | NA |
| **STN1** | vSTN-dSTN | 192 | NA | 0.105989<br>**0.043797** | NA |
|  | dSTN-M1 | 192 | NA | **0.006399** | **0.047595** |
|  | Par-M1 | 192 | NA | **0.022698** | NA |
| **STN2** | vSTN-dSTN | 182 | NA | NA | NA |
|  | dSTN-M1 | 182 |  | NA | NA |
|  | Par-M1 | 182 | **0.048195** | **0.021898** | NA |

**Table 17.** Baseline testing of movement-related β amplitude desynchronization: *p*-values. Blank cells indicate no time regions passed initial thresholding. Multiple values in a single cell correspond to multiple time regions that passed initial thresholding.

|  |  | *p*-value |  |  |  |
| --- | --- | --- | --- | --- | --- |
|  | Region | *n* | Day 1 | Day 3 | Day 4 |
| **GP1** | GP | 238 | **0.000100** | **0.000100** | **0.000100** |
|  | Put | 238 | **0.000900** | **0.000100** | **0.000300** |
|  | M1/S1 | 238 | **0.000100** | **0.000100** | **0.000100** |
|  | PM | 238 | **0.000100** | **0.000100** | **0.000100** |
| **GP2** | GP | 206 | 0.518448 | **0.002800** | 0.839816 |
|  | Put | 206 | **0.000100** | **0.000300** | **0.001000** |
|  | M1 | 206 | **0.000100** | **0.000100** | **0.000100** |
|  | PM | 206 | **0.000100** | **0.000500** | **0.000100** |
| **STN1** | vSTN | 154 | 0.072893 | 0.128887 | 0.230977 |
|  | dSTN | 154 | **0.001900** | **0.017898** | **0.009299** |
|  | M1 | 154 | **0.000100** | **0.000100** | **0.000100** |
|  | Par | 154 | **0.004400** | **0.000100** | **0.000100** |
| **STN2** | vSTN | 184 | 0.878312 | 0.796820 | 0.958004 |
|  | dSTN | 184 | 0.200280 | 0.142286 | 0.298270 |
|  | M1 | 184 | **0.000600** | **0.003700** | **0.000100** |
|  | Par | 184 | 0.931507 | 0.510249 | 0.300370 |

**Table 18.** Baseline testing of intraregional δ-β^δ^ PPC: sequence learning-related *p*-values. Blank cells indicate no time regions passed initial thresholding. Multiple values in a single cell correspond to multiple time regions that passed initial thresholding

|  |  | *p*-value |  |  |  |
| --- | --- | --- | --- | --- | --- |
|  | Region | *n* | Day 1 | Day 3 | Day 4 |
| **GP1** | GP | 238 | 0.078792<br>**0.007699** | **0.002000** | **0.000100** |
|  | Put | 238 | 0.101590 | 0.137790 |  |
|  | M1/S1 | 238 | **0.008199**<br>0.122790 | **0.000100** | 0.082492 |
|  | PM | 238 | **0.011399** | **0.002400** | **0.006799** |
| **GP2** | GP | 202 | **0.013999** | 0.227380<br>0.175280 |  |
|  | Put | 202 | 0.066593 |  | **0.048995** |
|  | M1 | 202 |  | **0.000200** | **0.017298** |
|  | PM | 202 | **0.019398** | 0.060294 | 0.154080 |
| **STN1** | vSTN | 192 |  |  |  |
|  | dSTN | 192 |  |  | **0.015498** |
|  | M1 | 192 |  | **0.000500** | **0.001600** |
|  | Par | 192 |  |  | **0.002300** |
| **STN2** | vSTN | 182 |  |  |  |
|  | dSTN | 182 | **0.000100** | **0.027397** | **0.000100** |
|  | M1 | 182 | 0.134290<br>**0.015798** | 0.085591 | **0.004100** |
|  | Par | 182 |  |  |  |

**Table 19.** Across-day testing of intraregional δ-β^δ^ PPC: sequence learning-related *p*-values. Blank cells indicate no time regions passed initial thresholding. Multiple values in a single cell correspond to multiple time regions that passed initial thresholding.

|  |  | *p*-value |  |  |  |
| --- | --- | --- | --- | --- | --- |
|  | Region | *n* | Day 1 vs. 3 | Day 3 vs. 4 | Day 1 vs. 4 |
| **GP1** | GP | 476 |  | **0.045295** | **0.025497** |
|  | Put | 476 |  |  |  |
|  | M1/S1 | 476 | 0.153880 | **0.033597** | **0.009299** |
|  | PM | 476 | 0.212180 |  |  |
| **GP2** | GP | 404 |  |  |  |
|  | Put | 404 |  |  | 0.244980<br>**0.006199** |
|  | M1 | 404 | **0.003300** | **0.000400** | 0.239280<br>0.158480 |
|  | PM | 404 |  | 0.220780 |  |
| **STN1** | vSTN | 384 |  |  |  |
|  | dSTN | 384 |  | 0.193880 |  |
|  | M1 | 384 |  |  |  |
|  | Par | 384 |  |  | **0.005699** |
| **STN2** | vSTN | 364 |  |  |  |
|  | dSTN | 364 |  |  |  |
|  | M1 | 364 |  |  |  |
|  | Par | 364 |  |  |  |

**Table 20.**
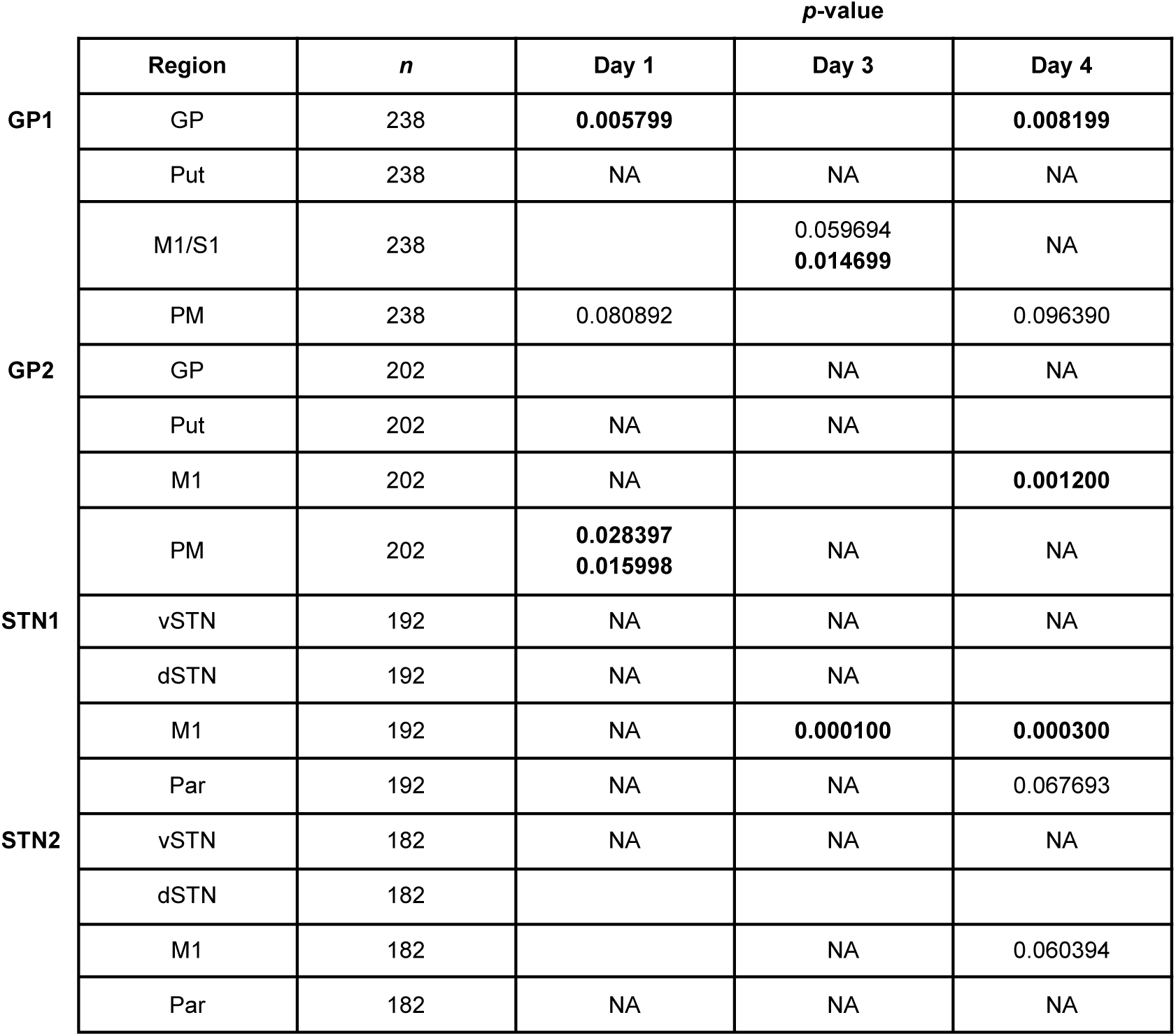
Baseline testing of intraregional δ-β^δ^ PSI: sequence learning-related *p*-values. Blank cells indicate no time regions passed initial thresholding. Multiple values in a single cell correspond to multiple time regions that passed initial thresholding. Earlier time regions are listed first. NA, not applicable.

|  |  | *p*-value |  |  |  |
| --- | --- | --- | --- | --- | --- |
|  | Region | *n* | Day 1 | Day 3 | Day 4 |
| **GP1** | GP | 238 | **0.005799** |  | **0.008199** |
|  | Put | 238 | NA | NA | NA |
|  | M1/S1 | 238 |  | 0.059694<br>**0.014699** | NA |
|  | PM | 238 | 0.080892 |  | 0.096390 |
| **GP2** | GP | 202 |  | NA | NA |
|  | Put | 202 | NA | NA |  |
|  | M1 | 202 | NA |  | **0.001200** |
|  | PM | 202 | **0.028397**<br>**0.015998** | NA | NA |
| **STN1** | vSTN | 192 | NA | NA | NA |
|  | dSTN | 192 | NA | NA |  |
|  | M1 | 192 | NA | **0.000100** | **0.000300** |
|  | Par | 192 | NA | NA | 0.067693 |
| **STN2** | vSTN | 182 | NA | NA | NA |
|  | dSTN | 182 |  |  |  |
|  | M1 | 182 |  | NA | 0.060394 |
|  | Par | 182 | NA | NA | NA |

### Network β does not gate motor cortical δ

Using the same approach, we finally tested whether network β gated motor cortical excitability by assessing δ-β coupling. We calculated coupling between δ phase and the δ phase of the β amplitude envelope (β^δ^). Cortical excitability reflected as a deflection in cortical δ, could be led by cortical β, subcortical β or a combination of both. Thus, we assessed both intraregional and interregional δ-β^δ^ coupling.

There was substantial overlap of regions with consistent movement-related β desynchronization and regions with cue-related δ phase alignment (all motor regions in improvers; motor cortex and putamen in nonimprovers) (**Figure 6A**, **Supplementary Figure 12**), suggesting widespread local δ-β coupling. However, significant local δ-β^δ^ directed coupling was rare, never occurred across all days and demonstrated inconsistent direction of phase lead in M1 across subjects (**Supplementary Figure 13**). Local β gating of low-frequency shifts in M1 excitability was not likely a general mechanism in motor initiation.

Likewise, there was no evidence for consistent interregional β gating of M1 excitability; however, striatocortical δ→β^δ^ coupling developed with sequence learning in GP1 (**Figure 8A–B**). This contrasts with sequence practice-related corticostriatal δ-β^δ^ coupling in GP2. With task exposure, primary motor cortex led PM in both GP subjects, but only in GP1 was GP led by all recorded brain regions (Put, M1/S1, PM) (**Figure 8C–D**). In STN subjects, no sequence practice-related effects were observed. However, with task exposure in STN1, Par led M1 and dSTN—effects absent in STN2. These results associate sequence learning and task exposure with unique, anatomically-specific patterns of δ-β coupling in improvers.

**Figure 8.**
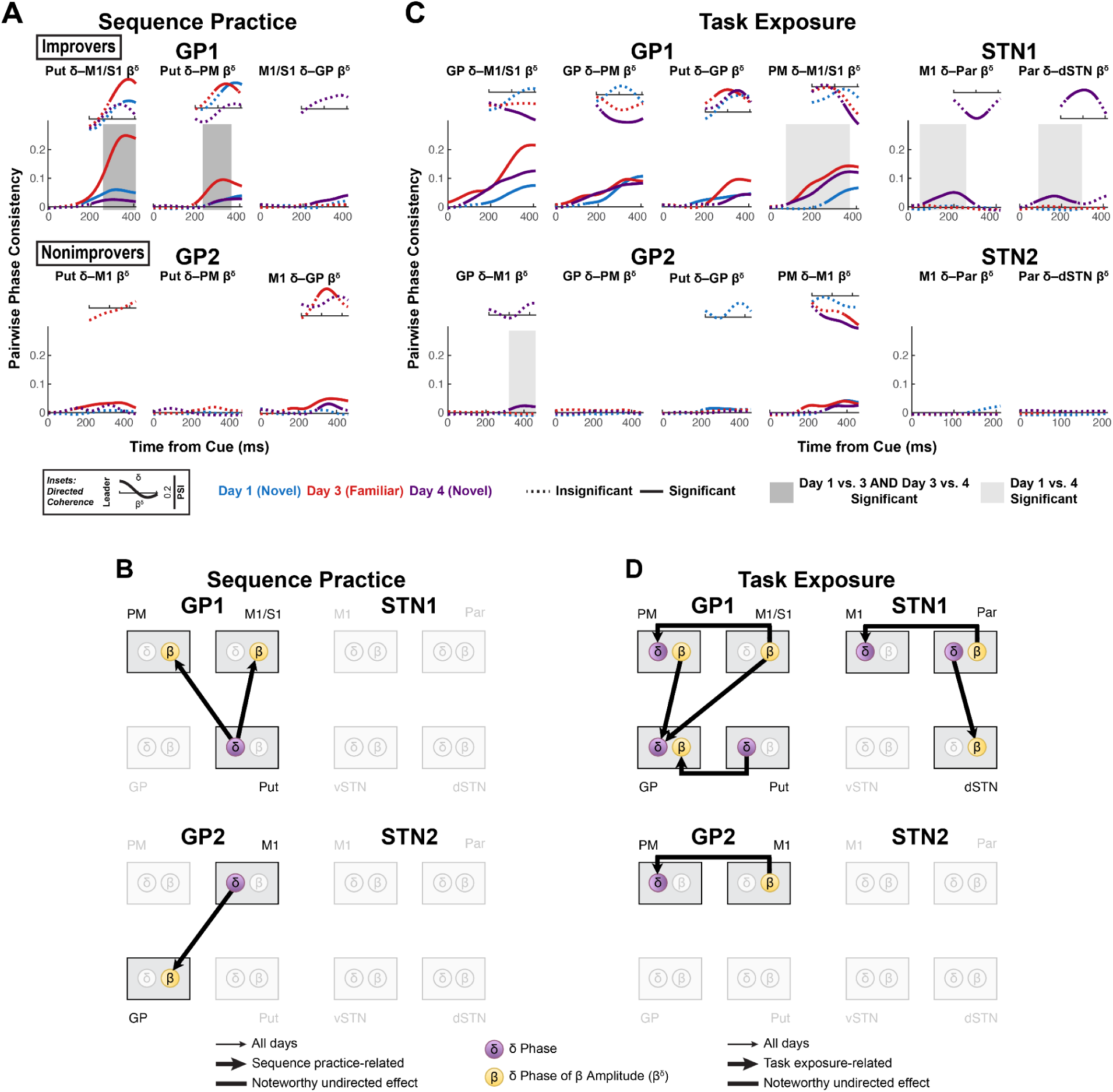
Striatocortical δ-β coupling increases with sequence learning, while task exposure brings a range of δ-β couplings in improvers mostly absent in nonimprovers. (**A**) Sequence practice-related effects in interregional δ-β^δ^ coherence. (Large plots) Pairwise phase consistency (PPC, undirected measure) was calculated between δ phase and the δ phase of the β amplitude envelope. Solid line indicates significant PPC (*h_0_* = coherence is not higher than expected given the phase distribution, *α =* 0.05, one-sided, cluster-based permutation with 10,000 resamples. See **Table 21** for *p*-values.). Shaded box indicates significant difference in PPC between days (*α =* 0.05, two-sided, cluster-based permutation with 10,000 resamples. See **Table 22** for *p*-values.). (Insets) Phase slope index (PSI, directed measure) for significant PPC time series. Solid line indicates significant PSI (*h_0_* = no channel leads, *α =* 0.05, two-sided, cluster-based permutation with 10,000 resamples. See **Table 23** for *p*-values.). (**B**) Network diagrams illustrating sequence practice-related δ-β^δ^ effects. (**C**) Task exposure-related effects in interregional δ-β^δ^ coherence. Visualization and statistics identical to (**A**) (See **Tables 24–25** for *p*-values.). (**D**) Network diagrams illustrating task exposure-related δ-β^δ^ effects.

**Table 21.** Baseline testing of interregional δ-β^δ^ PPC: sequence learning-related *p*-values. Blank cells indicate no time regions passed initial thresholding. Multiple values in a single cell correspond to multiple time regions that passed initial thresholding.

|  |  | *p*-value |  |  |  |
| --- | --- | --- | --- | --- | --- |
|  | Region | *n* | Day 1 | Day 3 | Day 4 |
| **GP1** | Put-M1/S1 | 238 | **0.002300** | **0.001100** | **0.005799** |
|  | Put-PM | 238 | **0.025697** | **0.003700** | **0.027197** |
|  | M1/S1-GP | 238 | 0.108990 |  | **0.013499** |
| **GP2** | Put-M1 | 202 |  | **0.000500** | 0.137590<br>0.066093 |
|  | Put-PM | 202 |  | 0.075092 | 0.148190 |
|  | M1-GP | 202 | 0.238080 | **0.000500** | **0.042296** |

**Table 22.** Across-day testing of interregional δ-β^δ^ PPC: sequence learning-related *p*-values. Blank cells indicate no time regions passed initial thresholding. Multiple values in a single cell correspond to multiple time regions that passed initial thresholding.

|  |  | *p*-value |  |  |  |
| --- | --- | --- | --- | --- | --- |
|  | Region | *n* | Day 1 vs. 3 | Day 3 vs. 4 | Day 1 vs. 4 |
| **GP1** | Put-M1/S1 | 238 | **0.016398** | **0.005500** |  |
|  | Put-PM | 238 | **0.028997** | **0.006799** |  |
|  | M1/S1-GP | 238 |  |  |  |
| **GP2** | Put-M1 | 202 | 0.175780 |  | 0.129790 |
|  | Put-PM | 202 |  | **0.033297** |  |
|  | M1-GP | 202 | 0.118190 |  |  |

**Table 23.**
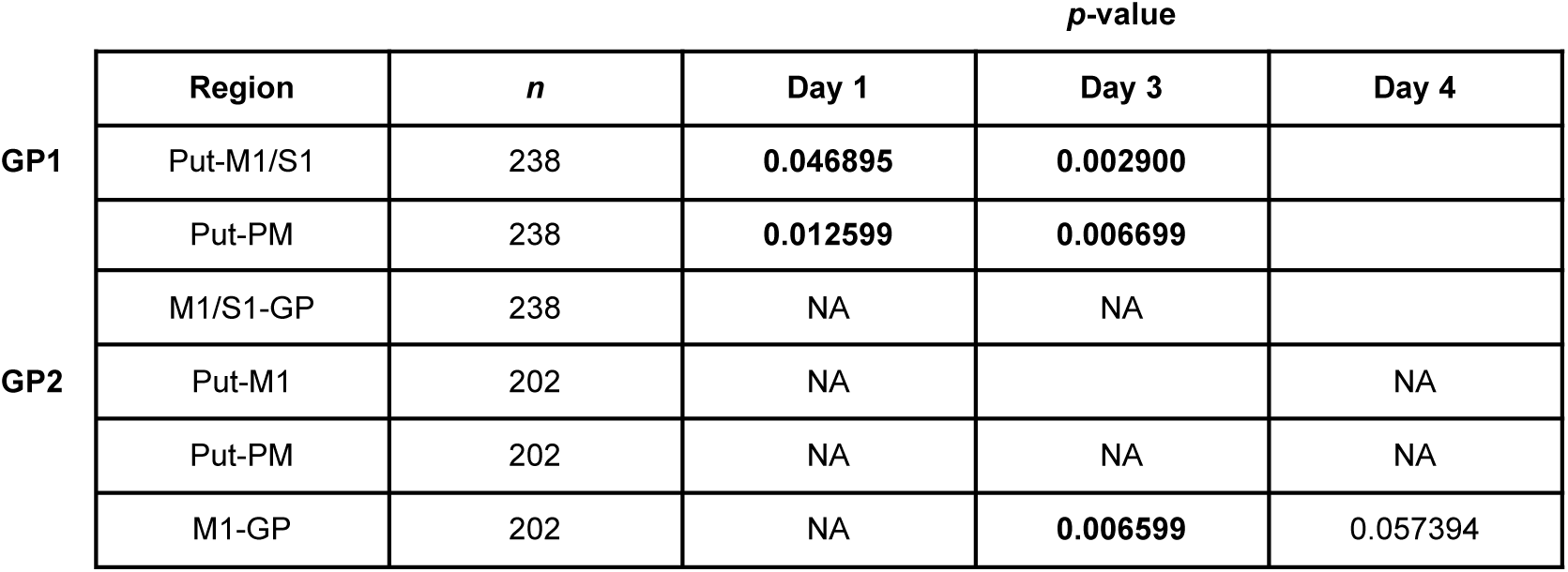
Baseline testing of interregional δ-β^δ^ PSI: sequence learning-related *p*-values. Blank cells indicate no time regions passed initial thresholding. Multiple values in a single cell correspond to multiple time regions that passed initial thresholding. Earlier time regions are listed first. NA, not applicable.

|  |  | *p*-value |  |  |  |
| --- | --- | --- | --- | --- | --- |
|  | Region | *n* | Day 1 | Day 3 | Day 4 |
| GP1 | Put-M1/S1 | 238 | **0.046895** | **0.002900** |  |
|  | Put-PM | 238 | **0.012599** | **0.006699** |  |
|  | M1/S1-GP | 238 | NA | NA |  |
| GP2 | Put-M1 | 202 | NA |  | NA |
|  | Put-PM | 202 | NA | NA | NA |
|  | M1-GP | 202 | NA | **0.006599** | 0.057394 |

**Table 24.** Baseline testing of interregional δ-β^δ^ PPC: task exposure-related *p*-values. Blank cells indicate no time regions passed initial thresholding. Multiple values in a single cell correspond to multiple time regions that passed initial thresholding.

|  |  | *p*-value |  |  |  |
| --- | --- | --- | --- | --- | --- |
|  | Region | *n* | Day 1 | Day 3 | Day 4 |
| **GP1** | GP-M1/S1 | 238 | **0.002600** | **0.000100** | **0.000400** |
|  | GP-PM | 238 | **0.001800** | **0.000100** | **0.000300** |
|  | Put-GP | 238 | **0.001400** | **0.004600** | **0.024698** |
|  | PM-M1/S1 | 238 | **0.020798** | **0.000300** | **0.000200** |
| **GP2** | GP-M1 | 202 |  |  | **0.043396** |
|  | GP-PM | 202 |  |  |  |
|  | Put-GP | 202 | **0.031797**<br>0.236980<br>0.231280 |  |  |
|  | PM-M1 | 202 | **0.018498** | **0.001000** | **0.020098** |
| **STN1** | M1-Par | 192 |  |  | **0.002600** |
|  | Par-dSTN | 192 |  |  | **0.006799**<br>0.100790 |
| **STN2** | M1-Par | 182 | 0.110690 |  |  |
|  | Par-dSTN | 182 |  |  |  |

**Table 25.** Across-day testing of interregional δ-β^δ^ PPC: task exposure-related *p*-values. Blank cells indicate no time regions passed initial thresholding. Multiple values in a single cell correspond to multiple time regions that passed initial thresholding.

|  | p-value |  |  |  |  |
| --- | --- | --- | --- | --- | --- |
|  | Region | n | Day 1 vs. 3 | Day 3 vs. 4 | Day 1 vs. 4 |
| GP1 | GP-M1/S1 | 238 | 0.036996<br>0.004900 | 0.137690 | 0.110190 |
|  | GP-PM | 238 |  |  |  |
|  | Put-GP | 238 |  | 0.131690 |  |
|  | PM-M1/S1 | 238 | 0.000200 |  | 0.000400 |
| GP2 | GP-M1 | 202 |  |  | 0.010399 |
|  | GP-PM | 202 | 0.240780 |  | 0.215780 |
|  | Put-GP | 202 |  |  |  |
|  | PM-M1 | 202 | 0.192780 |  |  |
| STN1 | M1-Par | 192 |  | 0.058994 | 0.004100 |
|  | Par-dSTN | 192 |  | 0.130390<br>0.156380 | 0.004600<br>0.140990 |
| STN2 | M1-Par | 182 | 0.157680 |  | 0.135390 |
|  | Par-dSTN | 182 |  |  |  |

**Table 26.** Baseline testing of interregional δ-β^δ^ PSI: task exposure-related *p*-values. Blank cells indicate no time regions passed initial thresholding. Multiple values in a single cell correspond to multiple time regions that passed initial thresholding. Earlier time regions are listed first. NA, not applicable.

|  |  | *p*-value |  |  |  |
| --- | --- | --- | --- | --- | --- |
|  | Region | *n* | Day 1 | Day 3 | Day 4 |
| **GP1** | GP-M1/S1 | 238 | 0.079892 |  | **0.002200** |
|  | GP-PM | 238 |  |  | **0.000400** |
|  | Put-GP | 238 |  | **0.005699** | **0.018698** |
|  | PM-M1/S1 | 238 |  | 0.057894 | **0.028197** |
| **GP2** | GP-M1 | 202 | NA | NA |  |
|  | GP-PM | 202 | NA | NA | NA |
|  | Put-GP | 202 |  | NA | NA |
|  | PM-M1 | 202 |  | **0.028497** | **0.000300** |
| **STN1** | M1-Par | 192 | NA | NA | **0.006099** |
|  | Par-dSTN | 192 | NA | NA | **0.001500** |
| **STN2** | M1-Par | 182 | NA | NA | NA |
|  | Par-dSTN | 182 | NA | NA | NA |

## DISCUSSION

This study is the first invasive electrophysiological investigation of human cortico-basal ganglia dynamics in motor sequence learning. Though all subjects demonstrated M1 β desynchronization, δ phase alignment to cue and γ synchronization, a consistent β→δ→γ cascade was absent—possibly reflecting PD-related neuropathophysiology (**Figures 9–10**). Instead, δ-β and δ-γ coupling emerged with learning, possibly supporting optimization of preparatory activity. However, consistent with our predictions, we observed increasingly predictive sequence-specific cortical and basal ganglia γ activity alongside cue-aligned cortico-BG δ synchrony in improvers. Strikingly, all brain regions were eventually recruited, producing network δ synchrony. Furthermore, M1 γ’s increasing predictive value was linked to its coupling with synchronized network δ, tying coordinated network δ to enhanced recruitment of sequence-specific neural activity. In contrast, in nonimprovers, decreasingly predictive γ corresponded with minimal CFC and absent coordinated network δ activity. Highly coherent inter-BG δ did not synchronize with cortex or align to visual cues, suggesting it was pathological. These results suggest that both cortex and basal ganglia supported multi-element preplanning and that cortico-basal ganglia communication was critical to learning-driven optimization of motor preparation. This highlights how motor learning can harness a hierarchical functional network architecture to optimize information transfer and temporally structure neural activity in PD.

**Figure 9.**
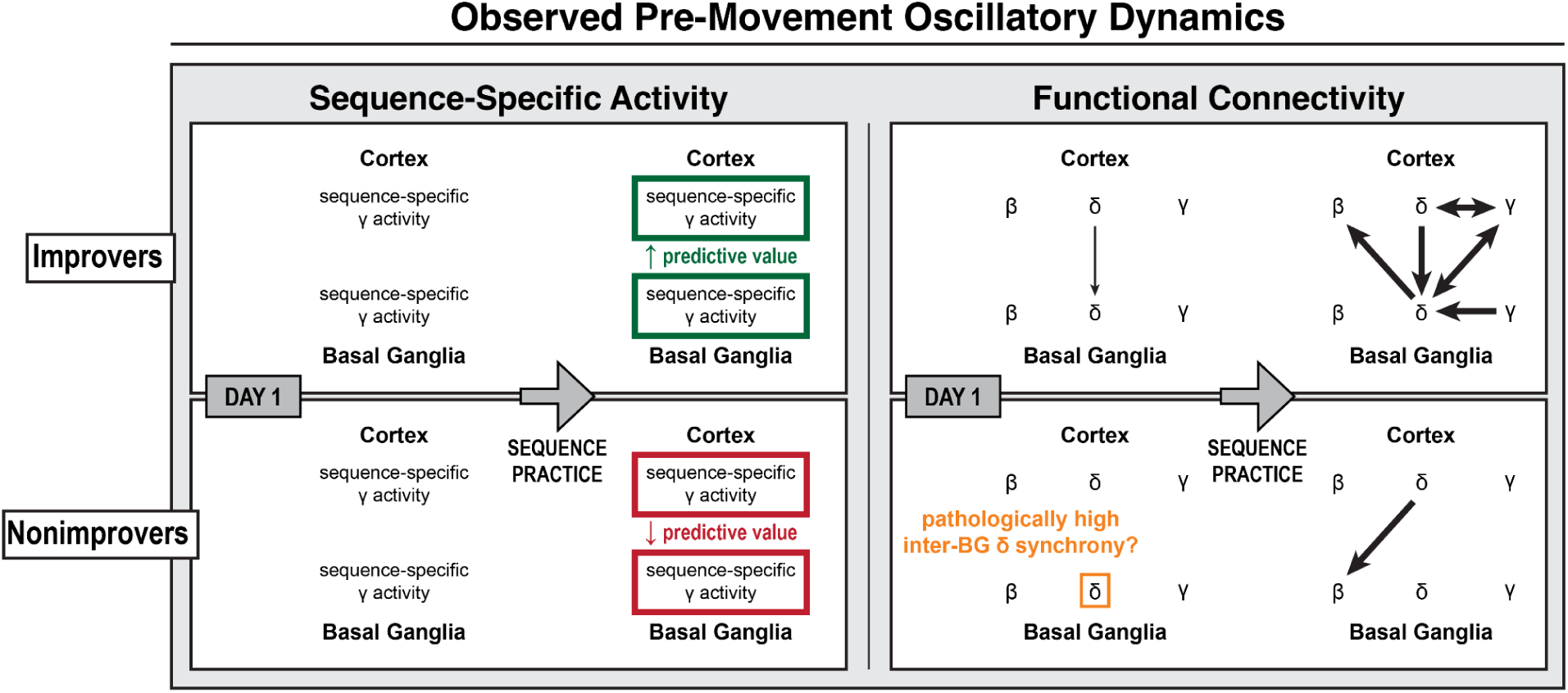
Observed learning-related cortico-basal ganglia activity during motor sequence initiation. Diagram of observed learning-related changes in sequence-specific activity and functional connectivity prior to the onset of motor sequences over multiple days of practice. (Left) Sequence-specific γ activity was present in all brain regions for all subjects. With practice, γ’s predictive value increased in improvers but decreased in nonimprovers. (Right) As predicted, successful sequence learning involved cortico-basal ganglia δ synchrony and δ→γ coupling with motor cortex γ, though δ→γ coupling was not observed with basal ganglia γ. Sequence learning also involved other cross-frequency couplings, most notably δ→γ and δ→β couplings in which basal ganglia led cortex. In nonimprovers, highly coherent inter-basal ganglia δ was uncoupled from cortex and unresponsive to task events. These results suggest 1) that sequence learning harnessed cortically-led δ phase coordination to organize distributed higher-frequency neural activity for the optimization of multi-element preplanning and 2) that pathological BG δ synchrony may interfere with BG δ phase coding by reducing BG sensitivity to cortical input, ultimately disrupting motor learning.

**Figure 10.**
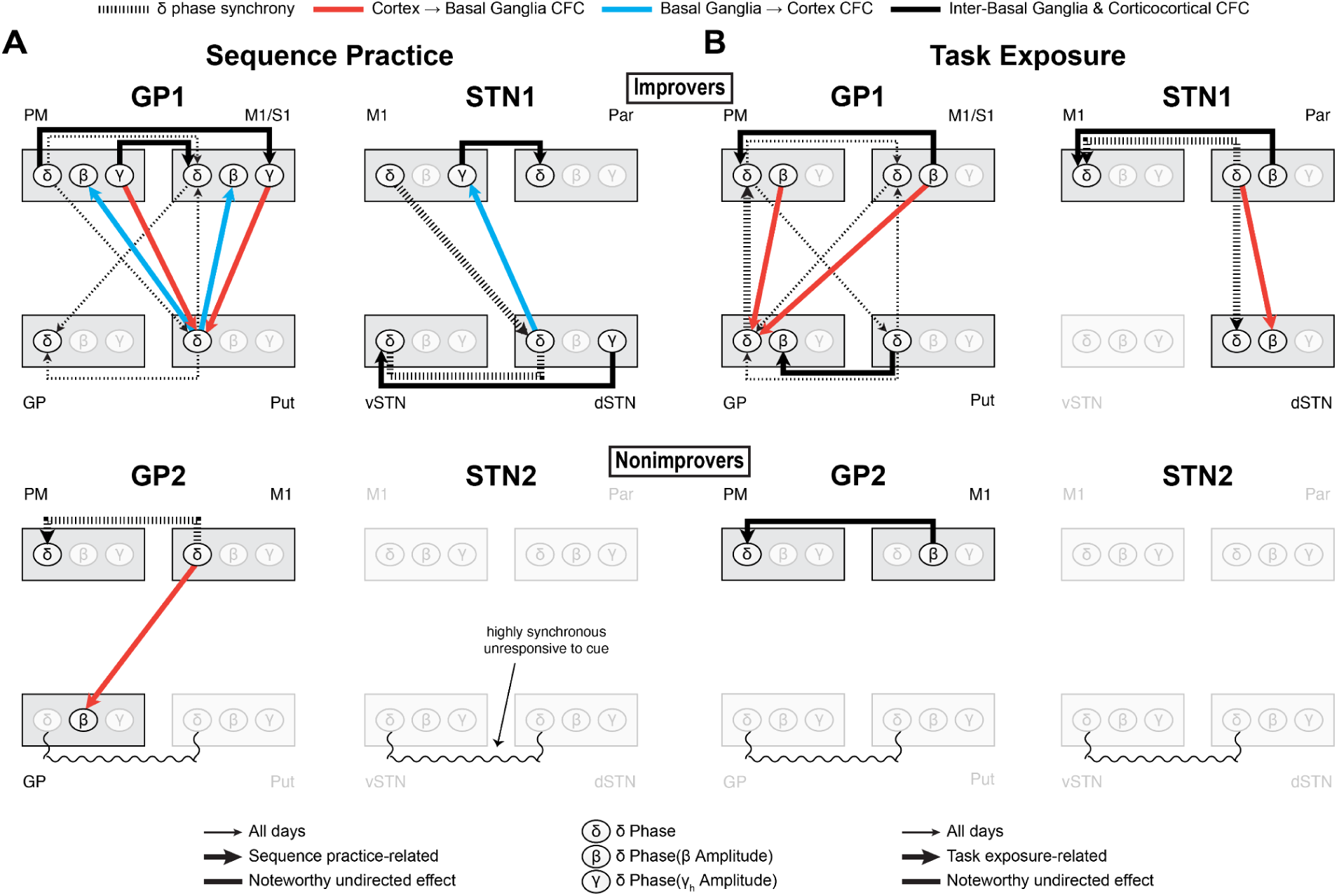
Coordinated δ activity supports a hierarchical functional network during sequence practice and task exposure that reflects the specialized roles of cortico-basal ganglia regions. Composite diagrams of interregional effects associated with (**A**) sequence practice and (**B**) task exposure. As motor sequence learning overlaps with increasing familiarity with the experimental process, we dissociated the functional connectivity effects related to sequence practice from those related to task exposure. In improvers, sequence learning and task exposure produced distinct patterns of CFC and changes to network δ dynamics, though sequence learning and task exposure were nonetheless linked by a common framework of cortically-led network δ synchrony. For each type of communication, the presence and direction of coupling depended on the involved brain regions, indicative of their specialized roles. In nonimprovers, the absence of both sequence practice- and task exposure-related effects involving subcortical δ further suggests that a locus of their interdependence may be cortico-basal ganglia δ synchrony. Furthermore, in all subjects, only the δ in regions that aligned to cue participated in interregional CFC. Except for motor cortex δ, this local δ alignment to cue was coupled with network δ synchrony. These findings support a model wherein a network capable of learning leverages preparatory coordinated network δ activity to organize both sequence learning- and task exposure-related effects, with the two effect types distinguished by a unique interplay of δ synchrony and CFC. This ultimately poses coordinated network δ as an ideal substrate for task familiarization to boost future sequence learning. Neurostimulation that enhances event-related cortico-basal ganglia δ coupling may thus prove a broadly effective strategy to improve skilled fine motor control in PD.

### The β→δ→γ framework for motor initiation: an assessment in Parkinson’s Disease

The expected cascade of network β→cortical δ→cortical γ for general motor initiation was absent in our subjects, indicating it was not prerequisite^29,41,44,50,51,67–74^. Our observation of both lead directions for δ-γ and δ-β couplings is novel, as most studies have employed undirected coupling measures^75–78^. Coupling in which β or γ led δ was often associated with enhanced δ alignment to cue or lead of other brain regions, suggesting a practice-driven role for β- and γ-related neural activity in influencing network δ dynamics. This may reflect compensatory or pathological mechanisms related to increased cortical δ sometimes observed with PD progression^79–81^.

### Practice-driven pre-movement sequence-specific activity

Pre-movement sequence-specific activity has been observed in nonhuman primate motor cortex, and we provide the first electrophysiological evidence of it in human cortex and basal ganglia^31,32^. Among canonical frequency bands, γ’s large bandwidth necessitated the most narrowband filters for time-frequency analysis, which inherently increased its representation in the total feature set. However, lasso regularization and within-time-frequency percentile-based feature selection helped prioritize features based on sparsity and discriminative value. Thus, sequence-specific γ activity may have arisen from unique neural ensembles, with distinct firing patterns and spatial proximities to the recording contacts resulting in differing field potential dynamics^53,82^.

While animal model motor learning studies suggest that increasingly predictive cortical and BG γ should couple with network δ, we did not observe δ→γ for BG γ^24–29,52^. Unrecorded regions may have facilitated BG γ’s increased predictive value. Alternatively, δ may have indirectly influenced sequence-specific activity through its lead of β, which may bind action plan-specific ensembles across the network^29^. Decreased decoding accuracy in some brain regions nonimprovers may have reflected reduced consistency of ensemble activity, potentially contributing to lack of motor improvement. Given our small sample size and recordings at the level of neural populations rather than individual neurons, these suggestions are conjecture and should inspire future experimental work.

### Sequence learning and task exposure produced distinct patterns of cortico-basal ganglia functional connectivity in improvers

Different patterns of both CFC and δ synchrony emerged with sequence learning and task exposure in improvers. For CFC, most notably, basal ganglia→cortex CFC developed only with sequence learning, with dSTN and putamen δ leading β or γ in cortex. This aligns with models implicating dSTN and putamen in action selection and learning and suggests that they perform these roles through CFC patterning of high-frequency cortical activity^1,3,83–88^. For δ synchrony, consistent with neuroimaging, we found corticostriatal δ coherence in GP1^89–91^. We further established a cortex→striatum direction of lead and found sequence learning-related M1→dSTN connectivity. In contrast, task exposure drove striatocortical δ synchrony. GP→PM δ synchrony completed a possible loop of cortico-basal ganglia δ phase coordination, via PM→Put→GP→PM. This suggests that general motor familiarization or the optimization of attentional processes could involve a positive feedback loop of network δ synchrony, which could aid future sequence learning^75,92–94^.

### Few effects of sequence practice and task exposure in nonimprovers: possible impact of pathologically synchronized basal ganglia δ

The absence of cue-related functional connectivity involving BG δ in nonimprovers may have been related to high session-wide inter-BG δ synchrony. Human and animal studies have detected δ-range spiking activity in GP and STN in low dopamine states and linked it to motor symptoms, suggesting that ineffective or absent dopamine replacement therapy can permit pathological BG δ activity^95–97^. Consistent with this, our subjects exhibiting elevated BG δ synchrony were those for whom general motor function remained the most compromised while on dopamine medication (**Table 1**). However, their daily post-task UPDRS upper limb scores were comparable to those of other subjects and demonstrated no apparent across-day drop alongside the across-day decrease in BG δ synchrony (**Table 3**). It is possible that task exposure drove the across-day BG δ synchrony decrease, promoting the possibility of future learning, and that the MDS-UPDRS upper limb component was too insensitive to detect associated changes in hallmark upper limb motor symptoms^98^. In any case, the concurrent lack of improvement and absence of practice-related effects involving BG δ suggests that event-related coordination of BG δ may have been important to motor learning in these subjects.

### Coordinated network δ as a facilitatory network state for learning-dependent cross-frequency coupling during sequence initiation

From the theoretical standpoint, δ is an ideal substrate for information multiplexing, local gain modulation and information transfer between distant phase-aligned brain regions^33,35,92–94^. Experimental evidence supports this, showing a role for δ in information sequencing through phase-specific ensemble patterning; sensory integration and attentional control through gain modulation achieved by changes in neural excitability; and functional network organization, information transfer and distributed representation through cross-area coordinated activity that coactivates distributed ensembles^24,33,35,36,38,52,77,99,100^.

Consistent with these roles, in our study, cross-area coordination of δ phase may have formed the infrastructure for a cue-responsive network state, within which learning-related δ-β and δ-γ coupling developed. While δ synchrony and phase alignment to cue occurred on Day 1 and increased across days, interregional δ-β and δ-γ directed couplings were initially mostly absent. Moreover, only the δ in brain regions that aligned to cue participated in interregional CFC, and, outside of motor cortex, local δ alignment to cue was always linked to network δ synchrony. These results highlight a potential role for cross-area δ synchrony in driving consistent event-related local δ phase dynamics, which can, in turn, support phase coding that patterns high-frequency activity.

This link between motor performance-related δ activity and CFC is consistent with lesion experiments in animal models and observational studies of the human cortical grasp network^24,52,77^. Other work has found reaction time-correlated cue-responsive δ phase reset in human hippocampus; reaction time-correlated δ phase in rat motor thalamus coupled to local spiking; motor learning-related M1-cerebellar δ synchrony in rats with enhanced cross-area spiking activity and even evidence for prefrontal guidance of motor plans via δ-β coupling in humans^101–104^. Coordinated δ may ultimately reflect a motor system-wide neural process facilitating local and distributed activity patterns over the course of learning.

However, the neurophysiological basis of the activity reflected in δ remains unclear. While δ has been associated with excitability and with single-unit spiking at δ frequency, it has also, similar to γ, been linked to the dynamics of population spiking activity^78,105^, and it could reflect synaptic input^106–109^. It is thus important to clarify that δ activity may not directly influence other neural activity, per sé, but rather reflect the underlying activity that drives the observed network effects.

### Clinical implications

Parkinson’s Disease often involves an impairment of action sequencing, but its neural basis is poorly understood and incompletely remediated by dopamine replacement therapy and conventional DBS^3,15,16^. Our observation that both sequence learning and task exposure involved cortico-basal ganglia δ phase coordination in improvers suggests that enhancing event-related cortico-basal ganglia δ synchrony may improve not only the learning of specific sequences but also the ability to adapt to new cued motor tasks. Improved task learning could, in turn, boost future sequence learning for a range of sequences or contexts, suggesting therapies aimed at restoring cortico-basal ganglia δ synchrony could be broadly effective for skilled hand movements. Electrically stimulating basal ganglia upon detection of motor intention and based on the ongoing phase of BG or cortical δ could promote cortico-basal ganglia δ synchrony and possibly disrupt pathological BG δ. Thus, our findings posit pre-movement δ phase-specific striatal or subthalamic DBS as a therapeutic neuromodulatory strategy to restore fine motor control.

### Central limitations

Our study was limited by a small sample size, as is common in human invasive electrophysiology. Anatomically nonoverlapping recordings in GP and STN subjects revealed the specialized neural activity of different brain regions but left unverifiable the consistency of effects across improvers or across nonimprovers. The lack of improvement in half the subjects allowed binary behavioral stratification but limited analysis to within-subjects comparisons with *n* = 1 per learner type per brain region. Future work should replicate this study in larger cohorts.

It is possible that performance floor effects led to incorrect behavioral stratification. Despite lack of performance improvement, nonimprovers typed at least as quickly as improvers. This indicates that lack of improvement was not due to difficulty manipulating the keyboard and could suggest that nonimprovers found the task too easy at baseline to show measurable improvement with sequence learning. However, performance decrements when switching between sequences within a given day—indicative of interference caused by sequence learning—were not apparent in nonimprovers’ single-trial performance data. It is possible that initial typing speeds reflected typical inter-individual variability.

### Conclusion

In individuals experiencing Parkinson’s Disease, we outline a hierarchical, learning-dependent functional architecture of oscillatory cortico-basal ganglia activity for the initiation of fine motor sequences. The findings illuminate how disparities in information content and flow may relate to disparities in motor learning outcomes. Extending this work in larger cohorts of individuals with PD could help elucidate the relationship between clinical characteristics and practice-related neural dynamics. This would further clarify the potential for phase-specific basal ganglia stimulation to modulate pathological neural dynamics in the production of fine motor skills.

## AUTHOR CONTRIBUTIONS

K.N.P. developed the experimental setup, including designing and fabricating custom behavioral hardware, programming firmware and developing protocols for neural-behavioral recording and alignment. K.N.P. performed the experiment. K.N.P. collected sleep journals and administered daily upper limb component of the MDS-UPDRS. K.N.P. performed all data curation, processing and analyses. K.N.P. and D.D.W. designed the experiment. K.N.P., R.A. and T.A.W. developed quantitative methodology. K.N.P. and K.H.L. set visual stimulus presentation. S.L. scored the UPDRS videos. D.D.W. and P.A.S. performed the neurosurgeries and funded the parent clinical trials. T.A.W. performed lead reconstruction. K.N.P. wrote the manuscript. All authors reviewed the manuscript.

## ACKNOWLEDGEMENTS

We thank the members of the OpenMind Consortium and the D.D.W., R.A., P.A.S. and S.L. laboratories for valuable discussion over the course of the study. We thank Rich Ivry for helpful guidance during experimental design and data interpretation, as well as critique of the manuscript. We thank Karunesh Ganguly for thoughtful advice during data interpretation and critique of the manuscript.

## SUPPLEMENTARY FIGURES

**Supplementary Figure 1.**
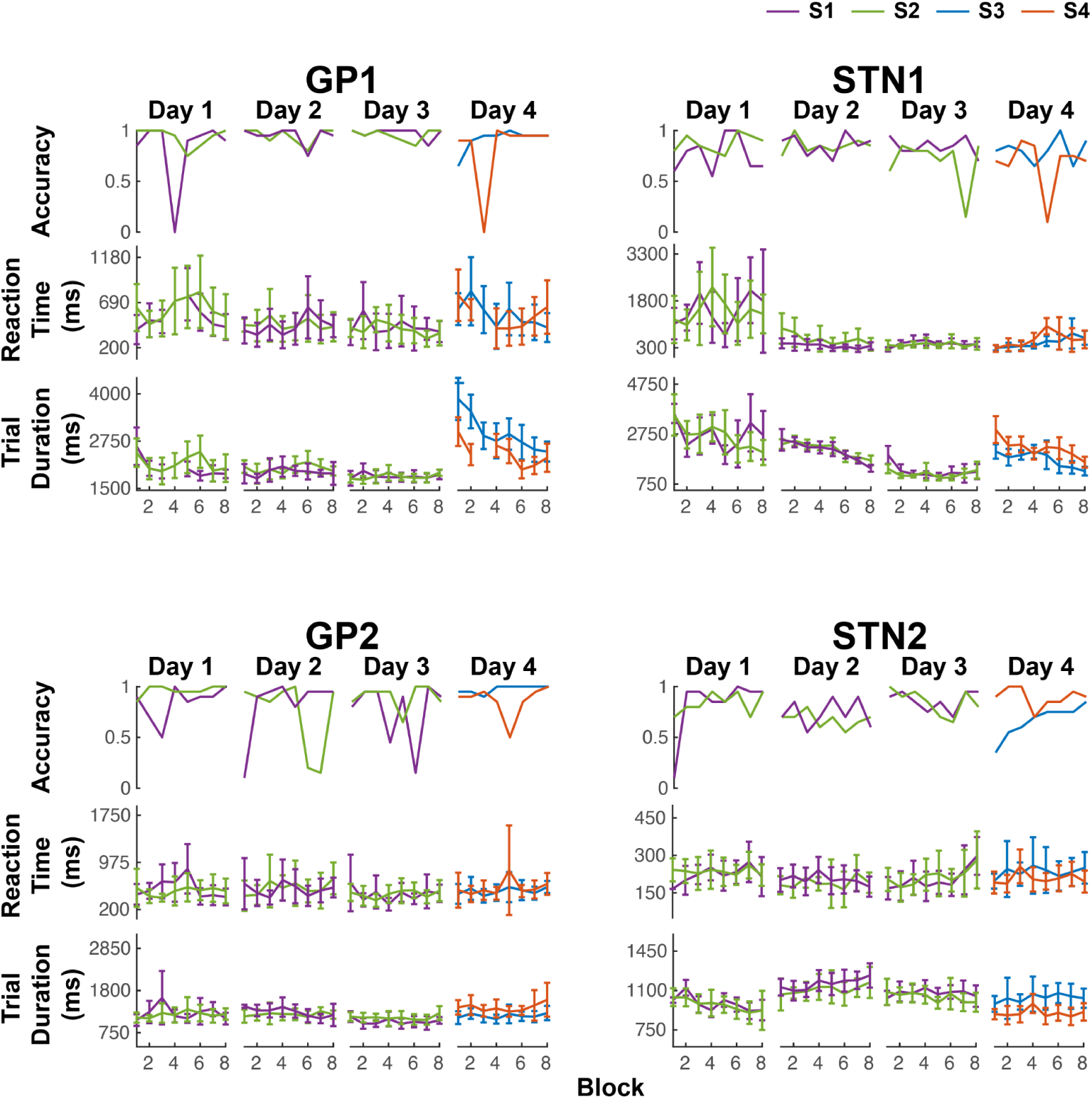
Block average performance data. Per subject: (Top) block average accuracy, (Middle) block average reaction time [cue onset to movement onset], (Bottom) block average trial duration [movement onset to offset]. Error bars indicate ± *s*.

**Supplementary Figure 2.**
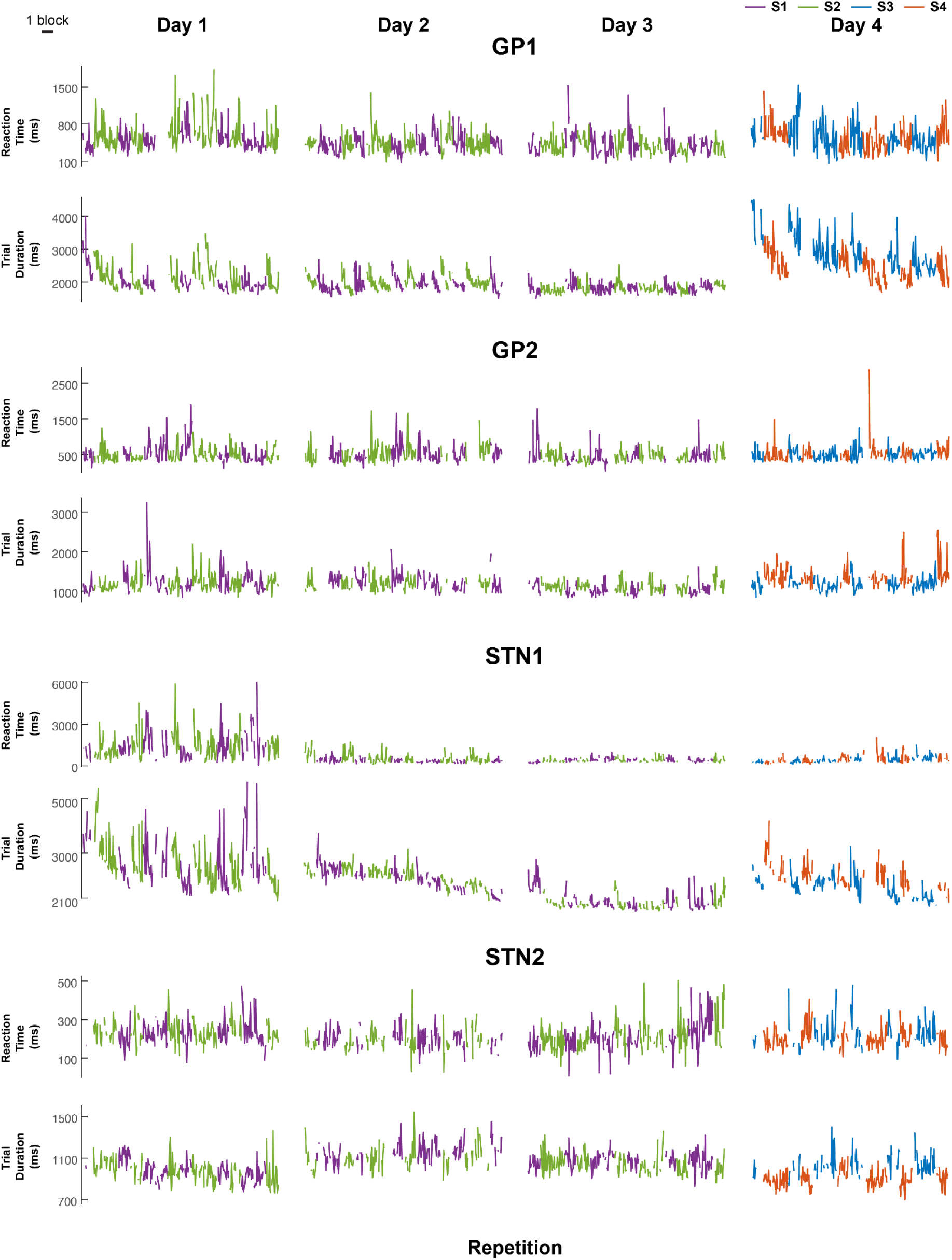
Single-trial performance data. Single-trial reaction time and trial duration for all fully correct trials.

**Supplementary Figure 3.**
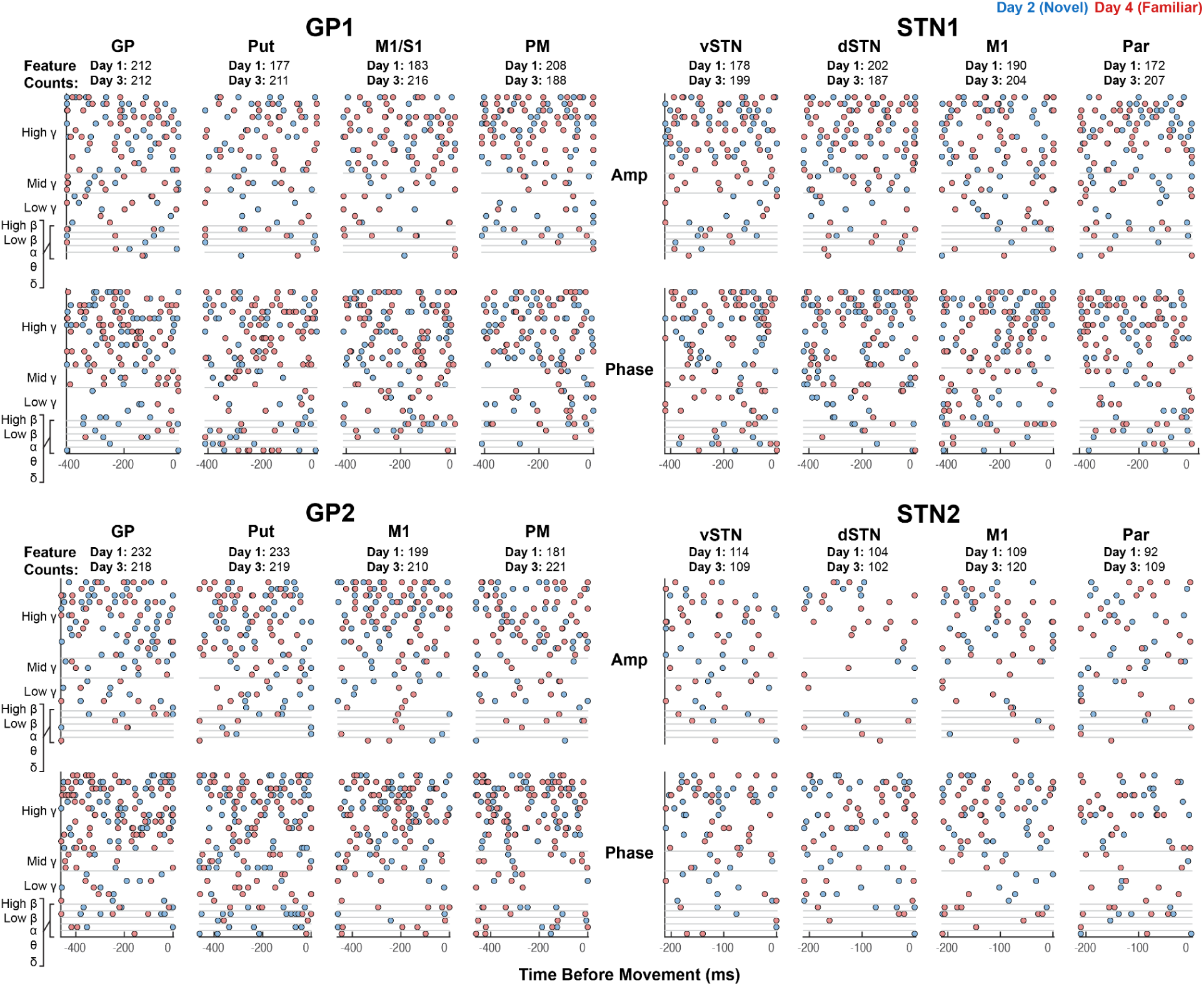
Features selected prior to lasso regularization. Features were selected separately for each model. Total feature count per model (i.e., per subject per channel per day) prior to lasso regularization is shown above each channel’s amplitude and phase feature plots. ***Feature counts are based on each design matrix and thus include a feature for both the sin(phase) and the cos(phase) of each selected phase time-frequency point.*** As a larger number of phase than amplitude time-frequency points tended to reach the 80th percentile cutoff for feature selection, the indicated total feature counts are at least 50% higher than the number of initially selected time-frequency points.

**Supplementary Figure 4.**
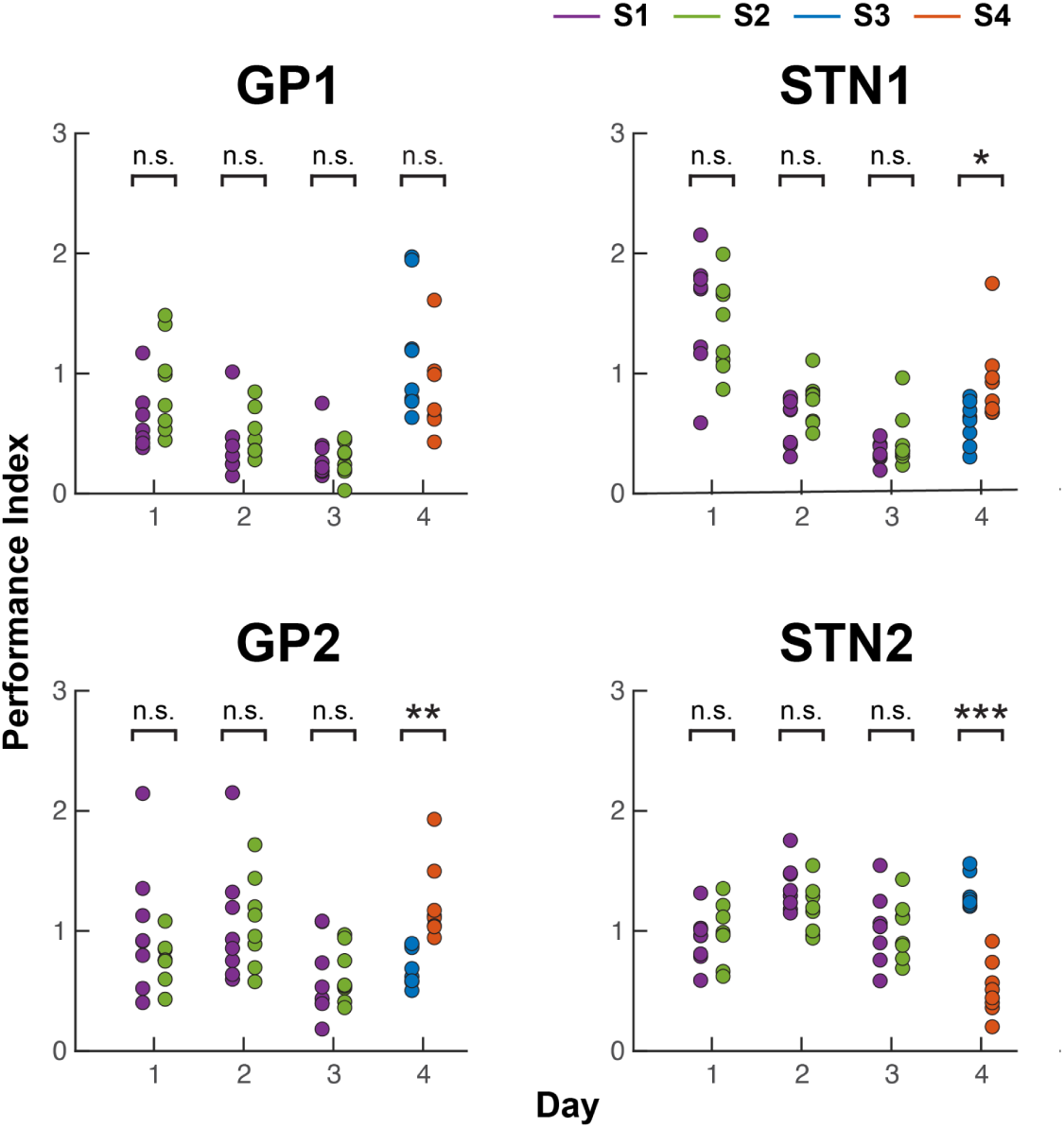
On Days 1–3, overall sequence performance was comparable between S1 and S2. Across-sequence comparison of performance within each day assessed differences in overall performance level (α = 0.05, two-sided, two-sample t-test with unequal variance, *n* = 8 sequence blocks per group except for GP1 Day 1 S1 for which *n* = 7 due to exclusion of 0% accuracy blocks, as composite performance would be poorly defined.). *p < 0.05, **p < 0.01, ***p < 0.001.

**Supplementary Figure 5.**
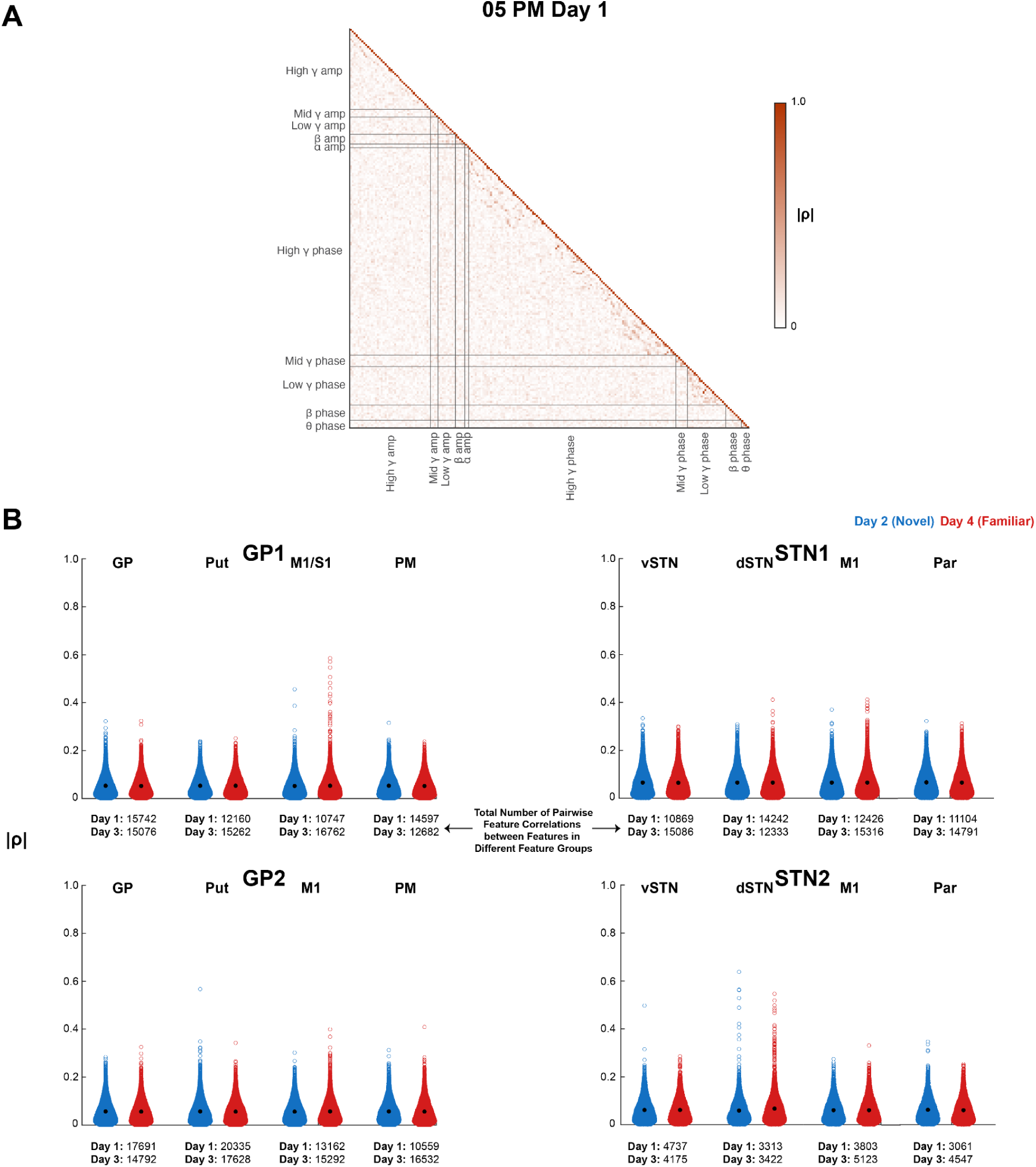
Correlations between features after grouping by canonical frequency band and signal property. (**A**) Example feature correlation matrix for a single model, where light gray lines separate feature groups. (**B**) For each model, a swarm plot of all possible feature correlations between features in different groups. Features were grouped by canonical frequency band and also by signal property, i.e., δ phase features were grouped separately from δ amplitude features, as well as from all other canonical frequency bands. Total number of feature correlations between features in different feature groups for a given model is shown below corresponding swarm plots.

**Supplementary Figure 6.**
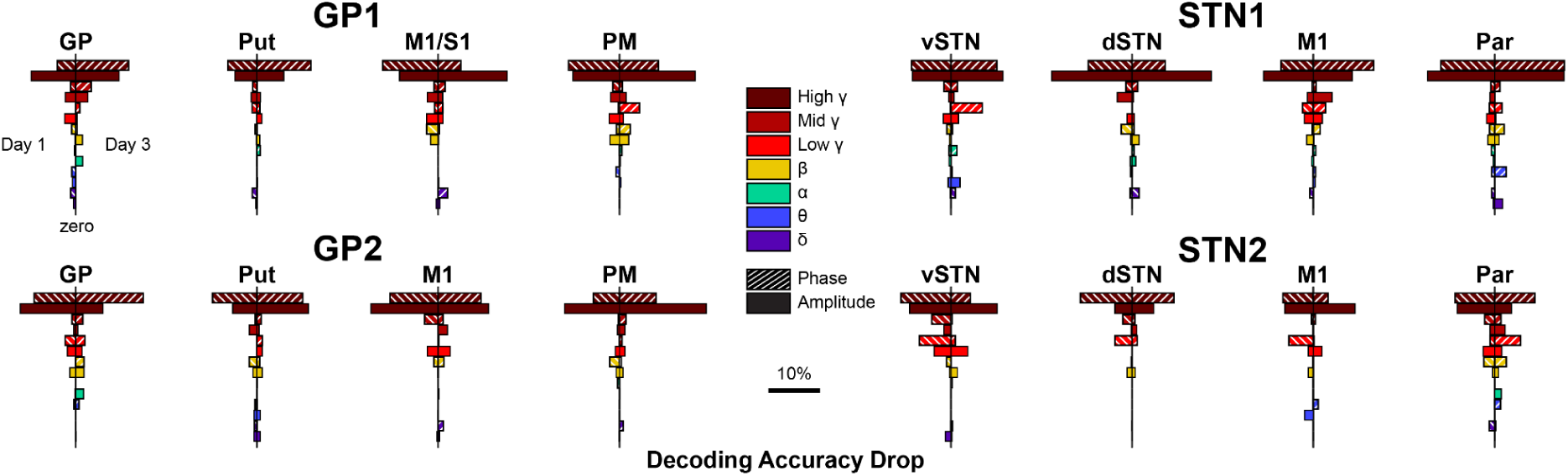
Absolute decoding accuracy decreases for features grouped by canonical frequency band and signal property.

**Supplementary Figure 7.**
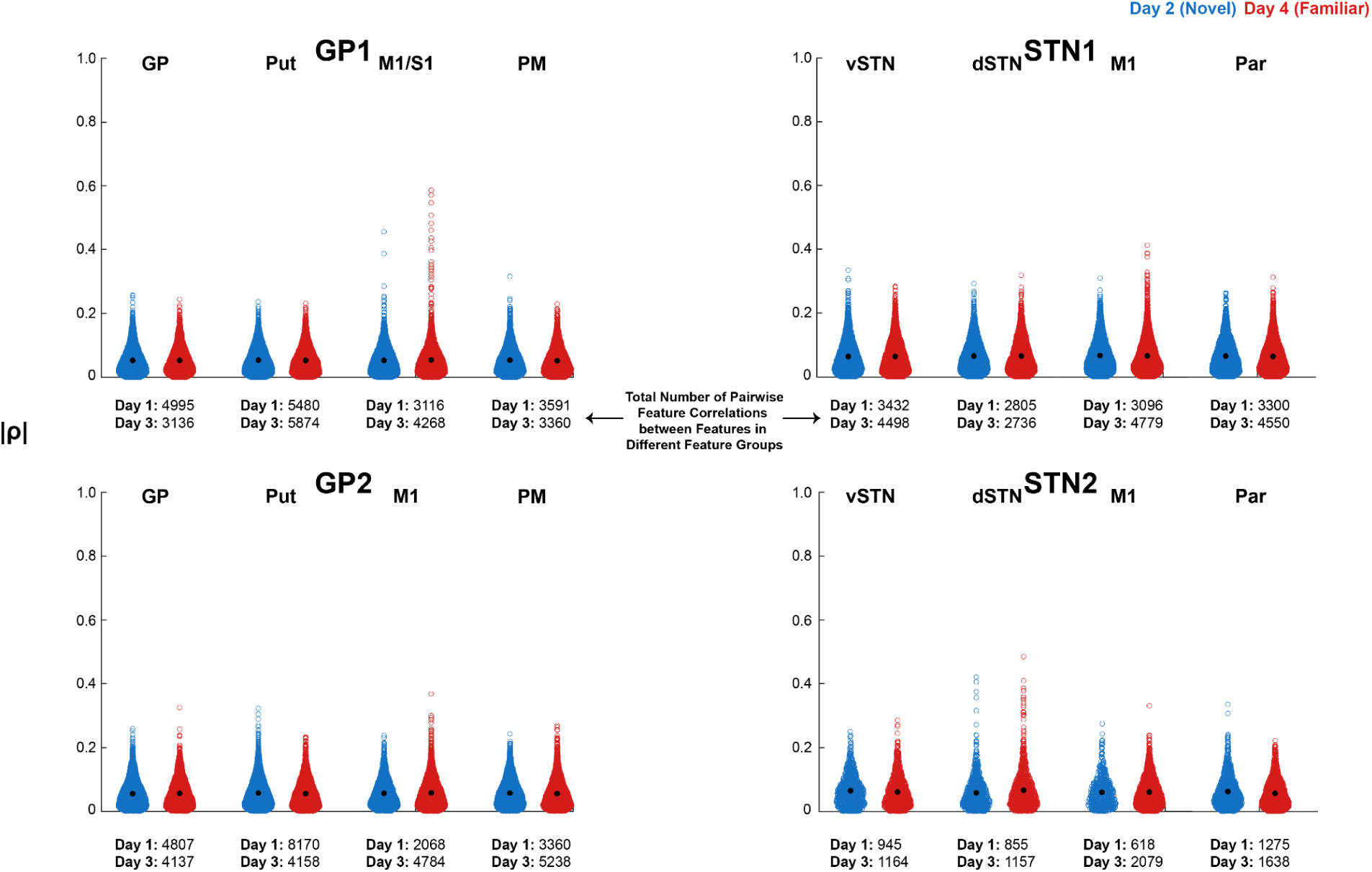
Correlations between features after grouping into 1) δ through β and 2) low γ through high γ. For each model, a swarm plot of all possible feature correlations between features in different groups. Features were grouped into δ through β (0.5–30 Hz) and low γ through high γ (30–250 Hz), with amplitude and phase features grouped together. Total number of feature correlations between features in different groups for a given model is shown below corresponding swarm plots.

**Supplementary Figure 8.**
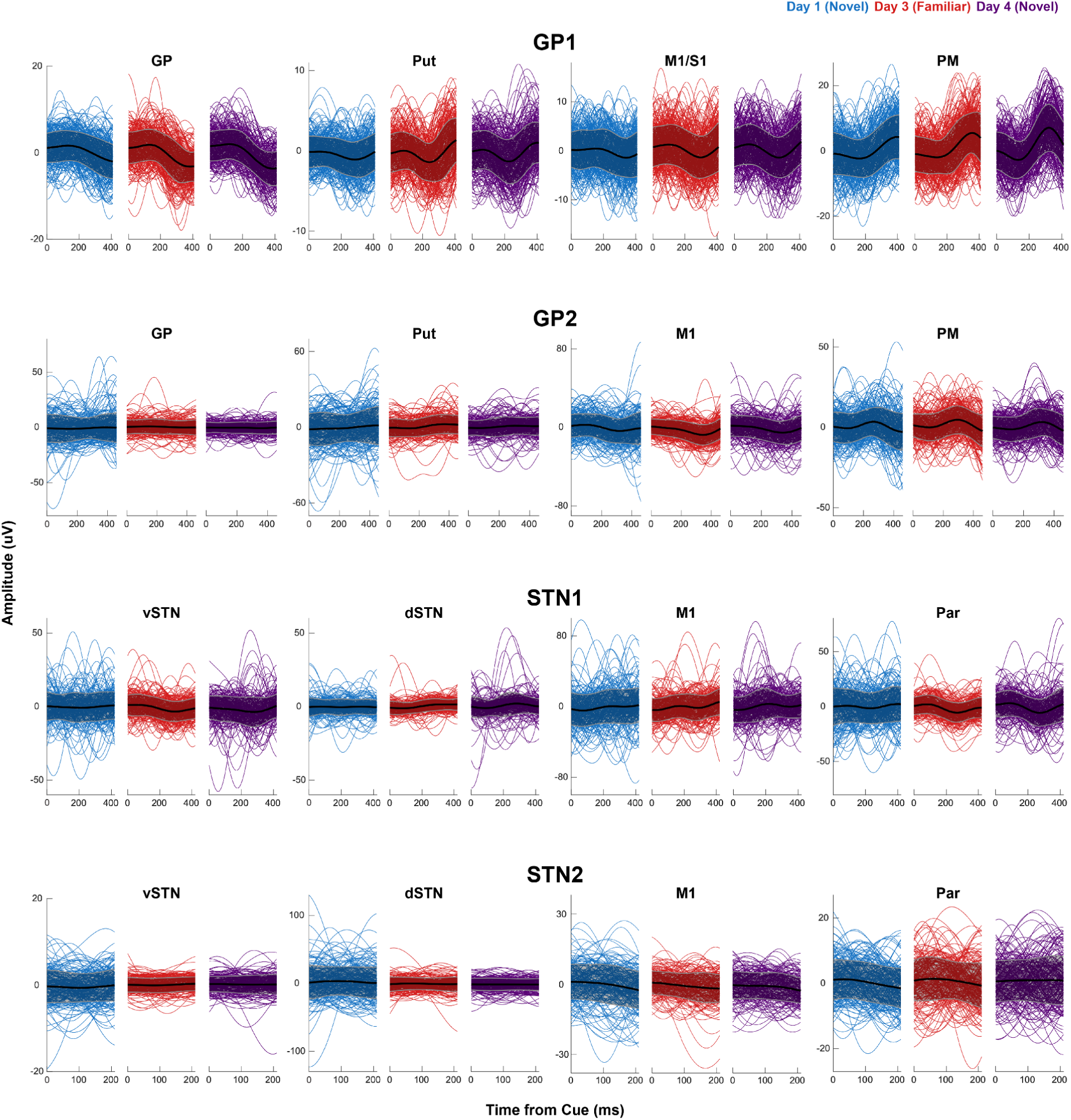
Single-trial δ time domain data. Data is aligned to cue onset and plotted as *x̄* ± *s*.

**Supplementary Figure 9.**
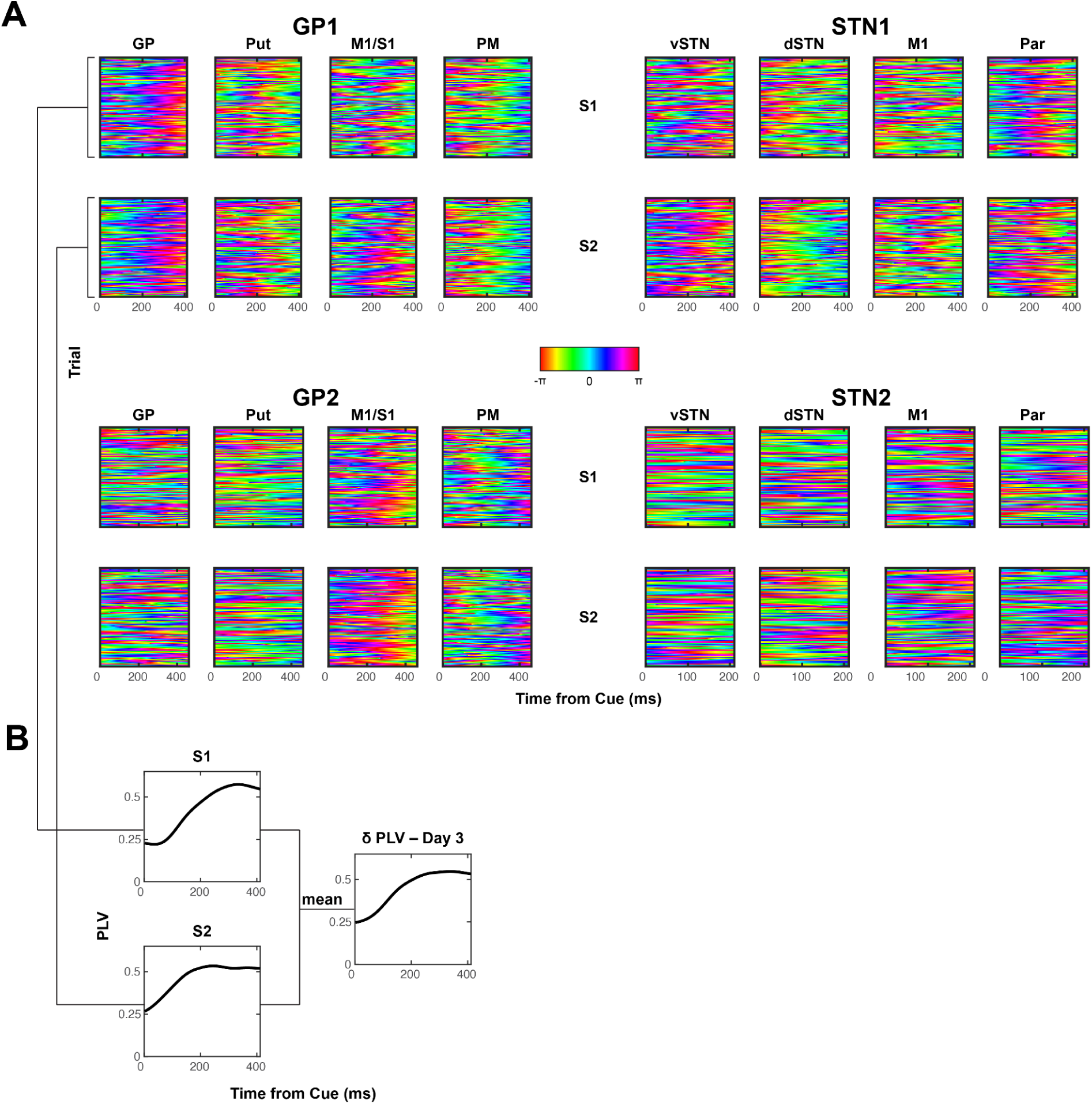
Day 3 single-trial δ phase. (**A**) δ phase data aligned to cue onset. (**B**) Example of phase locking value calculation. Phase locking value was first calculated within sequence and smoothed with a 150 ms-long Gaussian window. Resulting time series were averaged across the two sequences to compute the overall phase locking value for this brain region on this day.

**Supplementary Figure 10.**
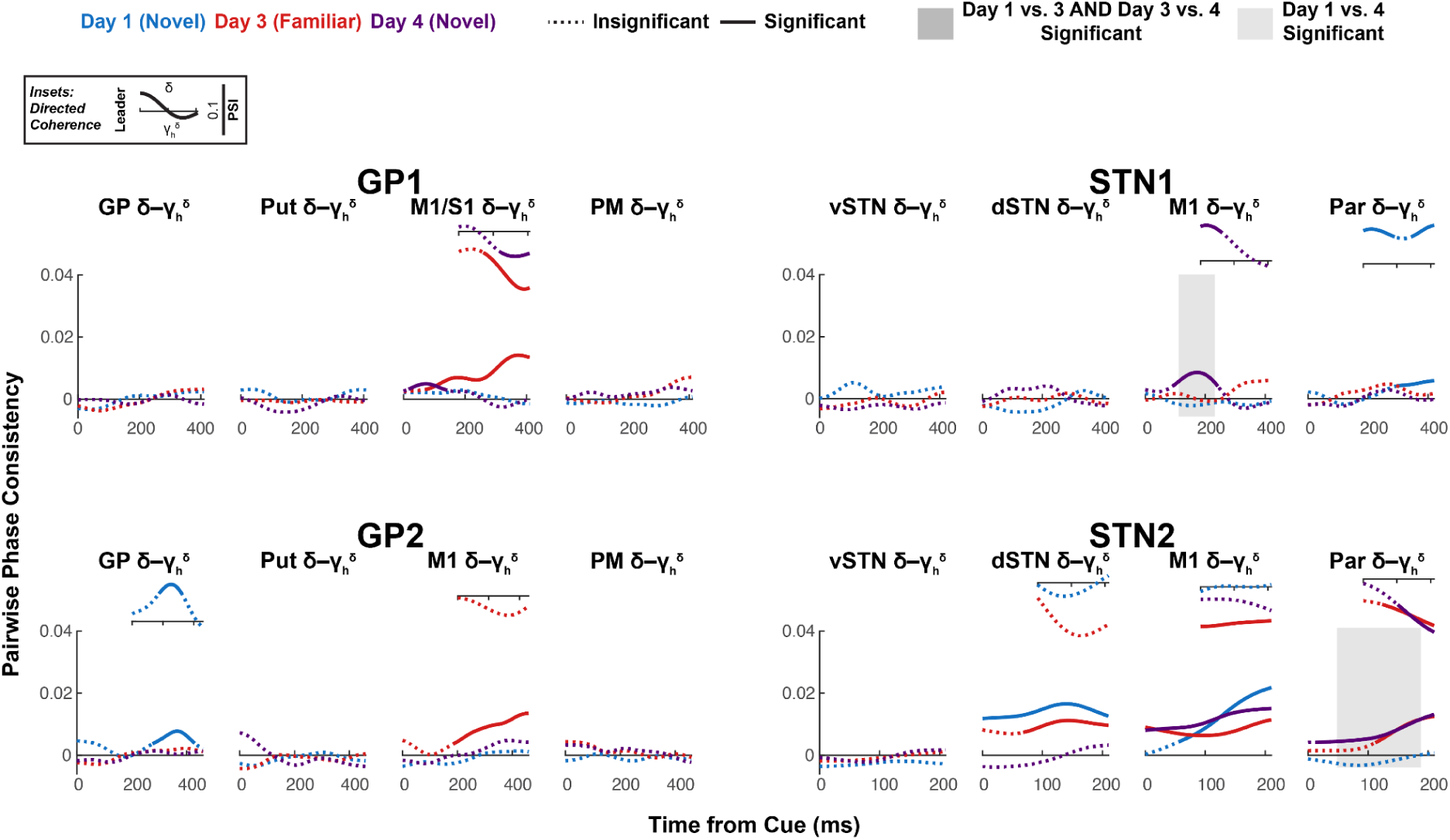
Intraregional δ-γ coupling. (Large plots) Pairwise phase consistency (PPC, undirected measure) was calculated between δ phase and the δ phase of the high γ amplitude envelope. Solid line indicates significant PPC (*h_0_* = coherence is not higher than expected given the phase distribution, *α =* 0.05, one-sided, cluster-based permutation with 10,000 resamples. See **Table 10** for *p*-values.). Shaded box indicates significant difference in PPC between days (*α =* 0.05, two-sided, cluster-based permutation with 10,000 resamples. See **Table 11** for *p*-values.). (Insets) Phase slope index (PSI, directed measure) for significant PPC time series. Solid line indicates significant PSI (*h_0_* = no channel leads, *α =* 0.05, two-sided, cluster-based permutation with 10,000 resamples. See **Table 12** for *p*-values.).

**Supplementary Figure 11.**
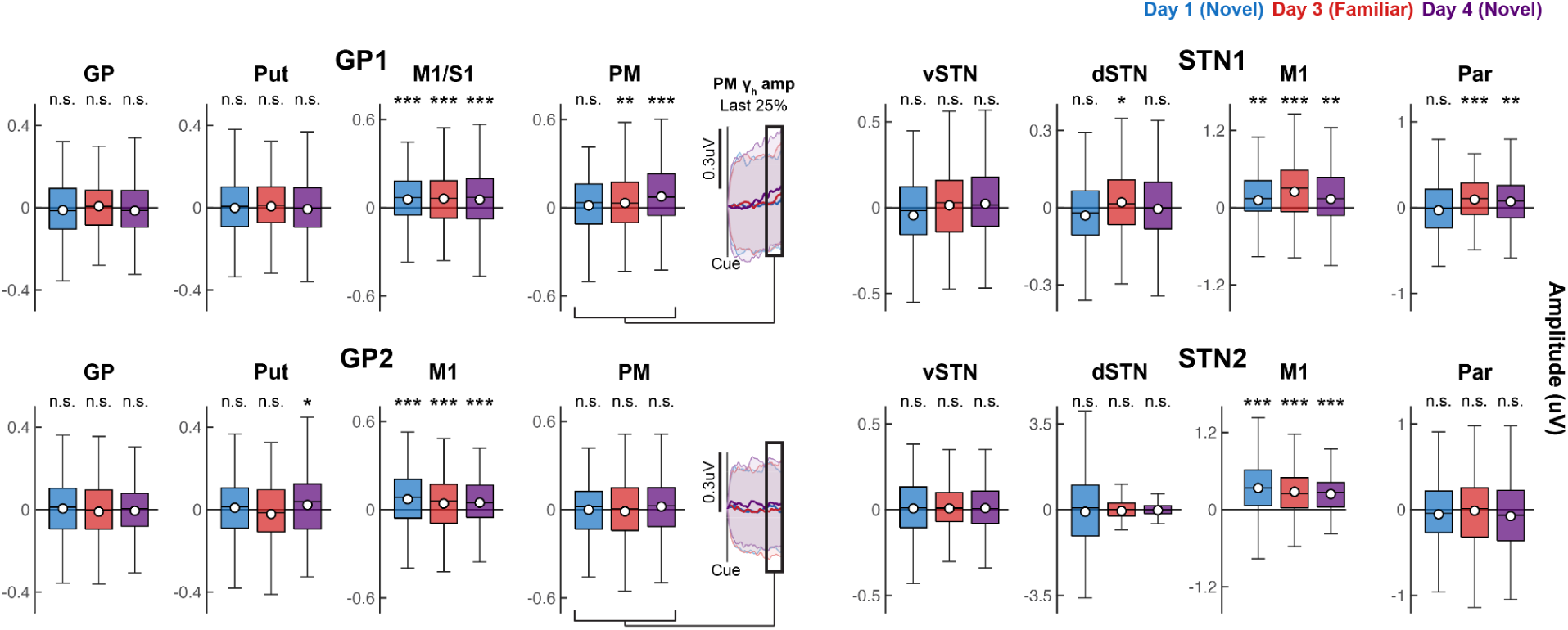
High γ amplitude distributions. (Box plots) Change in high γ amplitude from cue onset to the last 25% of the reaction time period, after linear interpolation of all RT period trials to the same length and smoothing of γ amplitude across time (*α =* 0.05, one-sided, bootstrap estimation of *x̄* with 10,000 resamples. See **Table 13** for *p*-values.). White circle reflects mean; black horizontal line reflects median. Box edges correspond to 25th and 75th percentiles. Whiskers span entire data range excluding outliers. Outliers were computed as 1.5·*IQR* away from the upper or lower quartile and are not shown. (Insets) GP1 and GP2 trial average γ amplitude time series in premotor cortex, with black rectangle indicating the window over which γ amplitude is averaged for individual trials. Error bars indicate ± *s*. \**p* < 0.05, \*\**p* < 0.01, \*\*\**p* < 0.001.

**Supplementary Figure 12.**
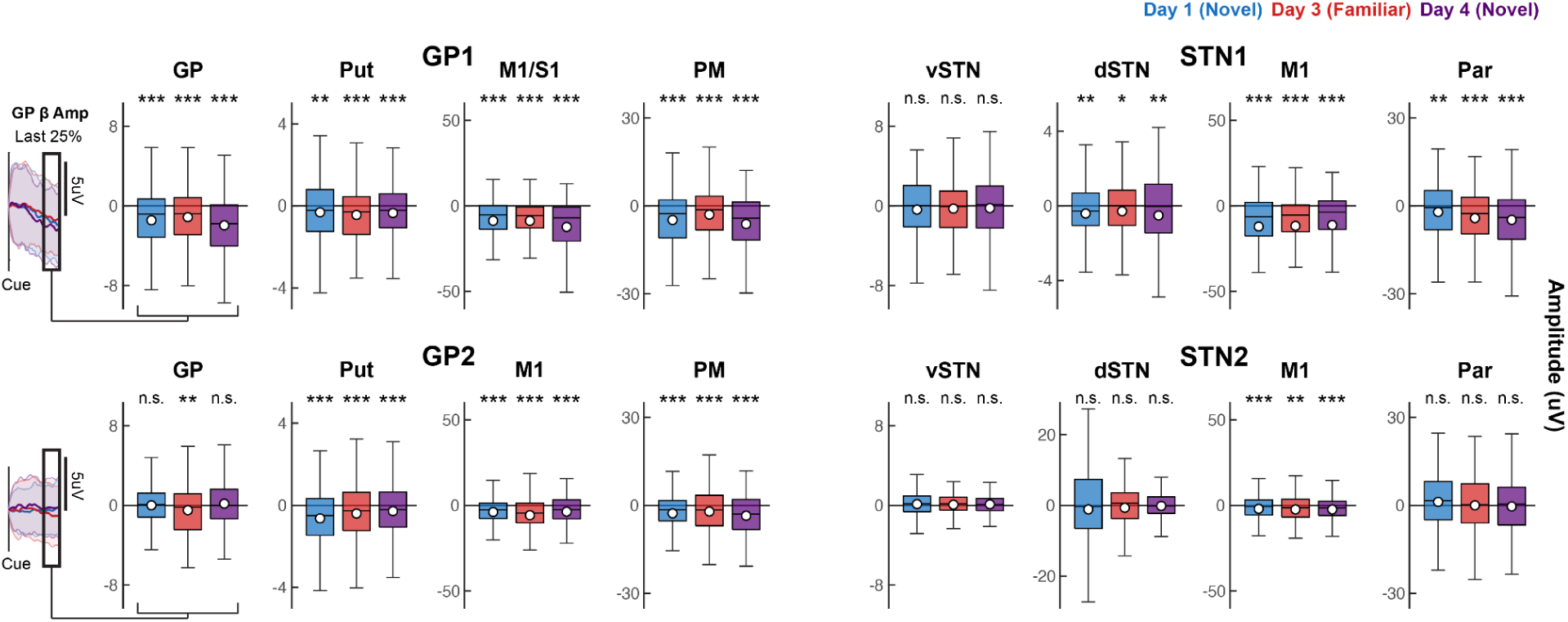
β amplitude distributions. (Box plots) Change in β amplitude from cue onset to the last 25% of the reaction time period, after linear interpolation of all RT period trials to the same length and smoothing of β amplitude across time (*α =* 0.05, one-sided, bootstrap estimation of *x̄* with 10,000 resamples. See **Table 17** for *p*-values.). White circle reflects mean; black horizontal line reflects median. Box edges correspond to 25th and 75th percentiles. Whiskers span entire data range excluding outliers. Outliers were computed as 1.5·*IQR* away from the upper or lower quartile and are not shown. (Insets) GP1 and GP2 trial average β amplitude time series in pallidum, with black rectangle indicating the window over which β amplitude is averaged for individual trials. Error bars indicate ± *s*. \**p* < 0.05, \*\**p* < 0.01, \*\*\**p* < 0.001.

**Supplementary Figure 13.**
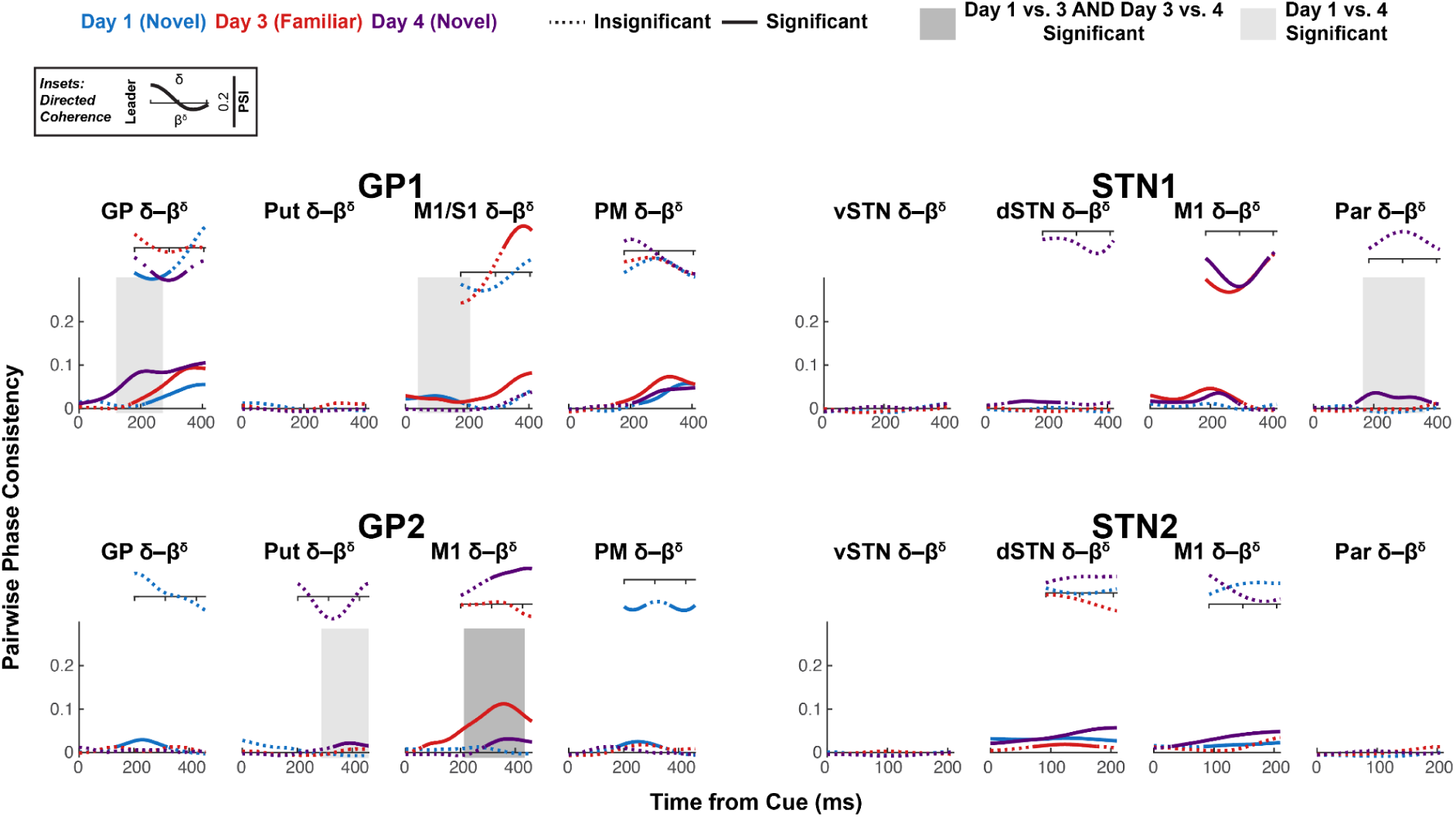
Intraregional δ-β coupling. (Large plots) Pairwise phase consistency (PPC, undirected measure) was calculated between δ phase and the δ phase of the β amplitude envelope. Solid line indicates significant PPC (*h_0_* = coherence is not higher than that expected given the phase distribution, *α =* 0.05, one-sided, cluster-based permutation with 10,000 resamples. See **Table 18** for *p*-values.). Shaded box indicates significant difference in PPC between days (*α =* 0.05, two-sided, cluster-based permutation with 10,000 resamples. See **Table 19** for *p*-values.). (Insets) Phase slope index (PSI, directed measure) for significant PPC time series. Solid line indicates significant PSI (*h_0_* = no channel leads, *α =* 0.05, two-sided, cluster-based permutation with 10,000 resamples. See **Table 20** for *p*-values.).

## REFERENCES

1. Doyon J. Motor sequence learning and movement disorders. Curr Opin Neurol. 2008;21(4):478–483. doi:10.1097/WCO.0b013e328304b6a3

2. Wu T, Hallett M, Chan P. Motor automaticity in Parkinson’s disease. Neurobiol Dis. 2015;82:226–234. doi:10.1016/j.nbd.2015.06.014

3. Marinelli L, Quartarone A, Hallett M, Frazzitta G, Ghilardi MF. The many facets of motor learning and their relevance for Parkinson’s disease. Clin Neurophysiol. 2017;128(7):1127–1141. doi:10.1016/j.clinph.2017.03.042

4. Vasu D, Hui LM, Fong W, Choong P, Chou LW. Evidence-Based Physiotherapeutic Interventions Enhancing Hand Dexterity, Activities of Daily Living and Quality of Life of Parkinson’s Disease Patients: A Systematic Review. Can J Neurol Sci. 2024;1(13). doi:10.1017/cjn.2024.53

5. Rahman S, Griffin HJ, Quinn NP, Jahanshahi M. Quality of life in Parkinson’s disease: The relative importance of the symptoms. Mov Disord. 2008;23(10):1428–1434. doi:10.1002/mds.21667

6. Lee MS, Lyoo CH, Lee MJ, Sim J, Cho H, Choi YH. Impaired finger dexterity in patients with parkinson’s disease correlates with discriminative cutaneous sensory dysfunction. Mov Disord. 2010;25(15):2531–2535. doi:10.1002/mds.23304

7. Foki T, Pirker W, Geißler A, et al. Finger dexterity deficits in Parkinson’s disease and somatosensory cortical dysfunction. Parkinsonism Relat Disord. 2015;21(3):259–265. doi:10.1016/j.parkreldis.2014.12.025

8. Vanbellingen T, Kersten B, Bellion M, et al. Impaired finger dexterity in Parkinson’s disease is associated with praxis function. Brain Cogn. 2011;77(1):48–52. doi:10.1016/j.bandc.2011.06.003

9. Gebhardt A, Vanbellingen T, Baronti F, Kersten B, Bohlhalter S. Poor dopaminergic response of impaired dexterity in Parkinson’s disease: Bradykinesia or limb kinetic apraxia? Mov Disord. 2008;23(12):1701–1706. doi:10.1002/mds.22199

10. Ramirez-Zamora A, Ostrem JL. Globus Pallidus Interna or Subthalamic Nucleus Deep Brain Stimulation for Parkinson Disease: A Review. JAMA Neurol. 2018;75(3):367–372. doi:10.1001/jamaneurol.2017.4321

11. Eisinger RS, Cernera S, Gittis A, Gunduz A, Okun MS. A review of basal ganglia circuits and physiology: Application to deep brain stimulation. Parkinsonism Relat Disord. 2019;59:9–20. doi:10.1016/j.parkreldis.2019.01.009

12. Muehlberg C, Fricke C, Wegscheider M, et al. Motor learning is independent of effects of subthalamic deep brain stimulation on motor execution. Brain Commun. 2023;5(2):fcad070. doi:10.1093/braincomms/fcad070

13. Ingram LA, Carroll VK, Butler AA, Brodie MA, Gandevia SC, Lord SR. Quantifying upper limb motor impairment in people with Parkinson’s disease: a physiological profiling approach. PeerJ. 2021;9:e10735. doi:10.7717/peerj.10735

14. Teixeira NB, Alouche SR. The dual task performance in Parkinson’s disease. Braz J Phys Ther. 2007;11:127–132. doi:10.1590/S1413-35552007000200007

15. Park JE. Apraxia: Review and Update. J Clin Neurol. 2017;13(4):317–324. doi:10.3988/jcn.2017.13.4.317

16. Heilman KM. Chapter 24 - Action programming disorders associated with Parkinson’s disease. In: Martin CR, Preedy VR, eds. Genetics, Neurology, Behavior, and Diet in Parkinson’s Disease. Academic Press; 2020:377–393. doi:10.1016/B978-0-12-815950-7.00024-2

17. Kumari LS, Kouzani AZ. Phase-Dependent Deep Brain Stimulation: A Review. Brain Sci. 2021;11(4):414. doi:10.3390/brainsci11040414

18. Simmonds DJ, Pekar JJ, Mostofsky SH. Meta-analysis of Go/No-go tasks demonstrating that fMRI activation associated with response inhibition is task-dependent. Neuropsychologia. 2008;46(1):224–232. doi:10.1016/j.neuropsychologia.2007.07.015

19. Ariani G, Diedrichsen J. Sequence learning is driven by improvements in motor planning. J Neurophysiol. 2019;121(6):2088–2100. doi:10.1152/jn.00041.2019

20. Ariani G, Kordjazi N, Pruszynski JA, Diedrichsen J. The Planning Horizon for Movement Sequences. eneuro. 2021;8(2):ENEURO.0085-21.2021. doi:10.1523/ENEURO.0085-21.2021

21. Verwey WB, Shea CH, Wright DL. A cognitive framework for explaining serial processing and sequence execution strategies. Psychon Bull Rev. 2015;22(1):54–77. doi:10.3758/s13423-014-0773-4

22. Wong AL, Haith AM, Krakauer JW. Motor Planning. The Neuroscientist. 2015;21(4):385–398. doi:10.1177/1073858414541484

23. Verwey WB. Evidence for the development of concurrent processing in a sequential keypressing task. Acta Psychol (Amst). 1994;85(3):245–262. doi:10.1016/0001-6918(94)90038-8

24. Khanna P, Totten D, Novik L, Jeffrey Roberts, Morecraft RJ, Ganguly K. Low-frequency stimulation enhances ensemble co-firing and dexterity after stroke. Cell. 2021;184(4):912–930.e20. doi:10.1016/j.cell.2021.01.023

25. Ganguly K, Carmena JM. Emergence of a Stable Cortical Map for Neuroprosthetic Control. PLOS Biol. 2009;7(7):e1000153. doi:10.1371/journal.pbio.1000153

26. Peters AJ, Chen SX, Komiyama T. Emergence of reproducible spatiotemporal activity during motor learning. Nature. 2014;510(7504):263–267. doi:10.1038/nature13235

27. Rostami V, Rost T, Schmitt FJ, Albada SJ van, Riehle A, Nawrot MP. Spiking attractor model of motor cortex explains modulation of neural and behavioral variability by prior target information. Published online March 5, 2024:2020.02.27.968339. doi:10.1101/2020.02.27.968339

28. Guo L, Kondapavulur S, Lemke SM, Won SJ, Ganguly K. Coordinated increase of reliable cortical and striatal ensemble activations during recovery after stroke. Cell Rep. 2021;36(2):109370. doi:10.1016/j.celrep.2021.109370

29. Ganguly K, Khanna P, Morecraft RJ, Lin DJ. Modulation of neural co-firing to enhance network transmission and improve motor function after stroke. Neuron. 2022;110(15):2363–2385. doi:10.1016/j.neuron.2022.06.024

30. Carrillo-Reid L. Neuronal ensembles in memory processes. Semin Cell Dev Biol. 2022;125:136–143. doi:10.1016/j.semcdb.2021.04.004

31. Lu X, Ashe J. Anticipatory activity in primary motor cortex codes memorized movement sequences. Neuron. 2005;45(6):967–973. doi:10.1016/j.neuron.2005.01.036

32. Hatsopoulos NG, Paninski L, Donoghue JP. Sequential movement representations based on correlated neuronal activity. Exp Brain Res. 2003;149:478–486. doi:10.1007/s00221-003-1385-9

33. Hahn G, Ponce-Alvarez A, Deco G, Aertsen A, Kumar A. Portraits of communication in neuronal networks. Nat Rev Neurosci. 2019;20(2):117–127. doi:10.1038/s41583-018-0094-0

34. Mazzoni A, Whittingstall K, Brunel N, Logothetis NK, Panzeri S. Understanding the relationships between spike rate and delta/gamma frequency bands of LFPs and EEGs using a local cortical network model. NeuroImage. 2010;52(3):956–972. doi:10.1016/j.neuroimage.2009.12.040

35. Schroeder CE, Lakatos P. Low-frequency neuronal oscillations as instruments of sensory selection. Trends Neurosci. 2009;32(1):10.1016/j.tins.2008.09.012. doi:10.1016/j.tins.2008.09.012

36. Lakatos P, Karmos G, Mehta AD, Ulbert I, Schroeder CE. Entrainment of neuronal oscillations as a mechanism of attentional selection. Science. 2008;320(5872):110–113. doi:10.1126/science.1154735

37. Muller L, Chavane F, Reynolds J, Sejnowski TJ. Cortical travelling waves: mechanisms and computational principles. Nat Rev Neurosci. 2018;19(5):255–268. doi:10.1038/nrn.2018.20

38. Whittingstall K, Logothetis NK. Frequency-band coupling in surface EEG reflects spiking activity in monkey visual cortex. Neuron. 2009;64(2):281–289. doi:10.1016/j.neuron.2009.08.016

39. Hamel-Thibault A, Thénault F, Whittingstall K, Bernier PM. Delta-Band Oscillations in Motor Regions Predict Hand Selection for Reaching. Cereb Cortex. 2018;28(2):574–584. doi:10.1093/cercor/bhw392

40. Ferreri F, Vecchio F, Ponzo D, Pasqualetti P, Rossini PM. Time-varying coupling of EEG oscillations predicts excitability fluctuations in the primary motor cortex as reflected by motor evoked potentials amplitude: An EEG-TMS study. Hum Brain Mapp. 2014;35(5):1969–1980. doi:10.1002/hbm.22306

41. Fasano A, Mazzoni A, Falotico E. Reaching and Grasping Movements in Parkinson’s Disease: A Review. J Park Dis. 2022;12(4):1083–1113. doi:10.3233/JPD-213082

42. Brown P. Oscillatory nature of human basal ganglia activity: Relationship to the pathophysiology of Parkinson’s disease. Mov Disord. 2003;18(4):357–363. doi:10.1002/mds.10358

43. Meissner SN, Krause V, Südmeyer M, Hartmann CJ, Pollok B. Pre-stimulus beta power modulation during motor sequence learning is reduced in ‘Parkinson’s disease. NeuroImage Clin. 2019;24:102057. doi:10.1016/j.nicl.2019.102057

44. Singh A. Oscillatory activity in the cortico-basal ganglia-thalamic neural circuits in Parkinson’s disease. Eur J Neurosci. 2018;48(8):2869–2878. doi:10.1111/ejn.13853

45. Rockhill AP, Mantovani A, Stedelin B, Nerison CS, Raslan AM, Swann NC. Stereo-EEG recordings extend known distributions of canonical movement-related oscillations. J Neural Eng. 2023;20(1):016007. doi:10.1088/1741-2552/acae0a

46. Mäki H, Ilmoniemi RJ. EEG oscillations and magnetically evoked motor potentials reflect motor system excitability in overlapping neuronal populations. Clin Neurophysiol. 2010;121(4):492–501. doi:10.1016/j.clinph.2009.11.078

47. Schutter DJLG, Hortensius R. Brain oscillations and frequency-dependent modulation of cortical excitability. Brain Stimulat. 2011;4(2):97–103. doi:10.1016/j.brs.2010.07.002

48. Berger B, Minarik T, Liuzzi G, Hummel FC, Sauseng P. EEG Oscillatory Phase-Dependent Markers of Corticospinal Excitability in the Resting Brain. BioMed Res Int. 2014;2014(1):936096. doi:10.1155/2014/936096

49. Bhatt MB, Bowen S, Rossiter HE, et al. Computational modelling of movement-related beta-oscillatory dynamics in human motor cortex. NeuroImage. 2016;133:224–232. doi:10.1016/j.neuroimage.2016.02.078

50. Kaufman MT, Churchland MM, Ryu SI, Shenoy KV. Cortical activity in the null space: permitting preparation without movement. Nat Neurosci. 2014;17(3):440–448. doi:10.1038/nn.3643

51. Churchland MM, Shenoy KV. Preparatory activity and the expansive null-space. Nat Rev Neurosci. 2024;25(4):213–236. doi:10.1038/s41583-024-00796-z

52. Lemke SM, Ramanathan DS, Guo L, Won SJ, Ganguly K. Emergent modular neural control drives coordinated motor actions. Nat Neurosci. 2019;22(7):1122–1131. doi:10.1038/s41593-019-0407-2

53. Chang EF. Towards Large-Scale, Human-Based, Mesoscopic Neurotechnologies. Neuron. 2015;86(1):68–78. doi:10.1016/j.neuron.2015.03.037

54. Abrahamse E, Ruitenberg M, De Kleine E, Verwey WB. Control of automated behavior: insights from the discrete sequence production task. Front Hum Neurosci. 2013;7. doi:10.3389/fnhum.2013.00082

55. VanRullen R. How to Evaluate Phase Differences between Trial Groups in Ongoing Electrophysiological Signals. Front Neurosci. 2016;10:426. doi:10.3389/fnins.2016.00426

56. Lachaux JP, Rodriguez E, Martinerie J, Varela FJ. Measuring phase synchrony in brain signals. Hum Brain Mapp. 1999;8(4):194–208. doi:10.1002/(SICI)1097-0193(1999)8:4<194::AID-HBM4>3.0.CO;2-C

57. Vinck M, van Wingerden M, Womelsdorf T, Fries P, Pennartz CMA. The pairwise phase consistency: A bias-free measure of rhythmic neuronal synchronization. NeuroImage. 2010;51(1):112–122. doi:10.1016/j.neuroimage.2010.01.073

58. Nolte G, Ziehe A, Nikulin VV, et al. Robustly Estimating the Flow Direction of Information in Complex Physical Systems. Phys Rev Lett. 2008;100(23):234101. doi:10.1103/PhysRevLett.100.234101

59. Taylor JA, Ivry RB. Flexible Cognitive Strategies during Motor Learning. PLOS Comput Biol. 2011;7(3):e1001096. doi:10.1371/journal.pcbi.1001096

60. Wulf G. Attentional focus and motor learning: a review of 15 years. Int Rev Sport Exerc Psychol. 2013;6(1):77–104. doi:10.1080/1750984X.2012.723728

61. Song JH. The role of attention in motor control and learning. Curr Opin Psychol. 2019;29:261–265. doi:10.1016/j.copsyc.2019.08.002

62. Clark D, Ivry RB. Multiple systems for motor skill learning. Wiley Interdiscip Rev Cogn Sci. 2010;1(4):461–467. 10.1002/wcs.56

63. Serrien DJ, Ivry RB, Swinnen SP. The missing link between action and cognition. Prog Neurobiol. 2007;82(2):95–107. doi:10.1016/j.pneurobio.2007.02.003

64. Doyon J, Benali H. Reorganization and plasticity in the adult brain during learning of motor skills. Curr Opin Neurobiol. 2005;15(2):161–167. doi:10.1016/j.conb.2005.03.004

65. Goetz CG, Tilley BC, Shaftman SR, et al. Movement Disorder Society-sponsored revision of the Unified Parkinson’s Disease Rating Scale (MDS-UPDRS): Scale presentation and clinimetric testing results. Mov Disord. 2008;23(15):2129–2170. doi:10.1002/mds.22340

66. Saleh M, Reimer J, Penn R, Ojakangas CL, Hatsopoulos NG. Fast and Slow Oscillations in Human Primary Motor Cortex Predict Oncoming Behaviorally Relevant Cues. Neuron. 2010;65(4):461–471. doi:10.1016/j.neuron.2010.02.001

67. Crone NE, Miglioretti DL, Gordon B, Lesser RP. Functional mapping of human sensorimotor cortex with electrocorticographic spectral analysis. II. Event-related synchronization in the gamma band. Brain. 1998;121(12):2301–2315. doi:10.1093/brain/121.12.2301

68. Herrojo Ruiz M, Brücke C, Nikulin VV, Schneider GH, Kühn AA. Beta-band amplitude oscillations in the human internal globus pallidus support the encoding of sequence boundaries during initial sensorimotor sequence learning. NeuroImage. 2014;85:779–793. doi:10.1016/j.neuroimage.2013.05.085

69. Herrojo Ruiz M, Rusconi M, Brücke C, Haynes JD, Schönecker T, Kühn AA. Encoding of sequence boundaries in the subthalamic nucleus of patients with Parkinson’s disease. Brain J Neurol. 2014;137(Pt 10):2715–2730. doi:10.1093/brain/awu191

70. Jenkinson N, Kühn AA, Brown P. Gamma oscillations in the human basal ganglia. Exp Neurol. 2013;245:72–76. doi:10.1016/j.expneurol.2012.07.005

71. Muralidharan V, Aron AR. Behavioral Induction of a High Beta State in Sensorimotor Cortex Leads to Movement Slowing. J Cogn Neurosci. 2021;33(7):1311–1328. doi:10.1162/jocn_a_01717

72. Torrecillos F, Tinkhauser G, Fischer P, et al. Modulation of Beta Bursts in the Subthalamic Nucleus Predicts Motor Performance. J Neurosci. 2018;38(41):8905–8917. doi:10.1523/JNEUROSCI.1314-18.2018

73. Yin Z, Zhu G, Zhao B, et al. Local field potentials in Parkinson’s disease: A frequency-based review. Neurobiol Dis. 2021;155:105372. doi:10.1016/j.nbd.2021.105372

74. Pascual-Leone A, Valls-Solé J, Brasil-Neto JP, Cohen LG, Hallett M. Akinesia in Parkinson’s disease. I. Shortening of simple reaction time with focal, single-pulse transcranial magnetic stimulation. Neurology. 1994;44(5):884–884. doi:10.1212/WNL.44.5.884

75. Combrisson E, Perrone-Bertolotti M, Soto JL, et al. From intentions to actions: Neural oscillations encode motor processes through phase, amplitude and phase-amplitude coupling. NeuroImage. 2017;147:473–487. doi:10.1016/j.neuroimage.2016.11.042

76. Attaheri A, Choisdealbha ÁN, Di Liberto GM, et al. Delta- and theta-band cortical tracking and phase-amplitude coupling to sung speech by infants. NeuroImage. 2022;247:118698. doi:10.1016/j.neuroimage.2021.118698

77. Natraj N, Silversmith DB, Chang EF, Ganguly K. Compartmentalized dynamics within a common multi-area mesoscale manifold represent a repertoire of human hand movements. Neuron. 2022;110(1):154–174.e12. doi:10.1016/j.neuron.2021.10.002

78. Ramanathan DS, Guo L, Gulati T, et al. Low-frequency cortical activity is a neuromodulatory target that tracks recovery after stroke. Nat Med. 2018;24(8):1257–1267. doi:10.1038/s41591-018-0058-y

79. Pal A, Pegwal N, Behari M, Sharma R. High delta and gamma EEG power in resting state characterise dementia in Parkinson’s patients. Biomark Neuropsychiatry. 2020;3:100027. doi:10.1016/j.bionps.2020.100027

80. Ponsen MM, Stam CJ, Bosboom JLW, Berendse HW, Hillebrand A. A three dimensional anatomical view of oscillatory resting-state activity and functional connectivity in Parkinson’s disease related dementia: An MEG study using atlas-based beamforming. NeuroImage Clin. 2013;2:95–102. doi:10.1016/j.nicl.2012.11.007

81. Rucco R, Lardone A, Liparoti M, et al. Brain Networks and Cognitive Impairment in Parkinson’s Disease. Brain Connect. 2022;12(5):465–475. doi:10.1089/brain.2020.0985

82. Ray S, Crone NE, Niebur E, Franaszczuk PJ, Hsiao SS. Neural Correlates of High-Gamma Oscillations (60–200 Hz) in Macaque Local Field Potentials and Their Potential Implications in Electrocorticography. J Neurosci. 2008;28(45):11526–11536. doi:10.1523/JNEUROSCI.2848-08.2008

83. Rusu SI, Pennartz CMA. Learning, memory and consolidation mechanisms for behavioral control in hierarchically organized cortico-basal ganglia systems. Hippocampus. 2020;30(1):73–98. doi:10.1002/hipo.23167

84. Caligiore D, Arbib MA, Miall RC, Baldassarre G. The super-learning hypothesis: Integrating learning processes across cortex, cerebellum and basal ganglia. Neurosci Biobehav Rev. 2019;100:19–34. doi:10.1016/j.neubiorev.2019.02.008

85. Simonyan K. Recent advances in understanding the role of the basal ganglia. F1000Research. 2019;8:F1000 Faculty Rev-122. doi:10.12688/f1000research.16524.1

86. Klaus A, Silva JA da, Costa RM. What, If, and When to Move: Basal Ganglia Circuits and Self-Paced Action Initiation. Annu Rev Neurosci. 2019;42(Volume 42, 2019):459–483. doi:10.1146/annurev-neuro-072116-031033

87. Balleine BW, Dezfouli A, Ito M, Doya K. Hierarchical control of goal-directed action in the cortical–basal ganglia network. Curr Opin Behav Sci. 2015;5:1–7. doi:10.1016/j.cobeha.2015.06.001

88. Milardi D, Quartarone A, Bramanti A, et al. The Cortico-Basal Ganglia-Cerebellar Network: Past, Present and Future Perspectives. Front Syst Neurosci. 2019;13. doi:10.3389/fnsys.2019.00061

89. Toni I, Rowe J, Stephan KE, Passingham RE. Changes of Cortico-striatal Effective Connectivity during Visuomotor Learning. Cereb Cortex. 2002;12(10):1040–1047. doi:10.1093/cercor/12.10.1040

90. Doyon J, Penhune V, Ungerleider LG. Distinct contribution of the cortico-striatal and cortico-cerebellar systems to motor skill learning. Neuropsychologia. 2003;41(3):252–262. doi:10.1016/S0028-3932(02)00158-6

91. Debas K, Carrier J, Barakat M, et al. Off-line consolidation of motor sequence learning results in greater integration within a cortico-striatal functional network. NeuroImage. 2014;99:50–58. doi:10.1016/j.neuroimage.2014.05.022

92. Canavier CC. Phase-resetting as a tool of information transmission. Curr Opin Neurobiol. 2015;31:206–213. doi:10.1016/j.conb.2014.12.003

93. Harmony T. The functional significance of delta oscillations in cognitive processing. Front Integr Neurosci. 2013;7. doi:10.3389/fnint.2013.00083

94. Voloh B, Womelsdorf T. A Role of Phase-Resetting in Coordinating Large Scale Neural Networks During Attention and Goal-Directed Behavior. Front Syst Neurosci. 2016;10. doi:10.3389/fnsys.2016.00018

95. Levy R, Ashby P, Hutchison WD, Lang AE, Lozano AM, Dostrovsky JO. Dependence of subthalamic nucleus oscillations on movement and dopamine in Parkinson’s disease. Brain. 2002;125(6):1196–1209. doi:10.1093/brain/awf128

96. Zhuang P, Hallett M, Meng D, Zhang Y, Li Y. Characteristics of oscillatory activity in the globus pallidus internus in patients with Parkinson’s disease (P1.8-028). Neurology. 2019;92(15 Supplement). doi:10.1212/WNL.92.15_supplement.P1.8-028

97. Whalen TC, Willard AM, Rubin JE, Gittis AH. Delta oscillations are a robust biomarker of dopamine depletion severity and motor dysfunction in awake mice. J Neurophysiol. 2020;124(2):312–329. doi:10.1152/jn.00158.2020

98. Tosin MHS, Stebbins GT, Comella C, Patterson CG, Hall DA, SPARX Study Group. Does MDS-UPDRS Provide Greater Sensitivity to Mild Disease than UPDRS in De Novo Parkinson’s Disease? Mov Disord Clin Pract. 2021;8(7):1092–1099. doi:10.1002/mdc3.13329

99. Dann B, Michaels JA, Schaffelhofer S, Scherberger H. Uniting functional network topology and oscillations in the fronto-parietal single unit network of behaving primates. Stephan KE, ed. eLife. 2016;5:e15719. doi:10.7554/eLife.15719

100. Wyart V, de Gardelle V, Scholl J, Summerfield C. Rhythmic fluctuations in evidence accumulation during decision making in the human brain. Neuron. 2012;76(4):847–858. doi:10.1016/j.neuron.2012.09.015

101. Riddle J, McFerren A, Frohlich F. Causal role of cross-frequency coupling in distinct components of cognitive control. Prog Neurobiol. 2021;202:102033. doi:10.1016/j.pneurobio.2021.102033

102. Kleen JK, Testorf ME, Roberts DW, et al. Oscillation Phase Locking and Late ERP Components of Intracranial Hippocampal Recordings Correlate to Patient Performance in a Working Memory Task. Front Hum Neurosci. 2016;10. doi:10.3389/fnhum.2016.00287

103. Gaidica M, Hurst A, Cyr C, Leventhal DK. Interactions Between Motor Thalamic Field Potentials and Single-Unit Spiking Are Correlated With Behavior in Rats. Front Neural Circuits. 2020;14. doi:10.3389/fncir.2020.00052

104. Fleischer P, Abbasi A, Fealy AW, et al. Emergent Low-Frequency Activity in Cortico-Cerebellar Networks with Motor Skill Learning. eNeuro. 2023;10(2). doi:10.1523/ENEURO.0011-23.2023

105. Hall TM, de Carvalho F, Jackson A. A common structure underlies low-frequency cortical dynamics in movement, sleep, and sedation. Neuron. 2014;83(5):1185–1199. doi:10.1016/j.neuron.2014.07.022

106. Haberly LB, Shepherd GM. Current-density analysis of summed evoked potentials in opossum prepyriform cortex. J Neurophysiol. 1973;36(4):789–802. doi:10.1152/jn.1973.36.4.789

107. Rebert CS. Slow potential correlates of neuronal population responses in the cat’s lateral geniculate nucleus. Electroencephalogr Clin Neurophysiol. 1973;35(5):511–515. doi:10.1016/0013-4694(73)90027-8

108. Mitzdorf U. Current source-density method and application in cat cerebral cortex: investigation of evoked potentials and EEG phenomena. Physiol Rev. 1985;65(1):37–100. doi:10.1152/physrev.1985.65.1.37

109. Parker A, Derrington A, Blakemore C, Logothetis NK. The neural basis of the blood–oxygen–level–dependent functional magnetic resonance imaging signal. Philos Trans R Soc Lond B Biol Sci. 2002;357(1424):1003–1037. doi:10.1098/rstb.2002.1114

